# A chromosome-level genome for the nudibranch gastropod *Berghia stephanieae* helps parse clade-specific gene expression in novel and conserved phenotypes

**DOI:** 10.1101/2023.08.04.552006

**Authors:** Jessica A. Goodheart, Robin A. Rio, Neville F. Taraporevala, Rose A. Fiorenza, Seth R. Barnes, Kevin Morrill, Mark Allan C. Jacob, Carl Whitesel, Park Masterson, Grant O. Batzel, Hereroa T. Johnston, M. Desmond Ramirez, Paul S. Katz, Deirdre C. Lyons

## Abstract

How novel phenotypes originate from conserved genes, processes, and tissues remains a major question in biology. Research that sets out to answer this question often focuses on the conserved genes and processes involved, an approach that explicitly excludes the impact of genetic elements that may be classified as clade-specific, even though many of these genes are known to be important for many novel, or clade-restricted, phenotypes. This is especially true for understudied phyla such as mollusks, where limited genomic and functional biology resources for members of this phylum has long hindered assessments of genetic homology and function. To address this gap, we constructed a chromosome-level genome for the gastropod *Berghia stephanieae* (Valdés, 2005) to investigate the expression of clade-specific genes across both novel and conserved tissue types in this species. The final assembled and filtered *Berghia* genome is comparable to other high quality mollusk genomes in terms of size (1.05 Gb) and number of predicted genes (24,960 genes), and is highly contiguous. The proportion of upregulated, clade-specific genes varied across tissues, but with no clear trend between the proportion of clade-specific genes and the novelty of the tissue. However, more complex tissue like the brain had the highest total number of upregulated, clade-specific genes, though the ratio of upregulated clade-specific genes to the total number of upregulated genes was low. Our results, when combined with previous research on the impact of novel genes on phenotypic evolution, highlight the fact that the complexity of the novel tissue or behavior, the type of novelty, and the developmental timing of evolutionary modifications will all influence how novel and conserved genes interact to generate diversity.

## Background

One major question in biology is how novel phenotypes originate from conserved genes, processes, and tissues. Research in evolutionary developmental biology often focuses on the conserved modules that have been co-opted for new phenotypes, so called “toolkit” genes [1–3]. This approach has also been used to investigate the evolution of homologous adult phenotypes, such as in studies of sensory system evolution (e.g., G protein-coupled receptors [4]). However, a conservation-based approach explicitly excludes genetic elements that may be classified as clade-specific (i.e., taxonomically restricted, lineage-specific, lineage-restricted, or clade-restricted) that contribute to the development or function of a particular phenotype [1, 5–7]. Here we present the chromosome-level genome for the gastropod mollusk, *Berghia stephanieae* (Valdés, 2005) [8], which we used to identify clade-specific genes that may be important for both novel and conserved phenotypes, but have largely remained under investigated.

Many clade-specific genes are known to be involved in novel, or clade-restricted, phenotypes, including a number of cnidarian-specific genes exclusively expressed in specialized cell types called cnidocytes [9–11], the spiralian-specific gene trochin, expressed in the primary ciliated band [12], and spidroins in spiders, used for creating spider silk [13, 14], among others (further examples in [5]). A recent review by Wu & Lambert [5] highlighted that clade-specific genes deserve more attention when investigating evolutionary novelties.

In addition to their value in understanding phenotypic novelties, clade-restricted genes also play important roles in what might be considered more conserved phenotypes, and can quickly become essential to the viability of the organism (e.g., *Drosophila*, [15]). These clade-specific genes can become integrated into more conserved systems via some version of system drift (e.g., developmental system drift [16]), which may not result in a drastic change in function that we would classify as an evolutionary novelty. Most research on the impact of clade-specific genes has focused only on the presence or absence of clade-specific gene expression in novelties and how those genes have evolved [17–20], but has not provided comparisons with more conserved phenotypes in the same organism. A few studies have identified more clade-specific gene expression in novel tissues or cell types compared to those that are more conserved [21, 22], moving us closer to appreciating how clade-specific genes – and their expression – impact phenotypic evolution. Based on these previous studies, we can hypothesize that clade-specific genes are likely to be disproportionately upregulated in novel tissues. However, much of this research has centered on well-studied model systems, which limits our ability to generalize across other metazoan lineages.

Investigations into clade-specific genes in understudied groups or phenotypes have a high potential for generating exciting new hypotheses or expanding our technical creativity. Multiple excellent examples of this potential come from the phylum Mollusca, a clade containing taxa such as snails and slugs, cephalopods, bivalves, and chitons. Mollusca is the second most speciose metazoan phylum (after Arthropoda) and contains a great diversity of phenotypes that have already provided many useful insights, including cephalopods as alternative models to vertebrates for the evolution of complex brains and intelligence [23–25], bivalves as a means of understanding the nature of transmissible cancer [26], and gastropods for neuroscience research [27, 28] and as models of parasitism and immunity [29, 30]. However, a lack of genomes and functional biology resources for many members of this phylum has long hindered assessments of genetic homology and function [31]. This lack of resources has limited our ability to even identify clade-restricted genes in mollusk lineages, let alone characterize their expression or test their impact on phenotypes of interest. Luckily, some genomic resources, such as transcriptomes, are being sequenced at a much higher rate than whole genomes [32].

We propose that these resources can also be used to more accurately infer whether genes are clade-specific, so that we might further characterize their impact on the evolution of novel phenotypes.

In this paper, we present a chromosome-level genome for *Berghia* that we use to investigate clade-specific gene expression across multiple tissues. *Berghia stephanieae* (hereafter referred to as *Berghia*) is a species of gastropod in the order Nudibranchia, a clade of marine slugs that lose their shell during metamorphosis [33]. This species has been used as a model for the study of both more conserved systems, such as neurodevelopment [34] and reproductive development [35], as well as clade-restricted phenotypes such as the sequestration of cnidarian nematocysts [36, 37] and endosymbiosis [38, 39]. We combined an inferred proteome from *Berghia* with available genome and transcriptome data from other metazoan species – including mollusks such as cephalopods, bivalves, and other gastropods – to identify clade-specific *Berghia* genes (i.e., restricted to Mollusca, Gastropoda, Heterobranchia, Nudibranchia, Aeolidina, or *Berghia* alone). We then describe expression profiles of clade-specific and non-clade-specific genes among adult tissue samples in *Berghia,* including more ancient tissue types (like nervous system tissues; [40]) and clade-restricted ones (such as those associated with nematocyst sequestration; [41]). We find highly upregulated genes that are clade-specific in every tissue investigated, some of which are also highly upregulated in the same tissues during development. The proportion of clade-specific genes upregulated varied across tissues, but with no clear trend between the proportion of clade-specific genes and the novelty of the tissue. For example, the aeolidina-specific distal ceras did not express a significantly higher proportion of clade-specific genes compared to more “conserved” tissues, such as the foot or tail. Our results support previous assertions that clade-specific genes are important for our understanding of phenotypic evolution, and highlight that future studies in emerging model systems must account for these to truly describe how new phenotypes evolve.

## Results

### A high-quality *Berghia stephanieae* genome

We constructed a highly contiguous chromosome-level assembly of the 1.05 Gb long *Berghia stephanieae* genome using PacBio long read sequencing (16,476,079 total reads, 166,997 Gigabases, mean read length ∼10 kb, N50 = 15,977) that corresponds to ∼160x theoretical coverage based on the final assembly size. The assembly was scaffolded with Omni-C Illumina short read sequencing (Additional File 1). Post-assembly but pre-scaffolding, the assembly was 1.1 Gb with a contig N50 of 6,921,793 bp and L50 of 46 contigs (Additional File 1). Our scaffolded assembly contained 7945 scaffolds (Table 1) with an N50of 86 Mb (L50 = 5 scaffolds) and N90 of 34 Mb (L90 = 15). The rest of the scaffolds (7930 scaffolds) encompass less than 8% of the original genome assembly. A GenomeScope 2.0 [42] analysis of our Omni-C data indicated relatively low heterozygosity (0.693%) compared to many other mollusks, including gastropods such as *Elysia chloritica* (3.66% heterozygosity [43]), and many bivalves (1-3% [44–47]). The K-mer spectra and fitted models for *Berghia* are also consistent with a diploid genome (Additional File 2: Fig. S1). The final genome, filtered by GC content, BLASTn hits, and sequence length (see methods for details) contains 18 scaffolds (Table 1) with an N50 length of 85.5 Mb (L50 = 5 scaffolds) and N99 length of 26.7 Mb (L99 = 15 scaffolds). We found 93.3% complete, and 95.9% complete+fragmented, BUSCO core genes represented from the Metazoa (odb10) BUSCO database in the final dataset (Additional File 2: Fig. S2), and 76.6% of PacBio reads map to the final assembly (compared to 77.2% in the unfiltered assembly; Table 1). The 18 scaffolds in the final *Berghia* genome likely represent 15 chromosomes, based on the length distribution of the scaffolds and the linkage map (Fig. 2 A-B) from our Omni-C analyses. This is on the high end of the range for nudibranchs, which are known to have between 12 and 15 chromosomes in their haploid genomes [48]. All further analyses were performed on these 18 scaffolds.

**Figure 1.**
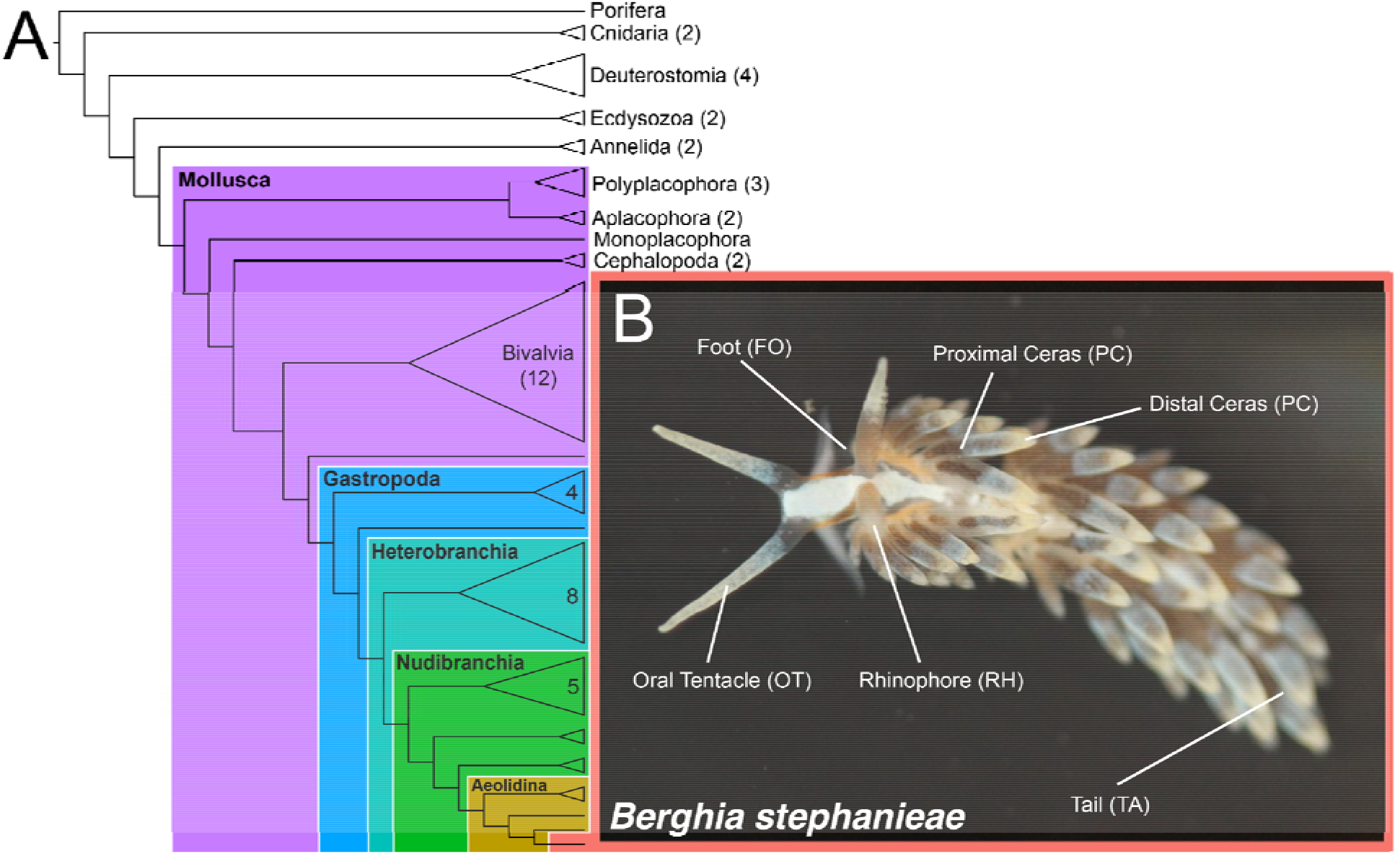
Cladogram showing broadly where the nudibranch *Berghia stephanieae* falls in the metazoan phylogeny. Colors indicate clades that we investigated for clade-specific genes in the *Berghia* genome: Mollusca (purple); Gastropoda (blue); Heterobranchia (teal); Nudibranchia (green); Aeolidina (gold); and (B) *Berghia stephanieae* (salmon). External tissues used in our analyses are indicated in (B). Major clades for outgroups included in our analyses are also shown, with the numbers in each collapsed clade or next to each name indicating the number of species from that group included in our analysis. Taxon details are available in Additional File 5: Table S3.

**Figure 2.**
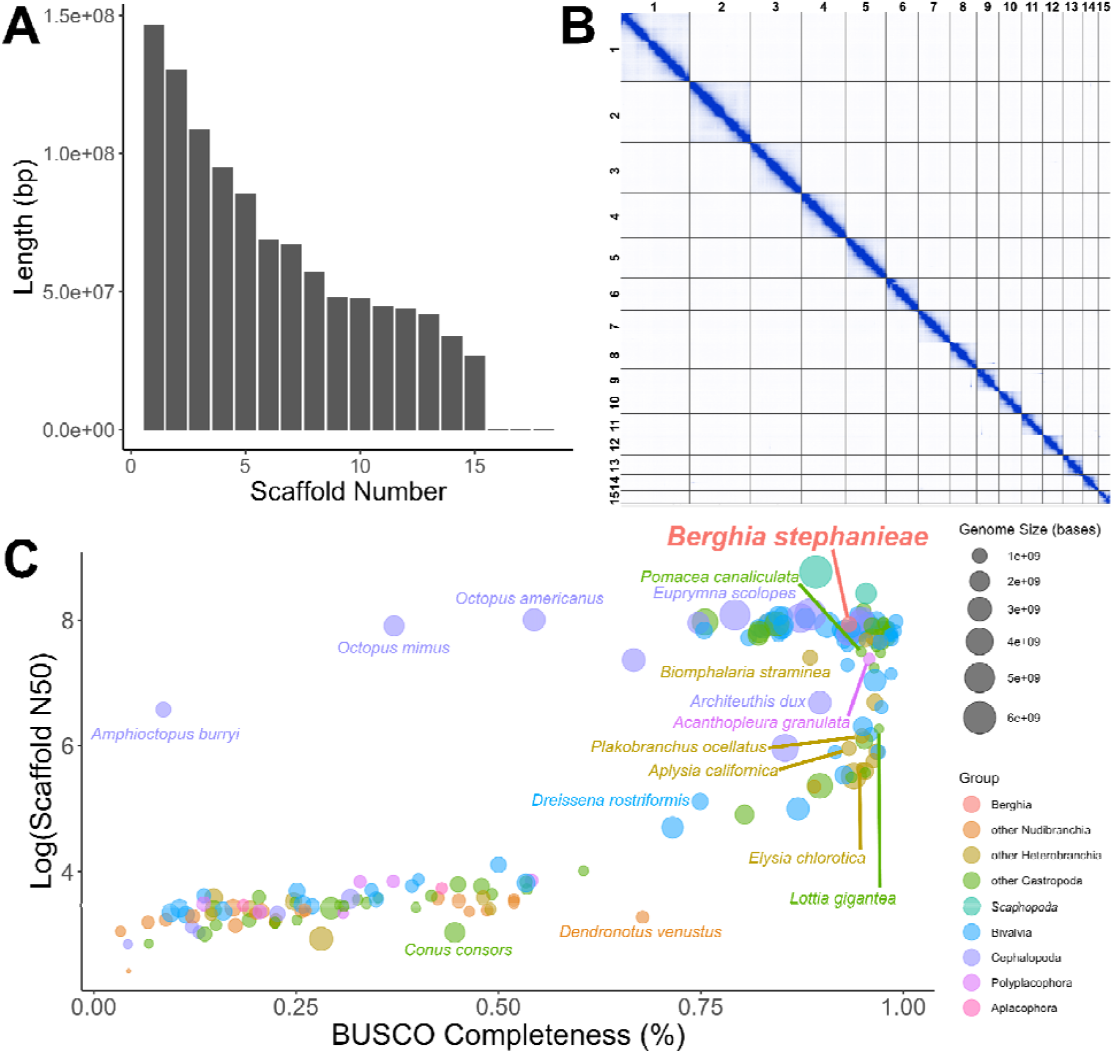
Summary statistics for the chromosome-level *Berghia stephanieae* genome, with comparisons to other mollusk genomes. (A) Bar chart showing the length (in bp) for each of the retained scaffolds, including 15 putative chromosomes. (B) Hi-C linkage plot showing identified links within and among chromosomes. Darker color of a block indicates higher frequency of contacts. (C) Plot visualizing the summary statistics for the *Berghia stephanieae* genome compared to assembled genomes of other mollusks by BUSCO completeness score (to the metazoa_odb10), log of the scaffold N50, and total genome assembly length. The *Berghia* genome is nearly average in size but is among the best genomes in terms of contiguity (scaffold N50) and BUSCO completeness score.

**Table 1.**
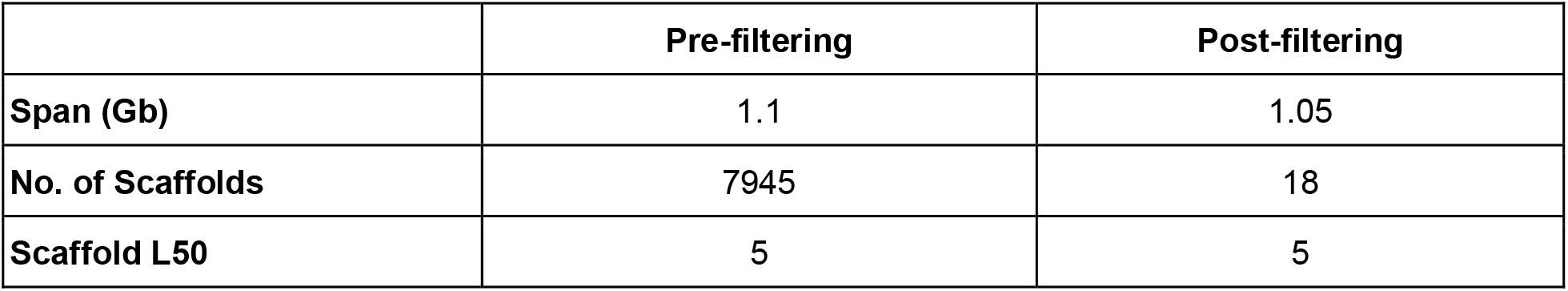
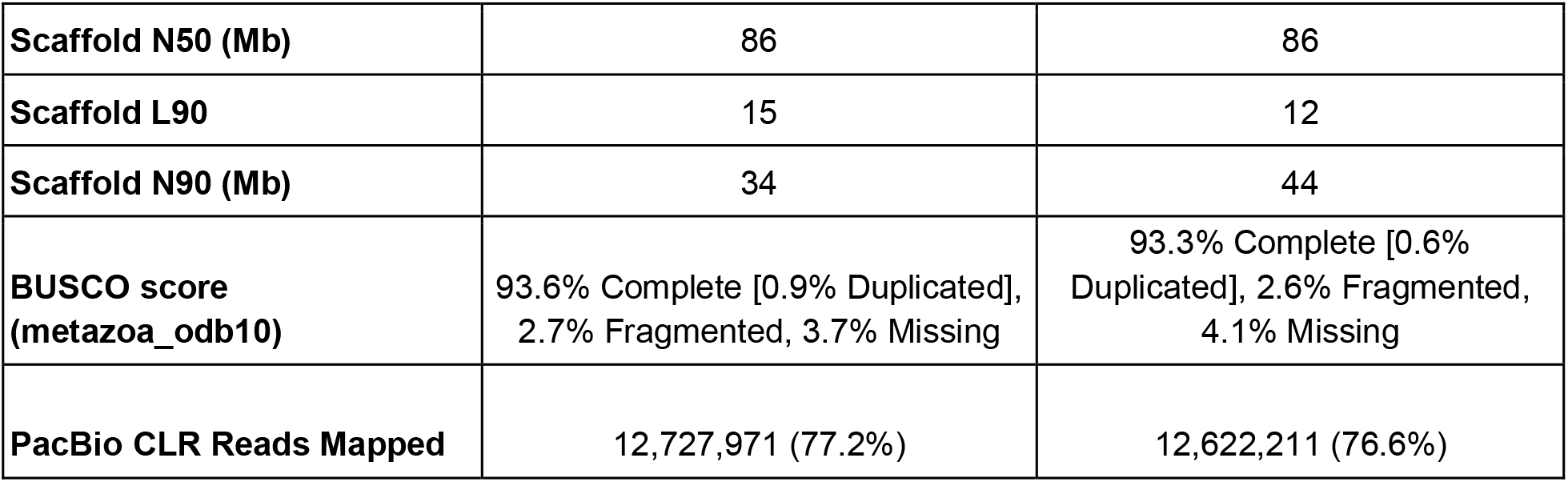
Genome assembly statistics for the *Berghia stephanieae* genome initial assembly (pre-filtering) and final assembly (post-filtering).

The *Berghia* genome compares favorably to other mollusk genomes in NCBI, with both a very high BUSCO score (when compared to the metazoa_odb10 database) and scaffold N50 (Fig. 2C; Additional File 3: Table S1). This analysis includes genomes from NCBI classified as either “scaffold”, “chromosome”, or “complete” from the phylum Mollusca. Of the 190 non-*Berghia* genomes analyzed, only 23 have a higher scaffold N50, the majority of which are larger genomes from cephalopods, bivalves, and scaphopods. Five are other gastropods, including the caenogastropods *Sinotaia purificata, Conus ventricosus* and *Monoplex corrugatus* and the patellogastropods *Patella vulgata* and *P. pellucida*, three of which have higher BUSCO scores.

Overall, 61 of the 190 species have higher BUSCO completeness scores, but the *Berghia* genome falls within a cluster of the most contiguous and highest quality genomes on NCBI (Fig. 2C).

The *Berghia stephanieae* genome is also well-annotated. Our RepeatModeler analysis identified 46.68% of the genome is repetitive elements, with the majority of bases characterized as unclassified repeats (27.45%, further details in Additional File 4: Table S2). BRAKER2 initially predicted 61,662 proteins (covering 59,494 genes; Table 2), which we subsequently filtered to a data set of 26,595 proteins and 24,960 genes for annotation and analysis using a script included with the BRAKER installation (selectSupportedSubsets.py) and the --anySupport flag to only include genes at least partially supported by hints. Prediction filtering resulted in a slightly lower BUSCO score (for both the Metazoa and Mollusca databases; Metazoa - 87.2% to 86.0% complete, Mollusca - 73.8% to 72.2% complete), though both scores were lower than the original BUSCO result using the whole genome, suggesting that gene predictions from BRAKER2 are incomplete. Prediction filtering also very slightly lowered IsoSeq mapping percentage (95.57% to 95.23% mapped reads), but did not change the percentage of short reads mapped to gene models (74.81% for both). Functional prediction rates, however, were much improved in our filtered data set, for both BLASTP hits (20,820 proteins, 78.3%, in filtered predictions) and InterProScan results (24,469 proteins, 92.0%, in filtered predictions) compared to 58.8% and 78.7%, respectively, in our initial predictions (Table 2). We used our filtered BRAKER2 predictions and functional annotations in subsequent analyses.

**Table 2.**
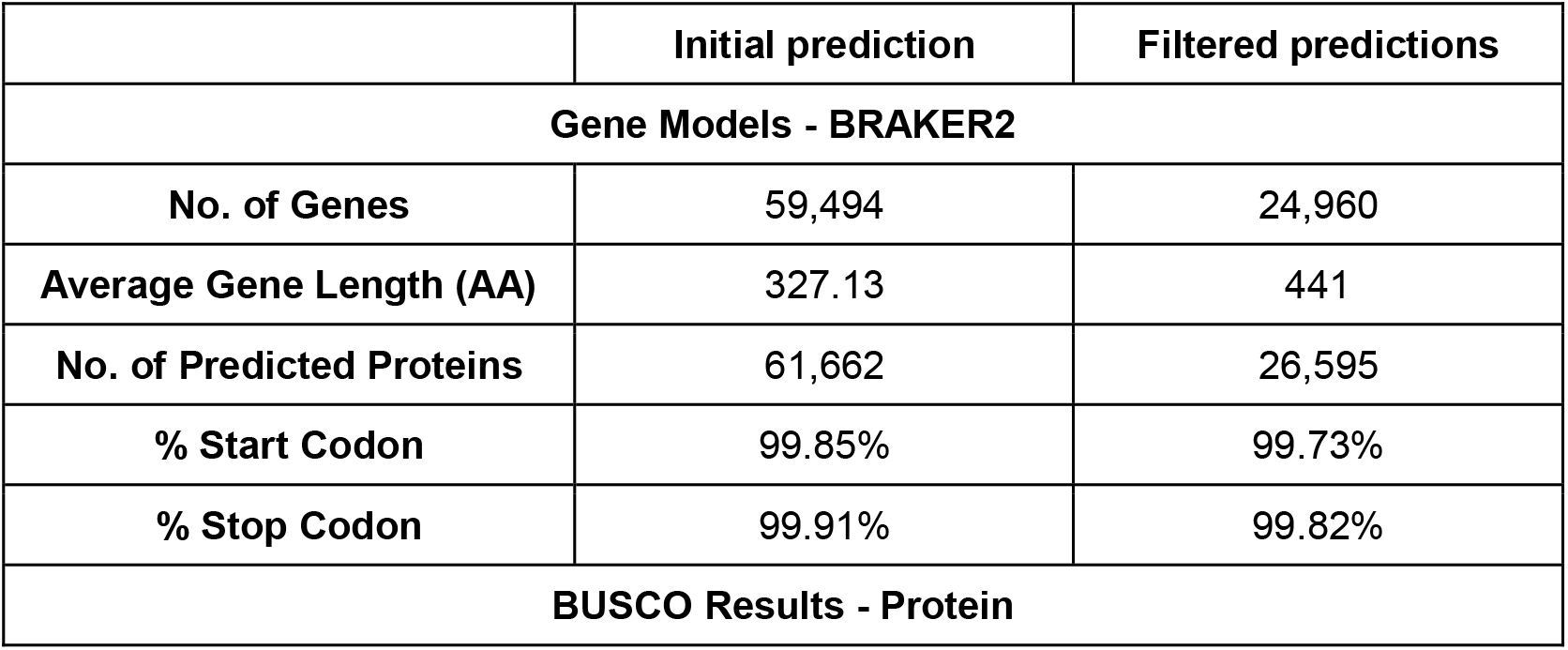
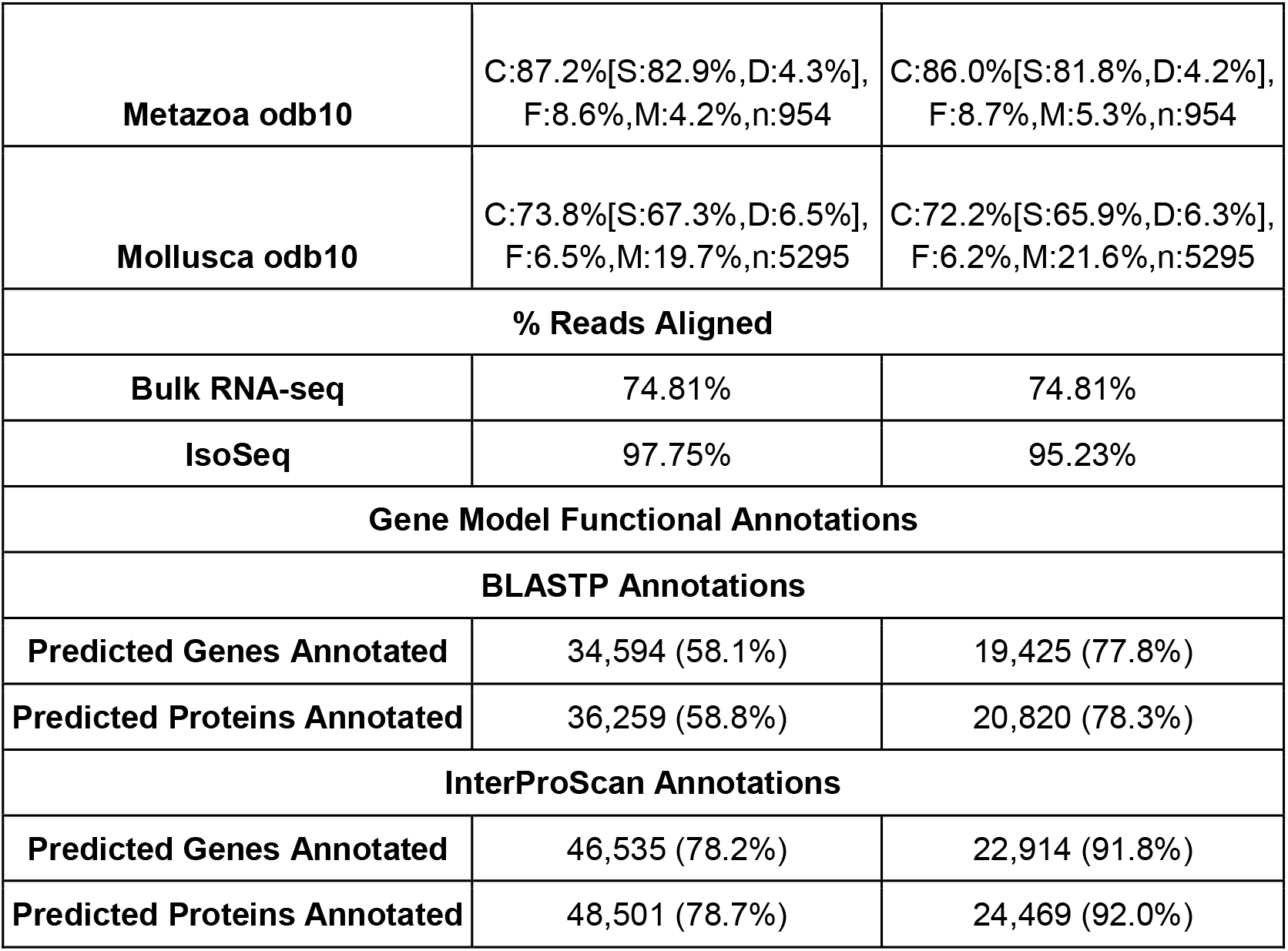
Gene prediction and annotation statistics for the *Berghia stephanieae* genome, including initial gene models predicted from BRAKER (pre-filtering) and filtered gene models intended to include only those models with external support (post-filtering).

### Identification of clade-specific genes in *Berghia* genome

Our OrthoFinder analysis compared the predicted *Berghia stephanieae* proteins from BRAKER2 with proteomes from 58 other metazoan species (Additional file 5: Table S3), including 27 gastropods, one scaphopod, twelve bivalves, two cephalopods, one monoplacophoran, 3 polyplacophorans, two aplacophorans and eleven non-molluscan species. The goal of this analysis was to generate orthologous groups among proteins from all proteomes to assess which *Berghia* genes are restricted to certain clades. It is important to note that genes classified as restricted to narrower taxonomic designations (e.g., Gastropoda) are also by definition restricted to higher taxonomic clades (e.g., Mollusca). We found 25,338 (95.2%) *Berghia stephanieae* proteins clustered into orthogroups, 1,027 (3.9%) of which were in *Berghia*-specific clusters of two or more sequences.

Our KinFin analysis, which provides taxon-aware annotation of inferred orthologous groups, identified *Berghia* genes restricted to Mollusca (n = 1,067, 4.3% of genes), Gastropoda (n = 463, 1.9% of genes), Heterobranchia (n = 1,154, 4.6% of genes), Nudibranchia (n = 1,030, 4.1% of genes), Aeolidina (n = 108, 0.4% of genes), as well as those genes only found in *Berghia* (n = 2,188, 8.8% of genes; Fig. 3A). These *Berghia-*specific genes include those clustered in *Berghia-* specific clusters using OrthoFinder in addition to singletons that were not clustered. This level of species-specificity is on the lower end compared to other nudibranchs and fairly average for mollusks more broadly (Additional file 2: Figs. S3-S4) This means that 18,957 *Berghia* genes (75.9% of genes) are not clade specific at the levels we investigated. Rarefaction curves for each taxonomic level (Additional file 2: Figs. S5-S9) suggest sufficient sampling in all major clades investigated. Both clade-specific and non-clade-specific (Other) genes are well distributed across the genome (Fig. 3C), with higher numbers of both types of genes per Mb in chromosomes 3 and 9 compared to other chromosomes (Additional file 2: Fig. S10). The proportion of clade-specific genes appears to be enriched on smaller chromosomes (Additional file 2: Fig. S11).

**Figure 3.**
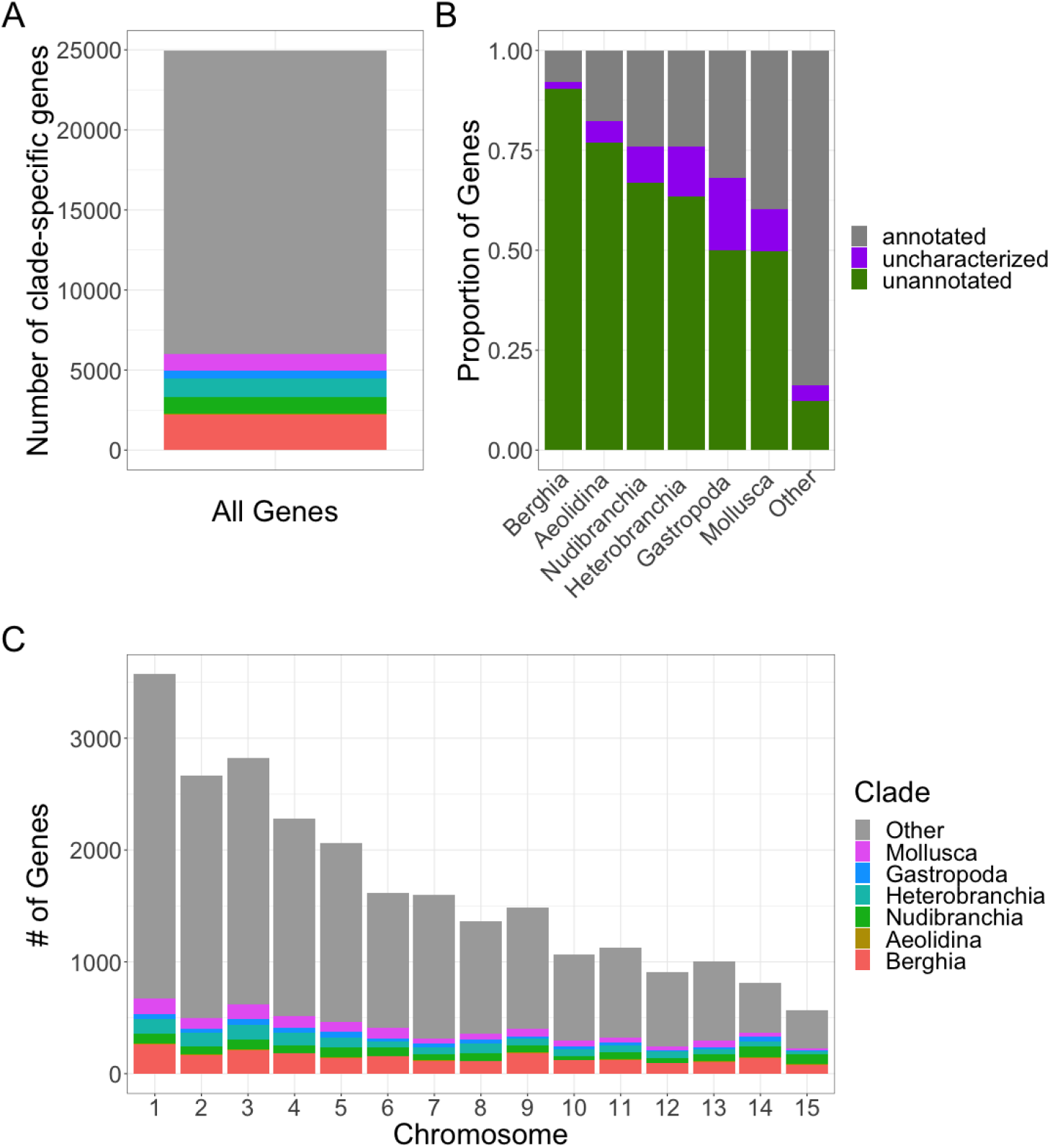
Results for clade-specific genes found within *Berghia stephanieae*. (A) A bar chart showing the total number of genes in *Berghia stephanieae* parsed by whether they fall into one of the clade-specific groupings or not (Other); (B) A bar chart indicating what proportion of genes within each group (clade-specific or Other) were annotated using BLASTP. In some cases, BLASTP found a match to an uncharacterized, hypothetical, or putative protein. These are separated into a different category (uncharacterized). (C) A bar chart showing the distribution of genes across the genome, parsed by whether they are clade-specific or not (a size normalized version of this chart is available in Additional file 5: Fig. S5)

As expected, non-clade specific genes (Other, Fig. 3B) were much more likely to be annotated via UniProt (79.9%) than genes identified as clade-specific. As we get phylogenetically deeper, the proportion of annotated genes increases: *Berghia*-specific (7.8%), Aeolidina-specific (17.5%), Nudibranchia-specific (24.2%), Heterobranchia-specific (24.2%), Gastropoda-specific (32.0%), and Mollusca-specific (39.8%). The proportion of matched but uncharacterized genes (including hypothetical proteins) ranged from 1.2% (Heterobranchia-specific) to 18.1% (Gastropoda-specific) across investigated clades.

### Clade-specific gene expression across tissues

#### Bulk tissue RNA-seq data mapped to the genome

We included seven *Berghia* tissues in our differential expression analyses, including: (1) the brain, which consists of the paired cerebral-pleural, pedal, buccal, and rhinophore ganglia, (2) rhinophores, which are chemosensory structures restricted to nudibranchs, (3) oral tentacles, which are sensory-motor appendages restricted to the clade Aeolidina, (4) distal cerata, which are also restricted to the nudibranch clade Aeolidina and contain the organ where nematocyst sequestration occurs, (5) proximal cerata, which are common in Aeolidina and a few other nudibranchs and contain branches of the digestive system, and (6) foot, and (7) tail, which are tissues associated with mollusks more broadly. Our RNA-seq samples ranged from 2.9 million (SRR14337001, brain) to 36.5 million (SRR12072210, oral tentacle) read pairs (xLJ = 25.3 ± 8.6 million reads per sample; Additional File 6: Table S4). On average, 72.1% (± 7.8%) of read pairs mapped uniquely to the *Berghia stephanieae* genome, and mapping percentage ranged from 53.4% (SRR14337002, brain) to 80.6% (SRR12072207, distal ceras). We identified expression (counts >10 across all tissues [49]) in ∼97.0% of genes (24,178 out of 24,960 predicted genes).

#### Differential expression among *Berghia* tissues

Our differential expression analyses compared expression of each gene in each tissue to an average of normalized expression of that gene across all other tissues. Genes with a Log2 Fold Change > 2 and adjusted p-value < 0.05 were considered upregulated in a given tissue. We identified 16,691 genes upregulated across all tissues, with the highest number of upregulated genes (Table 3; Additional File 7: Table S5) in brain tissue (15,210 genes), followed by proximal ceras (678 genes), foot (205 genes), distal ceras (188 genes), tail (169 genes), rhinophore (147 genes), and oral tentacle (94 genes), respectively. The proportion of upregulated genes that were clade-specific (Fig. 4A; Additional File 7: Table S6) was variable across tissues, with rhinophore, oral tentacle, distal ceras, and tail having the most similar proportions of clade-specific upregulated genes (28.8–34.4% of upregulated genes), followed by foot (21.0% of upregulated genes). The proportion of clade-specific upregulated genes (Fig. 4A) was much lower in brain tissue (12.0% of upregulated genes). Expression data from clade-specific genes with more recent homologs (i.e., Gastropoda-*Berghia*) were more useful for distinguishing all tissues except for the brain (Additional File: Figures S9-S14). Some genes upregulated in one tissue were also upregulated in another tissue (1,779 genes, or 10.7% of upregulated genes; Fig. 4C).

**Figure 4.**
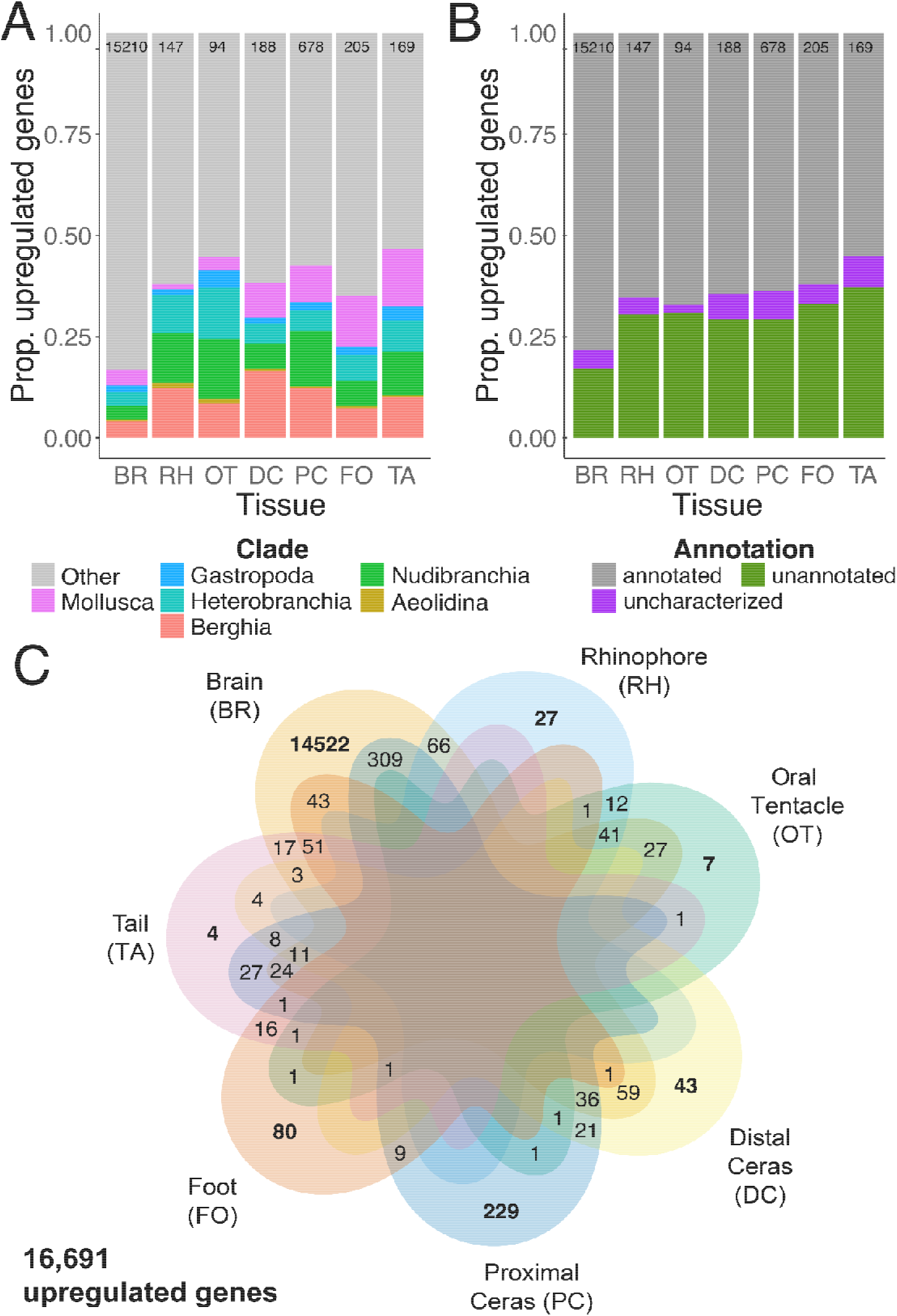
Results for upregulated genes for each tissue found within *Berghia stephanieae*. (A) A bar chart showing the proportion of upregulated genes that are clade-specific, or not (Other), for each tissue type. (B) A bar chart indicating what proportion of genes upregulated for each tissue type were annotated using BLASTP. In some cases, BLASTP found a match to an uncharacterized, hypothetical, or putative protein. (C) Venn diagram showing the tissues in which all upregulated genes were determined to be upregulated. Some genes are only upregulated in certain tissues, while others are upregulated in multiple tissues (maximum of 4 out of 6, 11 genes upregulated among the brain, distal ceras, proximal ceras, and tail). These are separated into a different category (uncharacterized). Abbreviations: BR, brain; DC, distal ceras; FO, foot; OT, oral tentacle; PC, proximal ceras; RH, rhinophore; TA, tail.

**Table 3.** Number upregulated genes for each tissue in *Berghia stephanieae*, by clade-specificity designation. Total numbers of upregulated genes are also reported for each tissue and clade.

|  | Brain | Rhinophore | Oral Tentacle | Distal Ceras | Proximal Ceras | Foot | Tail | Total |
| --- | --- | --- | --- | --- | --- | --- | --- | --- |
| **Berghia** | 623 | 18 | 8 | 31 | 83 | 15 | 17 | 795 |
| **Aeolidina** | 52 | 2 | 1 | 1 | 3 | 1 | 1 | 61 |
| **Nudibranchia** | 520 | 18 | 14 | 12 | 93 | 13 | 18 | 688 |
| **Heterobranchia** | 536 | 14 | 12 | 9 | 35 | 13 | 13 | 632 |
| **Gastropoda** | 238 | 2 | 4 | 3 | 13 | 4 | 6 | 270 |
| **Mollusca** | 593 | 2 | 3 | 16 | 62 | 26 | 24 | 726 |
| **Other** | 12648 | 91 | 52 | 116 | 389 | 133 | 90 | 13519 |
| **Total**<br>**upregulated** | 15210 | 147 | 94 | 188 | 678 | 205 | 169 | 16691 |

We then removed genes that were upregulated in multiple tissues and focused only on those genes upregulated in a single tissue (uniquely upregulated genes; 14,912 genes, or 89.3% of upregulated genes). The highest number of uniquely upregulated genes (Fig. 4C was still in brain tissue (14,522 genes), followed by proximal ceras (229 genes), foot (80 genes), distal ceras (43 genes), rhinophore (27 genes), oral tentacle (7 genes), and tail (4 genes). The clade-specificity and annotation distributions of uniquely upregulated genes differed in some tissues (Fig. 5; Additional File 8: Tables S7-S8), including the rhinophore, oral tentacle, and tail when compared to the distributions of all upregulated genes in those tissues. The proportions of clade-specific and annotated genes changed more significantly in those tissues where far fewer genes were uniquely upregulated (Fig. 5).

**Figure 5.**
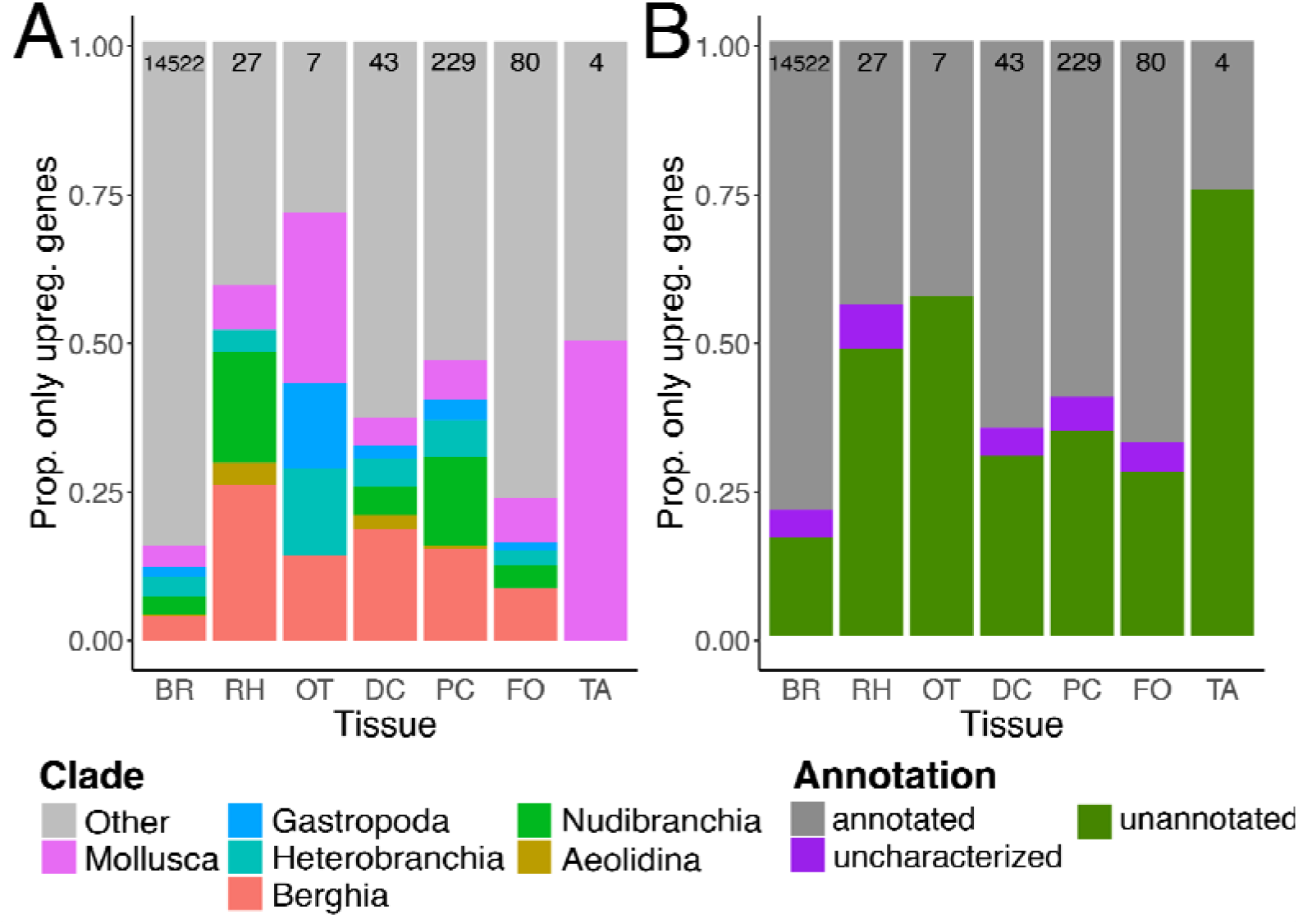
Results for genes only found upregulated in a single tissue of *Berghia stephanieae* (see **bold** values in Fig. 4C). (A) A bar chart showing the proportion of upregulated genes that are clade-specific, or not (Other), for each tissue type. (B) A bar chart indicating what proportion of genes upregulated for each tissue type were annotated using BLASTP. In some cases, BLASTP found a match to an uncharacterized, hypothetical, or putative protein. Abbreviations: BR, brain; DC, distal ceras; FO, foot; OT, oral tentacle; PC, proximal ceras; RH, rhinophore; TA, tail.

With regard to annotation, we noted that in addition to having the highest number of upregulated genes, brain tissue also had the highest proportion of upregulated genes (78.3%) that were annotated via BLASTP (Fig. 5B; Additional File 9: Table S9). The other tissues had slightly lower levels of annotation ranging from 55.0% (tail) to 67.0% (oral tentacle) with multiples included, though for some tissues this proportion of upregulated, annotated genes dropped significantly when considering genes upregulated in only one tissue (Fig. 5B). Of the upregulated genes with annotations (Fig. 4B), GO term enrichment analyses were consistent with what might be expected for particular tissues (Additional File 10: Tables S10-S16). For example, we noted enrichment of signal transduction, transmembrane transport, and ion binding GO terms in the brain, G protein-coupled receptor activity and sodium ion transport terms in the sensory oral tentacles and rhinophores, and transmembrane transporter activity, transmembrane transport, and extracellular region terms in the distal ceras, where nematocyst sequestration occurs.

#### Confirmation of tissue-restricted expression of clade-specific genes

In order to confirm the expression of clade-specific genes inferred to be upregulated in certain tissues, we localized expression of at least one gene from each of two tissues (rhinophores and distal ceras) using *in situ* Hybridization Chain Reaction (HCR) techniques [50] in *Berghia* juveniles. We also compared our list of genes upregulated in the *Berghia* brain to HCR gene expression profiling in the brain of adult *Berghia* available from Ramirez et al. [51] Juveniles were selected for our experiments because adult *Berghia* contain pigments that make localizing expression more difficult, and both the distal ceras and rhinophores are identifiable and functional at an early juvenile stage [34, 36]. We localized a distal ceras upregulated Nudibranchia-specific gene (jg13556; annotated as collagen alpha-1, match to UniProt ID Q7LZR2, e-value = 0.000129) in juvenile distal cerata (Fig. 6A-A’”). This gene appears to be exclusively expressed inside the cnidosac of the juvenile *Berghia*, where nematocyst sequestration is known to occur [36]. We also identified expression of a rhinophore upregulated *Berghia*-specific gene (small domain annotated as a Pancreatic trypsin inhibitor, match to UniProt ID P00974, e-value = 1.52E-07) in the juvenile rhinophores where it is expressed in patches on the external epithelium (Fig. 6B-B”’). For the brain, we found numerous clade specific genes are expressed in the *Berghia* brain based on single cell RNA-seq data [51]. These include: (1) two genes exclusively upregulated in the brain (jg44129, an unannotated *Berghia*- specific gene expressed in a cluster of glial cells, and jg54950, an unannotated Heterobranchia- specific gene upregulated in nitric oxide synthase (*Nos*) and pigment dispersing factor (*Pdf*)- expressing cells in the rhinophore ganglia (*rhg*)), (2) one gene upregulated in the brain, rhinophore, and oral tentacle in our analysis (jg22847, an unannotated *Berghia*-specific gene in *Nos/Pdf- rhg* cells, and (3) two genes not considered upregulated in the brain in our analysis but appear to be upregulated in certain cell clusters in the brain (jg57406, an unannotated Heterobranchia-specific gene that is upregulated in the distal ceras in our analysis but expressed in mature neurons in all *Berghia* ganglia; jg56194, an unannotated Nudibranchia-specific gene that is not upregulated in any tissue in our analysis but is found in a cluster of glial cells in the brain) [51].

**Figure 6.**
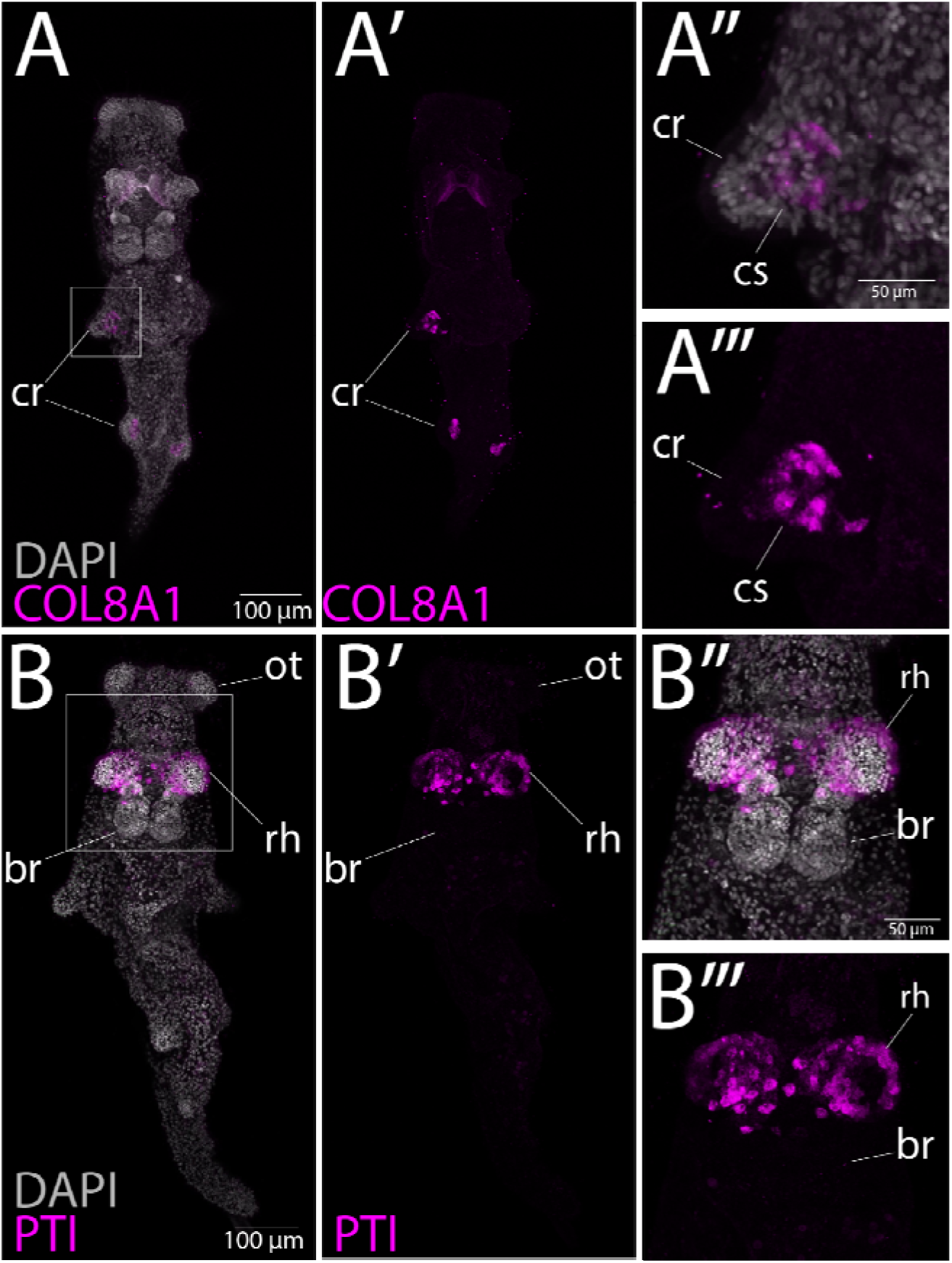
HCR results in *Berghia stephanieae* juveniles for selected clade-specific genes upregulated in particular adult tissues. (A-A’’’) Juveniles stained for a collagen alpha 1 VIII gene found upregulated in the distal ceras (COL8A1; jg13556) (A-A’) DAPI and Alexa 647 stained tissues in whole animal, and (A’’-A’’’) close up of cnidosac. (B-B’’’) Juveniles stained for a gene found upregulated in the rhinophores (annotated as a Pancreatic trypsin inhibitor, PTI; jg18351) (B-B’) DAPI and Alexa 647 stained tissues in whole animal, and (B’’-B’’’) close up of rhinophore. Abbreviated: cr, ceratas; cs, cnidosac; br, brain; rh, rhinophore; ot, oral tentacle.

## Discussion

### The *Berghia stephanieae* genome is highly contiguous

The *Berghia stephanieae* genome is among the most contiguous and highest quality gastropod genomes to date (Fig. 2C). The final assembled and filtered *Berghia* genome is comparable to other mollusk genomes [52] in terms of size (1.05 Gb) and number of predicted genes (24,960 genes). The *Berghia* genome also has high Metazoa, and moderate Mollusca, BUSCO scores (Table 2), both comparable to the scores of other high-quality mollusk genomes [53, 54]. Our analysis also identified a high percentage of repetitive elements in the *Berghia* genome (46.68%), similar to rates found in other mollusk species [32]. However, given that only 76.6% of PacBio CLR reads mapped to the final genome assembly (Table 1), it is likely that many repeat regions remain unresolved. The proportion of annotated genes in the *Berghia* genome was also quite high (77.8% with BLASTP hits), consistent with other published gastropod genomes [55–57].

### Clade-specific genes are a small percentage of the *Berghia* genome

The vast majority of predicted genes in the *Berghia* genome (18,957 genes, 75.9%; Fig. 3A) are not restricted to the clades of interest in our analysis (Mollusca, Gastropoda, Heterobranchia, Nudibranchia, Aeolidina, and *Berghia*). Of those identified as clade-specific, most (2,188 genes, ∼8.8%) were classified as *Berghia*-specific genes. This percentage of “orphan” or species-specific genes in *Berghia* is on the lower end of the range compared to other species from across Metazoa (∼1 to >30%; [7]), which may simply be a feature of the *Berghia* genome. This may also be because our strategy to perform orthologous gene inference with a clustering-based method using both unannotated transcriptome data and well annotated, high-quality genome data increased our chances of detecting homologs to *Berghia* genes. Clustering-based algorithmic approaches for inferring gene orthology apply normalization to pairwise similarity scores to account for sequence length of the query and length of its hits, ensuring that distantly related sequences receive comparable scores compared to the best-scoring sequences from closely related species [58–60]. This strategy, combined with more data from more closely related taxa, provides greater opportunity for similarity matches and limits the impacts of homology detection failure [61], which occurs when homologs have become undetectable by search algorithms even though they exist. Our analysis suggests that homology detection failure is likely to cause inflated estimates of the proportion of species-specific genes in analyses that: (1) include species from less widely distributed taxonomic levels (e.g., only including closely related taxa [19] or having limited outgroup sampling [62]); (2) rely only on pairwise similarity scores for assessing homology [21, 62]. In our analysis, this could mean that a subset of the ∼8.8% of *Berghia-*specific genes are likely to be restricted to clades that we have not explicitly investigated, such as those at the genus (*Berghia*) or family (Aeolidiidae) levels. This would mean that the actual percentage of species-specific *Berghia* genes is perhaps even lower than reported here.

As might be expected, we also note that the annotation rate goes down with an increase in clade specificity of the genes, meaning that non-clade-specific genes (“Other”) had the highest annotation rate and *Berghia*-specific genes the lowest (Fig. 3B). However, some *Berghia* genes matched to proteins in the NCBI RefSeq database that do not have functional data associated with them (i.e., are predicted, hypothetical, or unidentified proteins; purple color in Fig. 3B). It is possible that some of these putative annotations are due to protein domain-level similarities, which suggests that these clade-specific genes do share some homologous regions with genes in other taxonomic groups.

### Novel tissues do not express more novel genes

Clade-specific genes have long been thought to be a driver of morphological novelties [5, 63], and some researchers have hypothesized that novel phenotypes might require a higher frequency of clade-specific genes [22, 63, 64]. This type of increase in the expression of clade-specific genes has been identified in some taxa. These include cnidarians like *Nematostella* in novel cells called nematosomes [22], the mollusk radula [65], and in non-morphological novelties like eusocial evolution in honey bees [64]. Our results are inconsistent with this hypothesis. In *Berghia,* we did not see a clear increase in the proportion of upregulated clade-specific genes in morphological novelties (Fig. 4A-B). For example, the distal ceras, where nematocyst sequestration occurs inside a novel organ called the cnidosac [41, 66], appears to have an average level of clade-specific gene upregulation compared to other tissues (Fig. 4A- A’’’). This may be due to the fact that nematocyst sequestration relies on a largely conserved process, namely phagocytosis [36, 41, 67]. Our results also did not show a higher proportion clade-specific genes upregulated in the sensory rhinophores or oral tentacles compared to other tissues, which are functionally similar and homologous to tentacles in other heterobranch gastropods and caenogastropods [34, 68, 69]. However, the clade-specific genes that are upregulated in some of these tissues do appear to be functionally important, as our HCR results indicate that some genes (as shown in Fig. 6) are not only upregulated in the adult tissues but are also highly expressed in those same tissues in early stage juveniles. The expression of these genes in juveniles suggests that these clade-specific genes may be especially crucial for the core function of these tissues as soon as they form. Alternatively, these genes may be important for upstream developmental processes in addition to downstream functions.

So why are our results inconsistent with the hypothesis that clade-specific genes are likely to be upregulated in novel tissues? For one, we have only collected gene expression data under standard conditions, which means that we are largely capturing constitutive gene expression. It is possible that critical, clade-specific genes for certain tissues may be only expressed in response to certain stimuli. We have likely not captured many of these genes in our analyses.

Second, it is known that clade-specific genes are involved in both novel and conserved phenotypes [15, 21]. However, investigating gene expression at the level of tissues likely masks hidden differences in cell type diversity and complexity within tissues. We hypothesize that differences in the number and expression levels of clade-specific genes among novel tissues may be more related to the type of novelty (i.e., functional vs morphological) rather than the level of biological organization. For example, novelties that arise from a loss or modification of function may only require the loss of expression of certain genes. Future investigations focused on the expression of clade-specific genes (facultative and constitutive) at the cellular level – and on the function of these genes – will provide the necessary data to assess the relative impacts of these possibilities.

### Tissue complexity - the *Berghia* brain

Animal nervous systems are perhaps the most complex biological system, with their diverse components, cell types, and functions (e.g., [70, 71]). Although neurons in general express more genes than many other cell types in a variety of metazoan lineages [72], it has also been noted that genetic novelties appear to underlie cell diversification in multiple lineages (including octopuses [25] and teleost fish [73]). In these cases, the proportion of clade-specific genes expressed in novel cell types is higher than cell types with clear homologs. However, these differences would not be detectable in tissue-level analyses, such as those presented here.

Despite lower read counts and mapping percentages (Additional File 6: Table S4), we identified a large number of genes upregulated in *Berghia* brain tissue (>15,000; Fig. 4A), which is roughly two-thirds of predicted *Berghia* genes. This is consistent with the hypothesis that neurons express more genes than other cell types [72]. However, our results also show lower proportions of clade-specific genes in the brain (∼16.8%; Fig. 4A) compared to other tissues, even though the number of clade-specific genes upregulated in the brain (2,562 genes) is higher overall than the total number of upregulated genes in any other tissue (94–678 genes; Fig. 4B). It is possible then that consideration of clade-specific expression at the tissue level may not provide a comprehensive understanding of novelty.

Although some prior studies identified expression of more clade-specific genes in novel tissues or structures [21], others have found a higher frequency of clade-specific expression in cell type novelties [22, 74]. This suggests that the expression of clade-specific genes may have a higher impact on the evolution of novel cell types rather than novel tissues as a whole. This hypothesis is supported by the expression patterns of clade-specific genes in the *Berghia* brain [51]. For example, an unannotated gene that serves as a neuron-specific cell type marker is also a Heterobranchia-specific gene (jg57406). Similarly, serotonergic neurons in the brain express an unannotated gene that is Nudibranchia-specific (jg38442) [51].

### Type of novelty - the *Berghia* cnidosac

The novel-genes-drive-innovation hypothesis might suggest that the novel cnidophage cell type, in the distal ceras [36], may use more (or a higher proportion of) clade-specific genes than other, more conserved cell types in *Berghia*. However, it was previously hypothesized that the cnidophage cell type may simply be a more specialized homolog of a digestive cell type, due to the apparent endodermal nature of cnidophages [36, 41, 66, 75]. A logical inference from this hypothesis is that the novel function of sequestration in cnidophages does not necessarily require novel molecular processes. This is not a new idea. For example, some morphological novelties in plants have been shown to evolve via regulatory evolution [76] and social behavior evolution in ants has been tied to both conserved and novel genetic elements [77]. Under similar conditions, we might not expect an increase in the frequency of clade-specific genes expressed in cnidophages. However, these clade-specific genes may still be functionally important given their high levels of expression in the cnidosac in *Berghia* juveniles (Fig. 6B-B”).

## Conclusions

The *Berghia stephanieae* genome is the first high quality published genome for the order Nudibranchia, and is one of the most contiguous and highest quality gastropod genomes to date. However, it is likely that many repeat regions remain somewhat unresolved. We used this genome to investigate how clade-specific gene expression is distributed across functionally and evolutionarily diverse tissue types in adult *Berghia*, and showed that upregulated genes in novel tissue types are not necessarily more likely to be classified as clade-specific. The proportion of clade-specific genes upregulated varied across tissues, with novel tissues like the distal ceras unexpectedly expressing a fairly average frequency of clade-specific genes compared to other tissues. Our results, when combined with previous research on the impact of novel genes on phenotypic evolution, highlight the value of a more holistic approach to investigating how phenotypes arise and diversify. In particular, the complexity of the novel tissue or behavior, type of novelty [22], and where across development changes may have occurred [78] will all influence how novel and conserved interact to generate new phenotypes.

## Methods

### Sample preparation and genome sequencing

We isolated one *Berghia* juvenile from the Lyons lab culture prior to mating to minimize genomic contamination. While isolated, we fed the animal ∼½ of a medium *Exaiptasia diaphana* (defined by Taraporevala et al. [35]) each day for 34 days. We then starved the animal for 44 days prior to shipping. To minimize residual food in the gut diverticula, cerata were removed with forceps and the remaining body was blotted on a kimwipe to remove excess water, then the animal was placed in a cryotube and flash frozen in liquid nitrogen and stored at -80 until shipping to Dovetail Genomics (now Catana Bio, Scotts Valley, CA). Dovetail Genomics used an input of ∼101mg into a slow CTAB protocol to extract high molecular weight DNA. They measured the efficiency of DNA extraction using a Qubit 2.0 Fluorometer (Life Technologies, Carlsbad, CA, USA) High Sensitivity Kit. Overall, they obtained 12.1 ug of high molecular weight DNA. They then used a Mini Column for clean up and resuspended the pellet in 75 µl TE. They then quantified DNA samples using the Qubit. They constructed the PacBio SMRTbell library (∼20kb) for PacBio Sequel using SMRTbell Express Template Prep Kit V 2.0 (PacBio, Menlo Park, CA, USA) using the manufacturer recommended protocol. They then bound the library to polymerase using the Sequel II Binding Kit 2.0 (PacBio) and loaded onto PacBio Sequel II (PacBio) on 8M SMRT cells (SRR25687008).

For scaffolding, Dovetail fixed chromatin in place with formaldehyde in the nucleus for extraction and analysis via Dovetail® Omni-C® proximity ligation. They then digested the fixed chromatin with DNAse I, repaired the chromatin ends and ligated to a biotinylated bridge adapter followed by proximity ligation of adapter containing ends. After proximity ligation, they reversed crosslinks and purified the DNA. They treated purified DNA to remove biotin that was not internal to ligated fragments. They generated sequencing libraries using NEBNext Ultra enzymes and Illumina-compatible adapters. They then isolated biotin-containing fragments using streptavidin beads before PCR enrichment of each library. Technicians then sequenced the library on an Illumina HiSeqX platform to produce approximately 30x sequence coverage. They then used HiRise MQ>50 reads for scaffolding (see “read-pair” above for figures).

### Short read RNA sample collection and sequencing

We obtained *Berghia* adult tissue samples, including the: (1) brain, (2 samples; SRR14337001- SRR14337002) (2) oral tentacles (3 samples; SRR12072210, SRR25598600-SRR25598601), (3) rhinophores (3 samples; SRR12072209, SRR25598592-SRR25598593), (4) foot (2 samples; SRR12072206, SRR25598598), (5) tail (2 samples; SRR12072205, SRR25598599), and (6) proximal (3 samples; SRR12072208, SRR25598594-SRR25598595) and (7) distal ceras (3 samples; SRR12072207, SRR25598596-SRR25598597). We also obtained earlier stage transcriptome data from: (8) multiple embryonic stages (bulk sample of 500-600 individual embryos from each time point reared at 27°C (12, 24, 36, 48, 60 hpo and 4, 7, and 9 dpo; SRR12072213) and (9) juveniles 15 dpo at 27°C (500 individuals from 3 egg masses laid the same day; SRR12072212). We starved adults for ∼1 week prior to removal of some tissues (all but the brain) to reduce symbiont presence and minimize contamination. We extracted total RNA from most adult tissues (minus the brain) using the RNeasy Kit (QIAGEN, Redwood City, CA) and submitted the extracted total RNA to Novogene Ltd. (Sacramento, CA) for quality assessment, library preparation and sequencing (Illumina NovaSeq 6000; 150bp paired-end reads). We prepared the adult brain total RNA using the Clontech SmartSeq v4 Ultra-Low Input RNA Kit (Takara). We prepared libraries with the Nextera XT DNA Library Preparation Kit and 96-Sample Index Kit (Illumina, San Diego, CA) and quantified them using Qubit (ThermoFisher Scientific, Waltham, MA) and assessed quality using a Bioanalyzer (Agilent, Santa Clara, CA). We sequenced the brain sample on the Illumina NextSeq 500 (75bp paired-end reads) at the Genomics Resource Laboratory, University of Massachusetts, Amherst. For the first two samples (bulk embryonic stages and juveniles), total RNA was extracted using TRIzol (Ambion) following the standard protocol, quality was assessed using Tapestation (Agilent) and sent to the IGM UCSD Genomic Center for library preparation (TruSeq mRNA stranded library) and sequencing (Illumina NovaSeq 6000; 150bp paired-end reads).

### Reference Transcriptome Construction

*Berghia* samples used for reference transcriptome construction included a subset of samples to minimize computational cost while maximizing read breadth. These included single samples from multiple adult tissues, selected at random, including the (1) brain (SRR12072211), (2) oral tentacle (SRR12072210), (3) rhinophore (SRR12072209), (4) foot (SRR12072206), (5) tail (SRR12072205), (6) proximal (SRR12072208) and (7) distal ceras (SRR12072207), as well as samples (8) embryos (SRR12072213), and (9) juveniles (SRR12072212). We merged all FASTQ output files for the above samples into two files (Read 1 and Read 2) for downstream analysis. We used default parameters for all programs unless otherwise specified. We trimmed and filtered reads using fastp (version 0.20.0; [79]), and assembled transcripts using Trinity (version 2.9.1; [80]). We predicted open reading frames (ORFs) with TransDecoder (version 5.5.0; [81]). Duplication levels were quite high (∼56%), so we clustered predicted ORFs using CD-HIT-EST (version 4.8.1; [82, 83]) at 95% identity and word size of 11 (-c 0.95, -n 11). Post-clustering, we filtered transcripts with alien_index (https://github.com/josephryan/alien_index); based on an algorithm described in [84]). We constructed alien index databases using previously constructed metazoan and non-metazoan databases (obtained from http://ryanlab.whitney.ufl.edu/downloads/alien_index) and all “*Symbiodinium”* sequences present on UniProt [85] as of 31 March 2020. We removed all sequences with an alien index greater than 45 from the transcriptome. We then compiled full transcripts for each predicted ORF sequence remaining from the assembled transcriptome using a custom Python script (full_transcripts.py). We then scanned the transcriptome for vectors and possible contaminants via the NCBI VecScreen (https://www.ncbi.nlm.nih.gov/tools/vecscreen/). We removed vectors using a small script (trim_adapters.pl) available through the Trinity Community Codebase (https://github.com/trinityrnaseq/trinity_community_codebase). We removed or trimmed sequences containing contamination using the Contaminants.txt file provided by NCBI and a custom script (remove_contamination.py). Custom scripts are available at https://github.com/lyons-lab/berghia_reference_transcriptome). We assessed transcriptome quality across all steps using BUSCO v5.1.2 [86–88] scores by comparing assembled transcripts to the metazoa_odb10 (C:98.1%[S:83.1%,D:15.0%],F:0.9%,M:1.0%,n:954) and mollusca_odb10 (C:93.4%[S:76.9%,D:16.5%],F:1.3%,M:5.3%,n:5295) databases.

### Long read RNA sample collection and sequencing

We obtained *Berghia* adult tissue samples from animals starved for at least four weeks to minimize gut contaminants, including the (1) head (one animal), (2) oral tentacles (two animals), (3) rhinophores (three animals), (4) cerata (one animal), (5) mantle (one animal), and (6) homogenized mid-body tissue (one animal). We also collected two developmental samples, including (1) embryos from the trochophore (72 hours post oviposition; 300 animals) and eyed veliger stages (9-10 days post oviposition; 120 animals), and (2) post-metamorphic and post-feeding juveniles (34 animals). We extracted total RNA using the standard TRIzol Reagent (Life Technologies, Carlsbad, CA, USA) protocol, with some modifications: After the addition of chloroform, we centrifuged samples for 20 minutes at max speed (16,000 RCF) and precipitated samples in 100% isopropanol for ∼1 hour at -20°C. We assessed total RNA sample quality on a 1% agarose gel and quantified the RNA in each sample with a Qubit 2.0 High Sensitivity kit (ThermoFisher Scientific, Waltham, MA). We then pooled developmental stages (embryos and juveniles; DEV), adult rhinophore and oral tentacle samples (RHOT), and adult mantle and cerata samples (MCE) in equivalent amounts. We sent these five total RNA samples (DEV, MCE, RHOT, head, mid-body) to the Roy J. Carver Biotechnology Center at the University of Illinois at Urbana-Champaign for IsoSeq library construction (5 libraries) and sequencing. They performed sequencing (on the five pooled libraries) on a single SMRT 8M cell with the PacBio Sequel II (PacBio, Menlo Park, CA) and a 30h movie. They then clustered the raw subreads using the IsoSeq v3 clustering workflow (https://github.com/PacificBiosciences/IsoSeq/blob/master/isoseq-clustering.md).

### Genome assembly and scaffolding

Dovetail used 167 gigabase-pairs of PacBio CLR reads as an input to WTDBG2 v2.5 [89] with genome size 2.0g, minimum read length 20000, and minimum alignment length 8192. Additionally, they enabled realignment with the -R option and set read type with the option -x sq. They then used BLASTn results of the WTDBG2 output assembly against the nt database as input for blobtools v1.1.1, and scaffolds identified as possible contamination were removed from the assembly. Finally, they used purge_dups v1.2.3 [90] to remove haplotigs and contig overlaps.

Dovetail used input de novo assembly and Dovetail Omni-C library reads (3 samples; SRR25687005-SRR25687007) as input data for HiRise, a software pipeline designed specifically for using proximity ligation data to scaffold genome assemblies [91]. They aligned Dovetail Omni-C library sequences to the draft input assembly using BWA with default parameters [92]. They then analyzed separations of Dovetail Omni-C read pairs mapped within draft scaffolds by Hi-Rise to produce a likelihood model for genomic distance between read pairs, and used the model to identify and break putative misjoins, to score prospective joins, and make joins above a threshold.

We initially filtered the *Berghia stephanieae* genome with purge_dups v1.2.5 [90] to automatically identify and remove haplotigs and contig/scaffold overlaps from heterozygous sites. This Whole Genome Shotgun project has been deposited at DDBJ/ENA/GenBank under the accession JAWQJI000000000. The version described in this paper is version JAWQJI010000000. Following duplicate purging, we assessed completeness with BUSCO v5.1.2 [86–88] by comparing to metazoa_odb10 and mollusca_odb10. BUSCO further used the programs HMMER v3.1 [93] and MetaEuk v4.a0f584d [94] for gene prediction and analysis. We then used Nucleotide-Nucleotide BLAST 2.11.0+ [95, 96] to compare our scaffolds to the nt database (downloaded April 2021), and mapped the original PacBio reads used for assembly via minimap2 v2.18-r1015 [97]. With these results, we used BlobToolKit (Challis et al. 2020)(blobtools2 filter option) to remove additional scaffolds considered contamination. Scaffold selection for removal was based on GC content, PacBio read coverage results, BLASTn hits (we removed no-hit and bacterial contamination), and finally a minimum size threshold (150 kb). This size threshold was selected because it was the point at which the removal of sequences would not change the BUSCO score, as determined by the use of BlobToolKit Viewer v1.1 [98]. Most removed sequences contained differences in GC content and coverage compared to those that were retained in the final annotated genome, in addition to their smaller size. We also performed a Nucleotide-Nucleotide BLAST 2.11.0+ to compare the removed sequences with the final genome to assess duplication rates. Of those sequences removed from the final genome, 92.8% hit to one of the final 18 scaffolds (98.7% of which with an e-value of 0.0), and 1.6% of removed sequences were obvious contaminants. To compare the *Berghia* genome with other available Mollusca genomes, we downloaded assembled genomes from NCBI using the datasets command line function (v.15.24.0)[99] with flags for the taxon Mollusca and genomes at assembly levels “scaffold”, “chromosome”, or “complete”. We then assessed completeness of each genome with BUSCO v5.1.2 compared to the database metazoa_odb10, and assessed scaffold N50 using the command line tool n50 in SeqFu, a suite of FASTX utilities [100].

### Gene prediction and annotation

We analyzed filtered genome scaffolds with RepeatModeler v2.0.1 [101], which used Tandem Repeat Finder (TRF) v4.09 [102], RECON v1.08 [103], RepeatScout v1.0.6 [104], and RepeatMasker v4.1.2 (https://www.repeatmasker.org), to construct a de novo repeat library for *Berghia stephanieae*. This included the ---LTRStruct flag to run an LTR Structural Analysis, which used GenomeTools v1.6.1 (http://genometools.org), LTR_Retriever v2.9.0 [105], Ninja v1.10.2 (https://github.com/ninja-build/ninja), MAFFT v7.480 [106], and CD-HIT v4.8.1 [82, 83]. We then used this species-specific library to detect repeat sequences (via both soft and hardmasking) with RepeatMasker in the *Berghia* genome.

Following repeat masking, we mapped all short RNA-seq reads (unfiltered) to the hardmasked version of the genome using two separate read mapping software programs, HiSat2 v2.2.1 [107] and STAR v2.7.9a [108]. This was intended to account for mapping bias in order to maximize the possibility for support in our gene annotations. For long read (IsoSeq) mapping, we first obtained Full Length Non-Concatemer (FLNC) reads from step three of the IsoSeq v3 workflow. These FLNC reads were mapped directly to the non-repeatmasked genome using minimap2 v2.22-r1105-dirty [97] with the recommended options according to PacBio (https://github.com/Magdoll/cDNA_Cupcake/wiki/Best-practice-for-aligning-Iso-Seq-to-reference-genome:-minimap2,-deSALT,-GMAP,-STAR,-BLAT). Post-mapping, sam output files were reformatted into bam files and indexed using samtools v1.11[109]. We used BRAKER v2.1.6 [110–112] for preliminary gene prediction of the *Berghia* genome, which uses GeneMark-EP+ v4 [113, 114], DIAMOND v2.0.8 [115], spaln v2.4.3 [116, 117], and Augustus v3.4.0 [118].

We used long and short read RNA-seq mapping results used as expression support input. We also used a protein hints file generated by combining the mollusca_odb10 database with *B.stephanieae* sequences identified as BUSCO hits from our initial mollusca_odb10 BUSCO run. We ran BRAKER with the additional flags --etpmode, --gff3 and --softmasking. After initial gene prediction, we generated a filtered predicted gene set using a script included with the BRAKER installation (selectSupportedSubsets.py) and the --anySupport flag to only include genes at least partially supported by hints. IsoSeq and short read RNA-sequencing data were mapped to both sets of gene models to assess the impact of filtering. Both unfiltered and filtered gene prediction results are provided in Dryad (DOI: 10.6076/D1BS33), but we only used the filtered set (braker_annotations_anysupport.gff3) in subsequent functional annotation and clade-specific gene analyses.

For functional annotation of predicted genes, we used Protein-Protein BLAST 2.11.0+ (BLASTP) and InterProScan v.5.52-86.0. For BLASTP analyses, we used an e-value cutoff of 1e-3 with -max_target_seqs of one against three databases: (1) UniProtKB/Swiss-Prot, (2) RefSeq, and (3) Trembl (all downloaded April 2021). We then combined the hits to all three databases in a single blast annotation file. For InterProScan analyses, we used the default parameters with some additional flags, including: -goterms to look up gene ontology, -dp to disable pre-calculated match lookup, and -t p to indicate protein sequences.

### Assessment of clade-specific genes and their expression

To determine the distribution of clade-specific genes for *Berghia*, we created a proteome dataset containing 47 metazoan species using both published genomes and transcriptomes (Additional file 4: Table S3). Our final dataset included 36 molluscs, including fourteen nudibranchs (five from Anthobranchia and nine from Cladobranchia). We downloaded predicted proteomes from genome datasets from MolluscDB [52]. We downloaded the transcriptomes from the NCBI Sequence Read Archive (SRA), which we then filtered with fastp v0.20.0 [79] and assembled with Trinity v2.9.1; [80]. We predicted ORFs using Transdecoder v5.5.0 (https://github.com/TransDecoder/TransDecoder), with default parameters. Our final proteomes ranged from 17,606 to 72,541 proteins (x = 33,691; Additional file 4: Table S3). We identified orthologous gene families among our metazoan proteomes using the OrthoFinder package (Emms and Kelly, 2019) with default parameters and a user generated species tree as input (Additional File 11). Our user generated species tree topology was based on recent metazoan phylogenies [119–122], the MolluscDB phylogeny provided on the website [52], and recent Mollusca [123] and nudibranch [124–126] phylogenetic analyses. We then analyzed orthologous groups using the program KinFin v1.0.3 [127] to determine which predicted genes in *Berghia stephanieae* are clade-specific (meaning that they only cluster with sequences from a particular clade). We used the default parameters, with additional flags (--infer_singletons --plot_tree -r phylum,class,order,superfamily).

To determine the expression patterns of clade-specific genes, we mapped our short read RNAseq data (from the brain, oral tentacles, rhinophores, foot, tail, and proximal and distal ceras) to the *Berghia* genome using STAR v2.7.9a [108] with default parameters plus additional flags (--readFilesCommand zcat --outSAMtype BAM SortedByCoordinate --twopassMode Basic--sjdbGTFfeatureExon ‘CDS’). We counted reads using the command htseq-count from the HTSeq framework v1.99.2 [128], which is a Python package for analysis of high-throughput sequencing data. We analyzed counts using the DESeq function from DESeq2 v1.26.0 [129] to perform differential analysis, and generated results using the results function with contrasts comparing each focal tissue with an average of all other tissues. We considered genes upregulated if the adjusted p-value (padj) was greater than 0.05 and log2FoldChange was greater than 2.

### *In situ* Hybridization Chain Reaction (HCR) in *Berghia* juveniles

#### Probe Design

We designed all probe sets using the HCR 3.0 probe maker [130]. The sequences generated by the software were used to order probe sets (50Lpmol DNA oPools Oligo Pool) from Integrated DNA Technologies (Coralville, IA USA), which we resuspended to 1Lpmol/μl in 50LmL TE buffer (Tris, EDTA).

#### Hybridization Chain Reaction

We cultured *Berghia stephanieae* juveniles using the same methods as prior *B. stephanieae* imaging work [35, 36]. We starved juveniles for five days prior to fixation to decrease autofluorescence from digestive contents. We relaxed juveniles in 1 part 7.3% MgCl2 : 2 part fresh filtered sea water for 30 minutes prior to fixation. We then washed and incubated samples in 4% PFA (diluted in FSW from 16% ampules) overnight at 4L. We washed samples three times in 1X PBS, followed by a 50% 1X PBS/ 50% methanol solution wash, followed by three 100% methanol washes. All washes were 10 minutes. We stored samples in methanol at -20C.

We performed *in situ* HCR using the buffers and protocols detailed in Choi et al. [50] with the following modifications. All steps were performed in 1.5mL tubes and the volume of washes was decreased to 200µL to better suit our samples. We rehydrated samples into 5X SSCT from methanol, immediately followed by the detection stage of the protocol. We prepared probe solutions using 100µL of hybridization buffer and 1.0 pmol/oligo/µL of each probe. Following overnight probe hybridization, we washed samples with 30% probe hybridization wash buffer for 3 x 5 minutes, followed by 2 x 30 minutes. Following the 5X SSCT washes during the amplification stage, we placed samples in the hairpin solution (6pmol solution using 2µL of 3µM stock of snap-cooled hairpins in 100 µL of amplification buffer). After the completion of the HCR protocol, we incubated the samples in 1.0 µg/mL DAPI diluted in 5X SSCT for 30 minutes. Samples were then stored in 5X SSCT at 4 C for up to 5 days until mounting and imaging.

We mounted samples in a 20% 5X SSCT, 80% Glycerol solution and imaged samples with a Zeiss LSM 710 inverted confocal microscope with an AxioCam HRm camera. We analyzed images using image processing software ImageJ FIJI and Adobe Photoshop [131]. Samples were stitched together using the FIJI Pairwise Stitching Plugin [132]. Figures were created in Adobe Illustrator.

## Supporting information

supplements version 2

## Declarations

### Ethics approval and consent to participate

Not applicable.

### Consent for publication

Not applicable.

### Availability of data and materials

Raw sequencing data are accessible through the NCBI Sequence Read Archive (BioProjects PRJNA1004233 and PRJNA641185). This Whole Genome Shotgun project has been deposited at DDBJ/ENA/GenBank under the accession JAWQJI000000000. The version described in this paper is version JAWQJI010000000. The full commands we used for filtering and annotation steps are provided in Additional File 12. Input and intermediate files for our analyses are available in Dryad (https://doi.org/10.6076/D1BS33) and custom python, shell, and R scripts used in our analyses are available through Github (https://github.com/lyons-lab/berghia_reference_transcriptome and https://github.com/goodheart-lab/berghia_genome).

### Authors’ contributions

JAG, MDR, PSK and DCL conceived of the study; JAG, NFT, RAF, SRB, KM, MACJ, CW, PM, GOB, HTJ, MDR and DCL collected data; JAG, RAR, and RAF performed data analyses; JAG, MDR, RAF, PSK and DCL participated in data interpretation. All authors read and approved the final manuscript.

## Acknowledgements

We thank all members of the Lyons and Katz labs for help with animal care and feedback on this project and manuscript. We are also grateful to all members of the *Berghia* Brain Project (https://sites.google.com/umass.edu/berghiabrainproject/) for their feedback on this project. We also thank the Hamdoun Lab for the use of their confocal microscope and Rouse Lab for the use of their Qubit. This work is in memory of our outstanding colleague Dr. Maryna P. Lesoway, who provided significant intellectual and emotional support in executing this work.

## Supplementary Files

**Additional File 1.** *Berghia stephanieae* HiRise Scaffolding report from Dovetail.

**Additional File 2: Figure S1.** K-mer spectra and fitted models for a GenomeScope analysis of our Dovetail Omni-C data for *Berghia stephanieae*. **Figure S2.** Snail plot showing high contiguity of the filtered genome (18 scaffolds), where 99% of the genome is found in the top 15 scaffolds. This plot also indicates the high levels of completeness (93.3%) when compared to metazoan single-copy ortholog datasets (BUSCO), with low levels of duplication (0.6%). **Figure S3.** This chart shows the proteins in each proteome that were classified as species-specific based on the OrthoFinder and KinFin analyses. **Figure S4.** This chart shows the proportion of proteins in each proteome that were classified as species-specific based on the OrthoFinder and KinFin analyses. **Figure S5.** Rarefaction curve for gene sampling at the phylum level. This graph indicates that gene sampling for Mollusca (green) is reaching an asymptote, which suggests that our sampling of these genes is sufficient for identifying *Berghia* genes that would match to other Mollusca proteomes. **Figure S6.** Rarefaction curve for gene sampling at the class level. This graph indicates that gene sampling for Gastropoda (blue) and Bivalvia (green) are reaching an asymptote, which suggests that our sampling of these genes is sufficient for identifying *Berghia* genes that would match to other gastropod proteomes. **Figure S7.**

Rarefaction curve for gene sampling at the subclass level. This graph indicates that gene sampling for Heterobranchia (purple) are reaching an asymptote, which suggests that our sampling of these genes is sufficient for identifying *Berghia* genes that would match to other gastropod proteomes. **Figure S8.** Rarefaction curve for gene sampling at the order level. This graph indicates that gene sampling for Nudibranchia (red) is reaching an asymptote, which suggests that our sampling of these genes is sufficient for identifying *Berghia* genes that would match to other nudibranch proteomes. **Figure S9.** Rarefaction curve for gene sampling at the superfamily level. This graph indicates that gene sampling for Aeolidoidea is reaching an asymptote, though this group is not as well sampled as other levels. This still suggests that our sampling of these genes is sufficient for identifying *Berghia* genes that would match to other aeolid proteomes. **Figure S10.** Distribution of both clade-specific and non-clade-specific (Other) genes across the putative chromosomes in our *Berghia stephanieae* genome. To the left is the raw gene counts per chromosome, and to the right is the number of genes per Mb of sequence to account for differences in chromosome lengths. **Figure S11.** This chart shows the proportion of genes on each chromosome that are clade-specific and non-clade-specific (Other). **Figures S12-18.** Gene expression across tissues in genes classified as *Berghia*-specific (**Figure S12**), Aeolidina-specific (**Figure S13**), Nudibranchia-specific (**Figure S14**), Heterobranchia-specific (**Figure S15**),Gastropoda-specific (**Figure S16**), Mollusca-specific (**Figure S17**), and Other (**Figure S18**). Left: Heatmap of expression profiles; Right: PCA-plot showing similarity of expression within and among tissues.

**Additional File 3: Table S1.** Genome summary statistics for various mollusk genomes available in NCBI and both versions of MolluscDB.

**Additional File 4: Table S2.** RepeatModeler analysis for the Berghia stephanieae genome.

**Additional File 5: Table S3.** OrthoFinder cluster analysis statistics for each proteome.

**Additional File 6: Table S4.** Read statistics for short read transcriptome samples for *Berghia stephanieae*.

**Additional File 7: Table S5**. Number of genes upregulated in each tissue, categorized by assigned clade-specificity of each gene. **Table S6.** Percentage of genes upregulated in each tissue, categorized by assigned clade-specificity of each gene.

**Additional File 8: Table S7.** Number of genes uniquely upregulated in each tissue, categorized by assigned clade-specificity of each gene.**Table S8.** Percentage of genes uniquely upregulated in each tissue, categorized by assigned clade-specificity of each gene.

**Additional File 9: Table S9.** Number and proportion of upregulated genes in each tissue that were classified as annotated, unannotated, or uncharacterized.

**Additional File 10: Table S10.** GO term enrichment results for genes upregulated in the brain. **Table S11**. GO term enrichment results for genes upregulated in the rhinophores. **Table S12.** GO term enrichment results for genes upregulated in the oral tentacles. **Table S13.** GO term enrichment results for genes upregulated in the distal ceras. **Table S14.** GO term enrichment results for genes upregulated in the proximal ceras. **Table S15.** GO term enrichment results for genes upregulated in the foot. **Table S16.** GO term enrichment results for genes upregulated in the tail.

**Additional File 11.** Newick file of the species tree used in our OrthoFinder analysis.

**Additional File 12.** List of commands used in the *Berghia* genome annotation.

## References

1. Wilkins AS. “the genetic tool-kit”: The life-history of an important metaphor. In: Advances in Evolutionary Developmental Biology. Hoboken, NJ, USA: John Wiley & Sons, Inc.; 2013. p. 1–14.

2. Carroll SB. Evo-devo and an expanding evolutionary synthesis: a genetic theory of morphological evolution. Cell. 2008;134:25–36.

3. Newman SA. The developmental genetic toolkit and the molecular homology—analogy paradox. Biol Theory. 2006;1:12–6.

4. de Mendoza A, Sebé-Pedrós A, Ruiz-Trillo I. The evolution of the GPCR signaling system in eukaryotes: modularity, conservation, and the transition to metazoan multicellularity. Genome Biol Evol. 2014;6:606–19.

5. Wu L, Lambert JD. Clade-specific genes and the evolutionary origin of novelty; new tools in the toolkit. Semin Cell Dev Biol. 2022. 10.1016/j.semcdb.2022.05.025.

6. Johnson BR. Taxonomically Restricted Genes Are Fundamental to Biology and Evolution. Front Genet. 2018;9:407.

7. Khalturin K, Hemmrich G, Fraune S, Augustin R, Bosch TCG. More than just orphans: are taxonomically-restricted genes important in evolution? Trends Genet. 2009;25:404–13.

8. Valdés A. A new species of *Aeolidiella* Bergh, 1867 (Mollusca: Nudibranchia: Aeolidiidae) from the Florida keys, USA. Veliger. 2005;47:218–23.

9. Kurz EM, Holstein TW, Petri BM, Engel J, David CN. Mini-collagens in *Hydra* nematocytes. J Cell Biol. 1991;115:1159–69.

10. Koch AW, Holstein TW, Mala C, Kurz E, Engel J, David CN. Spinalin, a new glycine- and histidine-rich protein in spines of *Hydra* nematocysts. J Cell Sci. 1998;111 (Pt 11):1545–54.

11. Babonis LS, Martindale MQ. Old cell, new trick? Cnidocytes as a model for the evolution of novelty. Integr Comp Biol. 2014;54:714–22.

12. Wu L, Hiebert LS, Klann M, Passamaneck Y, Bastin BR, Schneider SQ, et al. Genes with spiralian-specific protein motifs are expressed in spiralian ciliary bands. Nat Commun. 2020;11:4171.

13. Garb JE, Ayoub NA, Hayashi CY. Untangling spider silk evolution with spidroin terminal domains. BMC Evol Biol. 2010;10:243.

14. Hinman MB, Lewis RV. Isolation of a clone encoding a second dragline silk fibroin. Nephila clavipes dragline silk is a two-protein fiber. J Biol Chem. 1992;267:19320–4.

15. Chen S, Zhang YE, Long M. New genes in *Drosophila* quickly become essential. Science. 2010;330:1682–5.

16. True JR, Haag ES. Developmental system drift and flexibility in evolutionary trajectories. Evol Dev. 2001;3:109–19.

17. Hwang JS, Takaku Y, Momose T, Adamczyk P, Özbek S, Ikeo K, et al. Nematogalectin, a nematocyst protein with GlyXY and galectin domains, demonstrates nematocyte-specific alternative splicing in *Hydra*. Proc Natl Acad Sci U S A. 2010;107:18539–44.

18. Khalturin K, Anton-Erxleben F, Sassmann S, Wittlieb J, Hemmrich G, Bosch TCG. A novel gene family controls species-specific morphological traits in *Hydra*. PLoS Biol. 2008;6:e278.

19. Milde S, Hemmrich G, Anton-Erxleben F, Khalturin K, Wittlieb J, Bosch TCG. Characterization of taxonomically restricted genes in a phylum-restricted cell type. Genome Biol. 2009;10:R8.

20. Santos ME, Le Bouquin A, Crumière AJJ, Khila A. Taxon-restricted genes at the origin of a novel trait allowing access to a new environment. Science. 2017;358:386–90.

21. Jasper WC, Linksvayer TA, Atallah J, Friedman D, Chiu JC, Johnson BR. Large-scale coding sequence change underlies the evolution of postdevelopmental novelty in honey bees. Mol Biol Evol. 2015;32:334–46.

22. Babonis LS, Martindale MQ, Ryan JF. Do novel genes drive morphological novelty? An investigation of the nematosomes in the sea anemone *Nematostella vectensis*. BMC Evol Biol. 2016;16:114.

23. Saleuddin S, Mukai S. Physiology of Molluscs: A Collection of Selected Reviews, Volume 1. CRC Press; 2021.

24. Albertin CB, Simakov O, Mitros T, Wang ZY, Pungor JR, Edsinger-Gonzales E, et al. The octopus genome and the evolution of cephalopod neural and morphological novelties. Nature. 2015;524:220–4.

25. Styfhals R, Zolotarov G, Hulselmans G, Spanier KI, Poovathingal S, Elagoz AM, et al. Cell type diversity in a developing octopus brain. Nat Commun. 2022;13:7392.

26. Metzger MJ, Villalba A, Carballal MJ, Iglesias D, Sherry J, Reinisch C, et al. Widespread transmission of independent cancer lineages within multiple bivalve species. Nature. 2016;534:705–9.

27. Katz PS, Quinlan PD. The importance of identified neurons in gastropod molluscs to neuroscience. Curr Opin Neurobiol. 2019;56:1–7.

28. Byrne JH. Learning and Memory in Aplysia and Other Invertebrates. In: Neurobiology of Comparative Cognition. 1st Edition. Psychology Press; 2014. p. 311–34.

29. Coustau C, Gourbal B, Duval D, Yoshino TP, Adema CM, Mitta G. Advances in gastropod immunity from the study of the interaction between the snail *Biomphalaria glabrata* and its parasites: A review of research progress over the last decade. Fish Shellfish Immunol. 2015;46:5–16.

30. Coustau C, Kurtz J, Moret Y. A Novel Mechanism of Immune Memory Unveiled at the Invertebrate-Parasite Interface. Trends in parasitology. 2016;32:353–5.

31. Davison A, Neiman M. Mobilizing molluscan models and genomes in biology. Philos Trans R Soc Lond B Biol Sci. 2021;376:20200163.

32. Gomes-dos-Santos A, Lopes-Lima M, Castro LFC, Froufe E. Molluscan genomics: the road so far and the way forward. Hydrobiologia. 2020;847:1705–26.

33. Wägele H, Willan RC. Phylogeny of the Nudibranchia. Zoological Journal of the Linnean Society. 2000;130:83–181.

34. Kristof A, Klussmann-Kolb A. Neuromuscular development of *Aeolidiella stephanieae* Valdez, 2005 (Mollusca, Gastropoda, Nudibranchia). Frontiers in Zoology. 2010;7:5.

35. Taraporevala NF, Lesoway MP, Goodheart JA, Lyons DC. Precocious sperm exchange in the simultaneously hermaphroditic nudibranch, Berghia stephanieae. Integrative Organismal Biology. 2022;4:obac030.

36. Goodheart JA, Barone V, Lyons DC. Movement and storage of nematocysts across development in the nudibranch *Berghia stephanieae* (Valdés, 2005). Front Zool. 2022;19:16.

37. Obermann D, Bickmeyer U, Wägele H. Incorporated nematocysts in Aeolidiella stephanieae (Gastropoda, Opisthobranchia, Aeolidoidea) mature by acidification shown by the pH sensitive fluorescing alkaloid Ageladine A. Toxicon. 2012;60:1108–16.

38. Silva RXG, Cartaxana P, Calado R. Prevalence and Photobiology of Photosynthetic Dinoflagellate Endosymbionts in the Nudibranch *Berghia stephanieae*. Animals. 2021;11:2200.

39. Clavijo Melo J, Sickinger C, Bleidißel S, Gasparoni G, Tierling S, Preisfeld A, et al. The nudibranch *Berghia stephanieae* (Valdés, 2005) is not able to initiate a functional symbiosome- like environment to maintain *Breviolum minutum* (J.E.Parkinson & LaJeunesse 2018). Frontiers in Marine Science. 2022;9.

40. Martín-Durán JM, Hejnol A. A developmental perspective on the evolution of the nervous system. Dev Biol. 2021;475:181–92.

41. Goodheart JA, Bely AE. Sequestration of nematocysts by divergent cnidarian predators: mechanism, function, and evolution. Invertebrate Biology. 2017;136:75–91.

42. Ranallo-Benavidez TR, Jaron KS, Schatz MC. GenomeScope 2.0 and Smudgeplot for reference-free profiling of polyploid genomes. Nat Commun. 2020;11:1432.

43. Cai H, Li Q, Fang X, Li J, Curtis NE, Altenburger A, et al. A draft genome assembly of the solar-powered sea slug *Elysia chlorotica*. Sci Data. 2019;6:190022.

44. Sun J, Zhang Y, Xu T, Zhang Y, Mu H, Zhang Y, et al. Adaptation to deep-sea chemosynthetic environments as revealed by mussel genomes. Nat Ecol Evol. 2017;1:121.

45. Uliano-Silva M, Dondero F, Dan Otto T, Costa I, Lima NCB, Americo JA, et al. A hybrid-hierarchical genome assembly strategy to sequence the invasive golden mussel, *Limnoperna fortunei*. Gigascience. 2018;7.

46. Du X, Fan G, Jiao Y, Zhang H, Guo X, Huang R, et al. The pearl oyster *Pinctada fucata martensii* genome and multi-omic analyses provide insights into biomineralization. Gigascience. 2017;6:1–12.

47. Jiao W, Fu X, Dou J, Li H, Su H, Mao J, et al. High-resolution linkage and quantitative trait locus mapping aided by genome survey sequencing: building up an integrative genomic framework for a bivalve mollusc. DNA Res. 2014;21:85–101.

48. Thiriot-Quiévreux C. Advances in chromosomal studies of gastropod molluscs. J Molluscan Stud. 2003;69:187–202.

49. Law CW, Alhamdoosh M, Su S, Dong X, Tian L, Smyth GK, et al. RNA-seq analysis is easy as 1-2-3 with limma, Glimma and edgeR. F1000Res. 2016;5.

50. Choi HMT, Schwarzkopf M, Fornace ME, Acharya A, Artavanis G, Stegmaier J, et al. Third-generation hybridization chain reaction: multiplexed, quantitative, sensitive, versatile, robust. Development. 2018;145.

51. Ramirez MD, Bui TN, Katz PS. Mapping neuronal gene expression reveals aspects of ganglionic organization in a gastropod mollusc. bioRxiv. 2023;:2023.06.22.546160.

52. Liu F, Li Y, Yu H, Zhang L, Hu J, Bao Z, et al. MolluscDB: an integrated functional and evolutionary genomics database for the hyper-diverse animal phylum Mollusca. Nucleic Acids Res. 2021;49:D988–97.

53. Farhat S, Bonnivard E, Pales Espinosa E, Tanguy A, Boutet I, Guiglielmoni N, et al. Comparative analysis of the *Mercenaria mercenaria* genome provides insights into the diversity of transposable elements and immune molecules in bivalve mollusks. BMC Genomics. 2022;23:192.

54. Holmes A, Derbyshire T, Brennan M, McTierney S, Small A, Marine Biological Association Genome Acquisition Lab, et al. The genome sequence of *Gari tellinella* (Lamarck, 1818), a sunset clam. Wellcome Open Research. 2022;7:116.

55. Chen Z, Doğan Ö, Guiglielmoni N, Guichard A, Schrödl M. The *de novo* genome of the “Spanish” slug *Arion vulgaris* Moquin-Tandon, 1855 (Gastropoda: Panpulmonata): massive expansion of transposable elements in a major pest species. bioRxiv. 2020.

56. Chueca LJ, Schell T, Pfenninger M. De novo genome assembly of the land snail *Candidula unifasciata* (Mollusca: Gastropoda). G3. 2021;11.

57. Linscott TM, González-González A, Hirano T, Parent CE. De novo genome assembly and genome skims reveal LTRs dominate the genome of a limestone endemic Mountainsnail (*Oreohelix idahoensis*). BMC Genomics. 2022;23:796.

58. Li L, Stoeckert CJ Jr, Roos DS. OrthoMCL: identification of ortholog groups for eukaryotic genomes. Genome Res. 2003;13:2178–89.

59. Emms DM, Kelly S. OrthoFinder: solving fundamental biases in whole genome comparisons dramatically improves orthogroup inference accuracy. Genome Biol. 2015;16:157.

60. Emms DM, Kelly S. OrthoFinder: phylogenetic orthology inference for comparative genomics. Genome Biol. 2019;20:238.

61. Weisman CM, Murray AW, Eddy SR. Many, but not all, lineage-specific genes can be explained by homology detection failure. PLoS Biol. 2020;18:e3000862.

62. Aguilera F, McDougall C, Degnan BM. Co-Option and De Novo Gene Evolution Underlie Molluscan Shell Diversity. Mol Biol Evol. 2017;34:779–92.

63. Tautz D, Domazet-Lošo T. The evolutionary origin of orphan genes. Nat Rev Genet. 2011;12:692–702.

64. Johnson BR, Tsutsui ND. Taxonomically restricted genes are associated with the evolution of sociality in the honey bee. BMC Genomics. 2011;12:164.

65. Hilgers L, Hartmann S, Hofreiter M, von Rintelen T. Novel Genes, Ancient Genes, and Gene Co-Option Contributed to the Genetic Basis of the Radula, a Molluscan Innovation. Mol Biol Evol. 2018;35:1638–52.

66. Goodheart JA, Bleidißel S, Schillo D, Strong EE, Ayres DL, Preisfeld A, et al. Comparative morphology and evolution of the cnidosac in Cladobranchia (Gastropoda: Heterobranchia: Nudibranchia). Front Zool. 2018;15:43.

67. Davy SK, Allemand D, Weis VM. Cell biology of cnidarian-dinoflagellate symbiosis. Microbiol Mol Biol Rev. 2012;76:229–61.

68. Brenzinger B, Schrödl M, Kano Y. Origin and significance of two pairs of head tentacles in the radiation of euthyneuran sea slugs and land snails. Sci Rep. 2021;11:21016.

69. Huber G. On the cerebral nervous system of marine Heterobranchia (Gastropoda). J Molluscan Stud. 1993;59:381–420.

70. Striedter GF. Principles of Brain Evolution. Sinauer Associates Incorporated; 2005.

71. Vickaryous MK, Hall BK. Human cell type diversity, evolution, development, and classification with special reference to cells derived from the neural crest. Biol Rev Camb Philos Soc. 2006;81:425–55.

72. Moroz LL. On the independent origins of complex brains and neurons. Brain Behav Evol. 2009;74:177–90.

73. Shafer MER, Sawh AN, Schier AF. Gene family evolution underlies cell-type diversification in the hypothalamus of teleosts. Nat Ecol Evol. 2022;6:63–76.

74. Hwang JS, Ohyanagi H, Hayakawa S, Osato N, Nishimiya-Fujisawa C, Ikeo K, et al. The evolutionary emergence of cell type-specific genes inferred from the gene expression analysis of *Hydra*. Proc Natl Acad Sci U S A. 2007;104:14735–40.

75. Greenwood PG, Mariscal RN. The utilization of cnidarian nematocysts by aeolid nudibranchs: nematocyst maintenance and release in *Spurilla*. Tissue Cell. 1984;16:719–30.

76. Rosin FM, Kramer EM. Old dogs, new tricks: regulatory evolution in conserved genetic modules leads to novel morphologies in plants. Dev Biol. 2009;332:25–35.

77. Mikheyev AS, Linksvayer TA. Genes associated with ant social behavior show distinct transcriptional and evolutionary patterns. Elife. 2015;4:e04775.

78. Yang L, Zou M, Fu B, He S. Genome-wide identification, characterization, and expression analysis of lineage-specific genes within zebrafish. BMC Genomics. 2013;14:65.

79. Chen S, Zhou Y, Chen Y, Gu J. fastp: an ultra-fast all-in-one FASTQ preprocessor. Bioinformatics. 2018;34:i884–90.

80. Grabherr MG, Haas BJ, Yassour M, Levin JZ, Thompson DA, Amit I, et al. Full-length transcriptome assembly from RNA-Seq data without a reference genome. Nat Biotechnol. 2011;29:644–52.

81. Haas BJ, Papanicolaou A, Yassour M, Grabherr M, Blood PD, Bowden J, et al. De novo transcript sequence reconstruction from RNA-seq using the Trinity platform for reference generation and analysis. Nat Protoc. 2013;8:1494–512.

82. Li W, Godzik A. Cd-hit: a fast program for clustering and comparing large sets of protein or nucleotide sequences. Bioinformatics. 2006;22:1658–9.

83. Fu L, Niu B, Zhu Z, Wu S, Li W. CD-HIT: accelerated for clustering the next-generation sequencing data. Bioinformatics. 2012;28:3150–2.

84. Gladyshev EA, Meselson M, Arkhipova IR. Massive horizontal gene transfer in bdelloid rotifers. Science. 2008;320:1210–3.

85. UniProt Consortium. The universal protein resource (UniProt). Nucleic Acids Res. 2008;36 Database issue:D190–5.

86. Seppey M, Manni M, Zdobnov EM. BUSCO: Assessing Genome Assembly and Annotation Completeness. Methods Mol Biol. 2019;1962:227–45.

87. Simão FA, Waterhouse RM, Ioannidis P, Kriventseva EV, Zdobnov EM. BUSCO: assessing genome assembly and annotation completeness with single-copy orthologs. Bioinformatics. 2015;31:3210–2.

88. Manni M, Berkeley MR, Seppey M, Simão FA, Zdobnov EM. BUSCO Update: Novel and Streamlined Workflows along with Broader and Deeper Phylogenetic Coverage for Scoring of Eukaryotic, Prokaryotic, and Viral Genomes. Mol Biol Evol. 2021;38:4647–54.

89. Ruan J, Li H. Fast and accurate long-read assembly with wtdbg2. Nat Methods. 2020;17:155–8.

90. Guan D, McCarthy SA, Wood J, Howe K, Wang Y, Durbin R. Identifying and removing haplotypic duplication in primary genome assemblies. Bioinformatics. 2020;36:2896–8.

91. Putnam NH, O’Connell BL, Stites JC, Rice BJ, Blanchette M, Calef R, et al. Chromosome-scale shotgun assembly using an in vitro method for long-range linkage. Genome Research. 2016;26:342–50.

92. Li H, Durbin R. Fast and accurate long-read alignment with Burrows-Wheeler transform. Bioinformatics. 2010;26:589–95.

93. Mistry J, Finn RD, Eddy SR, Bateman A, Punta M. Challenges in homology search: HMMER3 and convergent evolution of coiled-coil regions. Nucleic Acids Research. 2013;41:e121–e121.

94. Levy Karin E, Mirdita M, Söding J. MetaEuk-sensitive, high-throughput gene discovery, and annotation for large-scale eukaryotic metagenomics. Microbiome. 2020;8:48.

95. Camacho C, Coulouris G, Avagyan V, Ma N, Papadopoulos J, Bealer K, et al. BLAST : architecture and applications. BMC Bioinformatics. 2009;10:421.

96. Altschul SF, Gish W, Miller W, Myers EW, Lipman DJ. Basic local alignment search tool. Journal of Molecular Biology. 1990;215:403–10.

97. Li H. Minimap2: pairwise alignment for nucleotide sequences. Bioinformatics. 2018;34:3094–100.

98. Challis R, Richards E, Rajan J, Cochrane G, Blaxter M. BlobToolKit - Interactive Quality Assessment of Genome Assemblies. G3. 2020;10:1361–74.

99. Sayers EW, Bolton EE, Brister JR, Canese K, Chan J, Comeau DC, et al. Database resources of the national center for biotechnology information. Nucleic Acids Res. 2022;50:D20–6.

100. Telatin A, Fariselli P, Birolo G. SeqFu: A Suite of Utilities for the Robust and Reproducible Manipulation of Sequence Files. Bioengineering (Basel). 2021;8.

101. Flynn JM, Hubley R, Goubert C, Rosen J, Clark AG, Feschotte C, et al. RepeatModeler2 for automated genomic discovery of transposable element families. Proc Natl Acad Sci U S A. 2020;117:9451–7.

102. Benson G. Tandem repeats finder: a program to analyze DNA sequences. Nucleic Acids Research. 1999;27:573–80.

103. Bao Z, Eddy SR. Automated de novo identification of repeat sequence families in sequenced genomes. Genome Res. 2002;12:1269–76.

104. Price AL, Jones NC, Pevzner PA. De novo identification of repeat families in large genomes. Bioinformatics. 2005;21 Suppl 1:i351–8.

105. Ou S, Jiang N. LTR_retriever: A Highly Accurate and Sensitive Program for Identification of Long Terminal Repeat Retrotransposons. Plant Physiol. 2018;176:1410–22.

106. Katoh K, Standley DM. MAFFT multiple sequence alignment software version 7: improvements in performance and usability. Mol Biol Evol. 2013;30:772–80.

107. Kim D, Paggi JM, Park C, Bennett C, Salzberg SL. Graph-based genome alignment and genotyping with HISAT2 and HISAT-genotype. Nature Biotechnology. 2019;37:907–15.

108. Dobin A, Davis CA, Schlesinger F, Drenkow J, Zaleski C, Jha S, et al. STAR: ultrafast universal RNA-seq aligner. Bioinformatics. 2013;29:15–21.

109. Li H, Handsaker B, Wysoker A, Fennell T, Ruan J, Homer N, et al. The Sequence Alignment/Map format and SAMtools. Bioinformatics. 2009;25:2078–9.

110. Hoff KJ, Lange S, Lomsadze A, Borodovsky M, Stanke M. BRAKER1: Unsupervised RNA-Seq-Based Genome Annotation with GeneMark-ET and AUGUSTUS. Bioinformatics. 2016;32:767–9.

111. Brůna T, Hoff KJ, Lomsadze A, Stanke M, Borodovsky M. BRAKER2: automatic eukaryotic genome annotation with GeneMark-EP+ and AUGUSTUS supported by a protein database. NAR Genom Bioinform. 2021;3:lqaa108.

112. Hoff KJ, Lomsadze A, Borodovsky M, Stanke M. Whole-Genome Annotation with BRAKER. Methods Mol Biol. 2019;1962:65–95.

113. Brůna T, Lomsadze A, Borodovsky M. GeneMark-EP+: eukaryotic gene prediction with self-training in the space of genes and proteins. NAR Genom Bioinform. 2020;2:lqaa026.

114. Lomsadze A, Ter-Hovhannisyan V, Chernoff YO, Borodovsky M. Gene identification in novel eukaryotic genomes by self-training algorithm. Nucleic Acids Res. 2005;33:6494–506.

115. Buchfink B, Xie C, Huson DH. Fast and sensitive protein alignment using DIAMOND. Nat Methods. 2015;12:59–60.

116. Gotoh O. A space-efficient and accurate method for mapping and aligning cDNA sequences onto genomic sequence. Nucleic Acids Res. 2008;36:2630–8.

117. Iwata H, Gotoh O. Benchmarking spliced alignment programs including Spaln2, an extended version of Spaln that incorporates additional species-specific features. Nucleic Acids Res. 2012;40:e161.

118. Stanke M, Diekhans M, Baertsch R, Haussler D. Using native and syntenically mapped cDNA alignments to improve de novo gene finding. Bioinformatics. 2008;24:637–44.

119. Laumer CE, Fernández R, Lemer S, Combosch D, Kocot KM, Riesgo A, et al. Revisiting metazoan phylogeny with genomic sampling of all phyla. Proc Biol Sci. 2019;286:20190831.

120. Hejnol A, Obst M, Stamatakis A, Ott M, Rouse GW, Edgecombe GD, et al. Assessing the root of bilaterian animals with scalable phylogenomic methods. Proc Biol Sci. 2009;276:4261– 70.

121. Kocot KM, Struck TH, Merkel J, Waits DS, Todt C, Brannock PM, et al. Phylogenomics of Lophotrochozoa with Consideration of Systematic Error. Syst Biol. 2017;66:256–82.

122. Marlétaz F, Peijnenburg KTCA, Goto T, Satoh N, Rokhsar DS. A New Spiralian Phylogeny Places the Enigmatic Arrow Worms among Gnathiferans. Curr Biol. 2019;29:312–8.e3.

123. Sigwart JD, Lindberg DR. Consensus and confusion in molluscan trees: evaluating morphological and molecular phylogenies. Syst Biol. 2015;64:384–95.

124. Karmeinski D, Meusemann K, Goodheart JA, Schroedl M, Martynov A, Korshunova T, et al. Transcriptomics provides a robust framework for the relationships of the major clades of cladobranch sea slugs (Mollusca, Gastropoda, Heterobranchia), but fails to resolve the position of the enigmatic genus *Embletonia*. BMC Ecol Evol. 2021;21:226.

125. Goodheart JA, Bazinet AL, Valdés Á, Collins AG, Cummings MP. Prey preference follows phylogeny: evolutionary dietary patterns within the marine gastropod group Cladobranchia (Gastropoda: Heterobranchia: Nudibranchia). BMC Evol Biol. 2017;17:221.

126. Goodheart JA, Bazinet AL, Collins AG, Cummings MP. Relationships within Cladobranchia (Gastropoda: Nudibranchia) based on RNA-Seq data: an initial investigation. R Soc Open Sci. 2015;2:150196.

127. Laetsch DR, Blaxter ML. KinFin: Software for Taxon-Aware Analysis of Clustered Protein Sequences. G3. 2017;7:3349–57.

128. Putri GH, Anders S, Pyl PT, Pimanda JE, Zanini F. Analysing high-throughput sequencing data in Python with HTSeq 2.0. arXiv [q-bio.GN]. 2021.

129. Love MI, Huber W, Anders S. Moderated estimation of fold change and dispersion for RNA-seq data with DESeq2. Genome Biol. 2014;15:550.

130. Kuehn E, Clausen DS, Null RW, Metzger BM, Willis AD, Özpolat BD. Segment number threshold determines juvenile onset of germline cluster expansion in *Platynereis dumerilii*. J Exp Zool B Mol Dev Evol. 2022;338:225–40.

131. Schindelin J, Arganda-Carreras I, Frise E, Kaynig V, Longair M, Pietzsch T, et al. Fiji: an open-source platform for biological-image analysis. Nat Methods. 2012;9:676–82.

132. Preibisch S, Saalfeld S, Tomancak P. Globally optimal stitching of tiled 3D microscopic image acquisitions. Bioinformatics. 2009;25:1463–5.

133. Towns J, Cockerill T, Dahan M, Foster I, Gaither K, Grimshaw A, et al. XSEDE: Accelerating Scientific Discovery. Computing in Science & Engineering. 2014;16:62–74.

