## supplements version 2 for "A chromosome-level genome for the nudibranch gastropod *Berghia stephanieae* helps parse clade-specific gene expression in novel and conserved phenotypes": Additional_File_1.pdf

### HiRise Scaffolding Report

April 25, 2021
Nudibranch
Deirdre Lyons
Scripps Institution of Oceanography

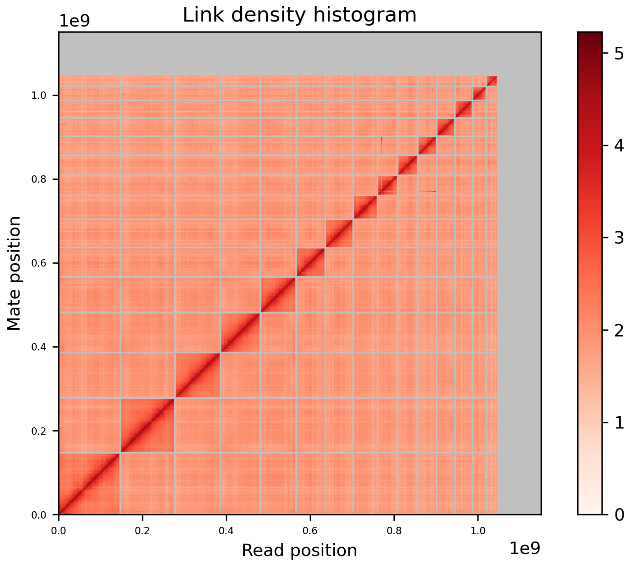


#### Contents

- Overview
- Contiguity Metrics
- BUSCO
- Pair Size Distribution
- Scaffolding Summary
- Materials & Methods

#### Overview

| Assembly | Total Length (bp) | N50 | L50 | N90 | L90 |
|---|---|---|---|---|---|
| Input Assembly | 1,150,334,909 | 6,921,793 | 46 | 232,553 | 294 |
| Dovetail HiRise Assembly | 1,150,404,709 | 68,992,636 | 6 | 26,817,905 | 15 |

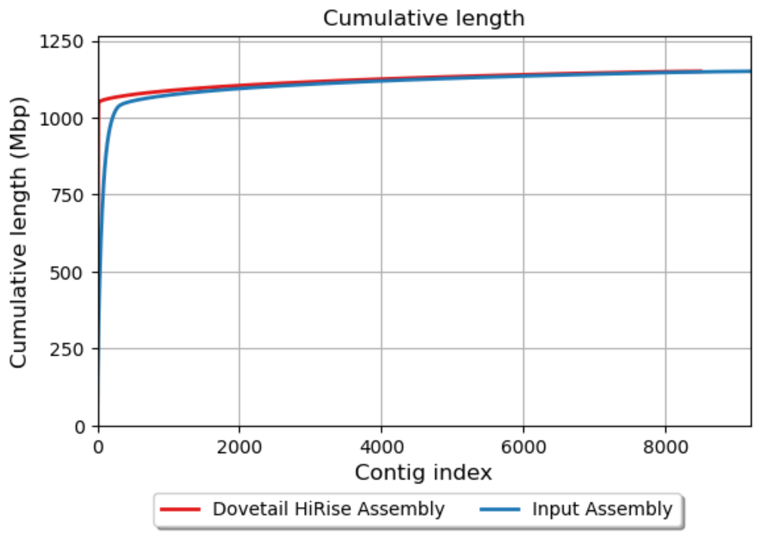

### Contiguity Metrics

| | Input Assembly | Dovetail HiRise Assembly |
|---|---|---|
| Largest scaffold | 49,550,825 | 146,869,960 |
| Number of scaffolds | 9,251 | 8,555 |
| Number of scaffolds > 1kbp | 9,189 | 8,493 |
| Number of gaps | 5,159 | 5,857 |
| Number of N's per 100 kbp | 44.85 | 50.91 |

### BUSCO

| Assembly | Complete BUSCOs (C) | Complete and single-copy BUSCOs (S) | Complete and duplicated BUSCOs (D) | Fragmented BUSCOs (F) | Missing BUSCOs (M) | Total BUSCO groups searched |
|---|---|---|---|---|---|---|
| Input Assembly | 231 (90.59%) | 227 (89.02%) | 4 | 12 | 12 | 255 |
| Dovetail HiRise Assembly | 229 (89.80%) | 225 (88.24%) | 4 | 10 | 16 | 255 |

- BUSCO version is: 4.0.5
- The lineage dataset is: eukaryota_odb10 (Creation date: 2020-09-10, number of species: 70, number of BUSCOs: 255)

### Pair Size Distribution

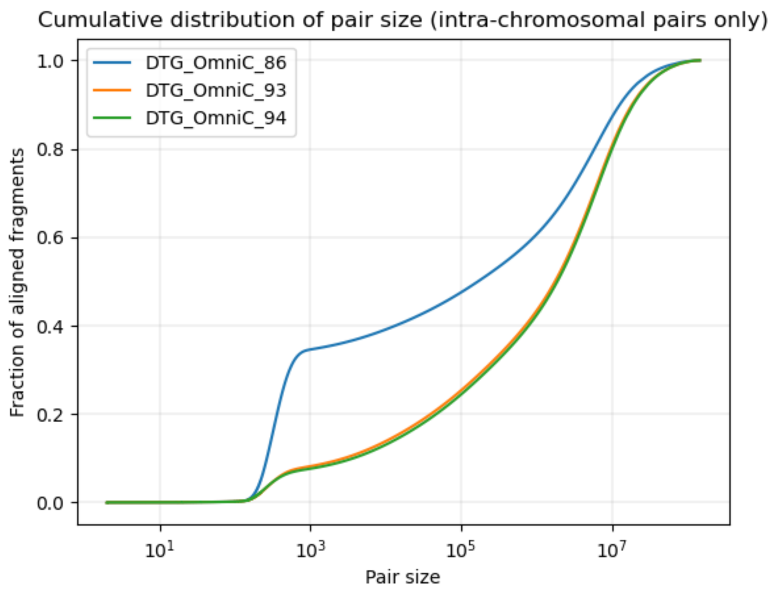

### HiRise Scaffolding Information

| | |
|---|---|
| Number of joins made by HiRise | 698 |
| Number of breaks made to input assembly by HiRise | 2 |
| Read-pairs | 0 |

### Materials and Methods

#### Dovetail Omni-C Library Preparation and Sequencing

For each Dovetail Omni-C library, chromatin was fixed in place with formaldehyde in the nucleus and then extracted. Fixed chromatin was digested with DNAse I, chromatin ends were repaired and ligated to a biotinylated bridge adapter followed by proximity ligation of adapter containing ends. After proximity ligation, crosslinks were reversed and the DNA purified. Purified DNA was treated to remove biotin that was not internal to ligated fragments. Sequencing libraries were generated using NEBNext Ultra enzymes and Illumina-compatible adapters. Biotin-containing fragments were isolated using streptavidin beads before PCR enrichment of each library. The library was sequenced on an Illumina HiSeqX platform to produce a approximately 30x sequence coverage. Then HiRise used MQ>50 reads for scaffolding (see "read-pair" above for figures).

#### Scaffolding the Assembly with HiRise

The input de novo assembly and Dovetail OmniC library reads were used as input data for HiRise, a software pipeline designed specifically for using proximity ligation data to scaffold genome assemblies (Putnam et al, 2016). Dovetail OmniC library sequences were aligned to the draft input assembly using bwa (https://github.com/lh3/bwa). The separations of Dovetail OmniC read pairs mapped within draft scaffolds were analyzed by HiRise to produce a likelihood model for genomic distance between read pairs, and the model was used to identify and break putative misjoins, to score prospective joins, and make joins above a threshold.

#### References

1. Putnam NH, O'Connell BL, Stites JC, Rice BJ, Blanchette M, Calef R, Troll CJ, Fields A, Hartley PD, Sugnet CW, Haussler D, Rokhsar DS, Green RE. Genome Research. 2016 Mar;26(3):342-50.
