## supplements version 2 for "A chromosome-level genome for the nudibranch gastropod *Berghia stephanieae* helps parse clade-specific gene expression in novel and conserved phenotypes": Additional_File_2_Figures_S1toS17.docx

### GenomeScope Profile - Figure S1

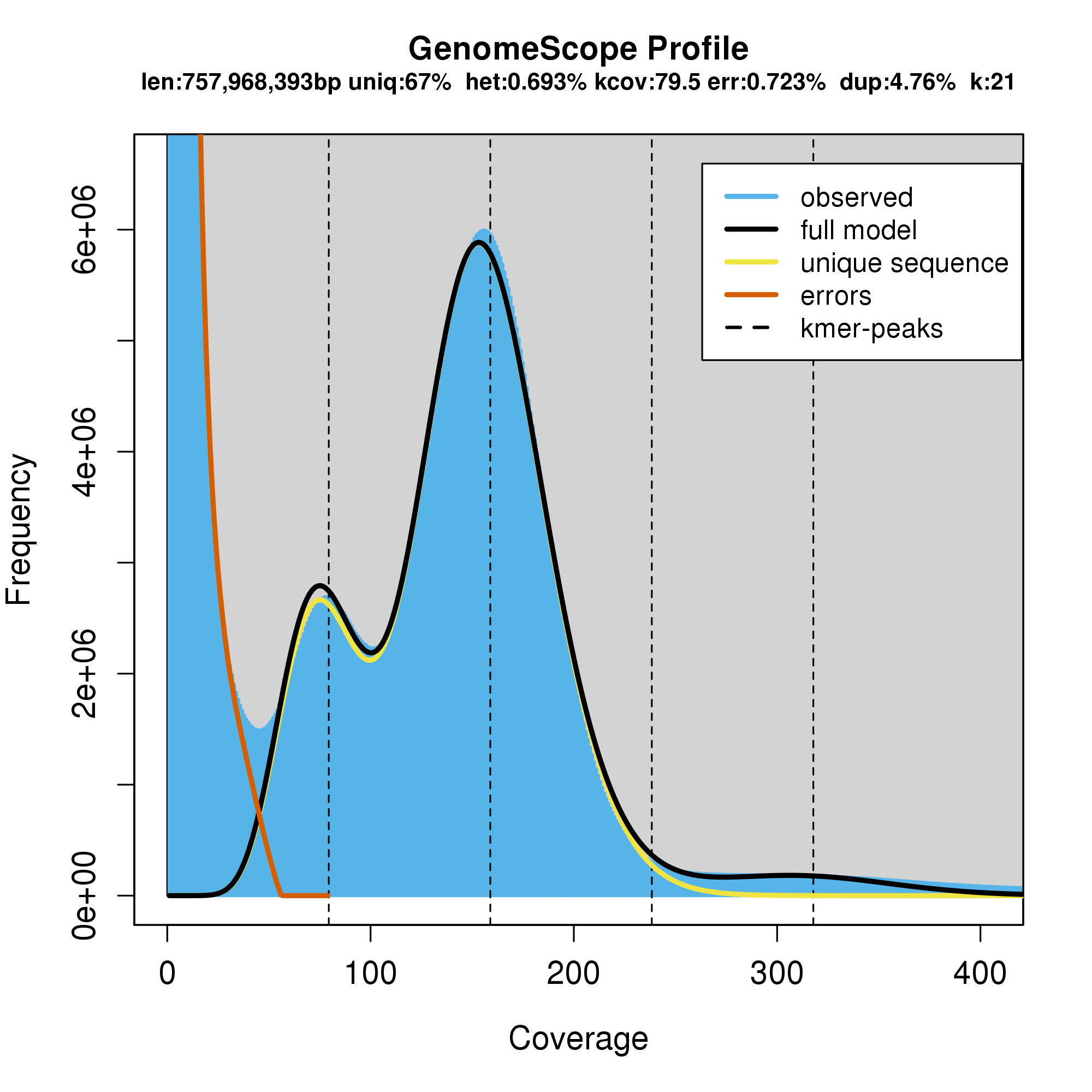

**Figure S1.** K-mer spectra and fitted models for a GenomeScope analysis of our Dovetail Omni-C data for *Berghia stephanieae*.

### Genome Summary Statistics - Figure S2

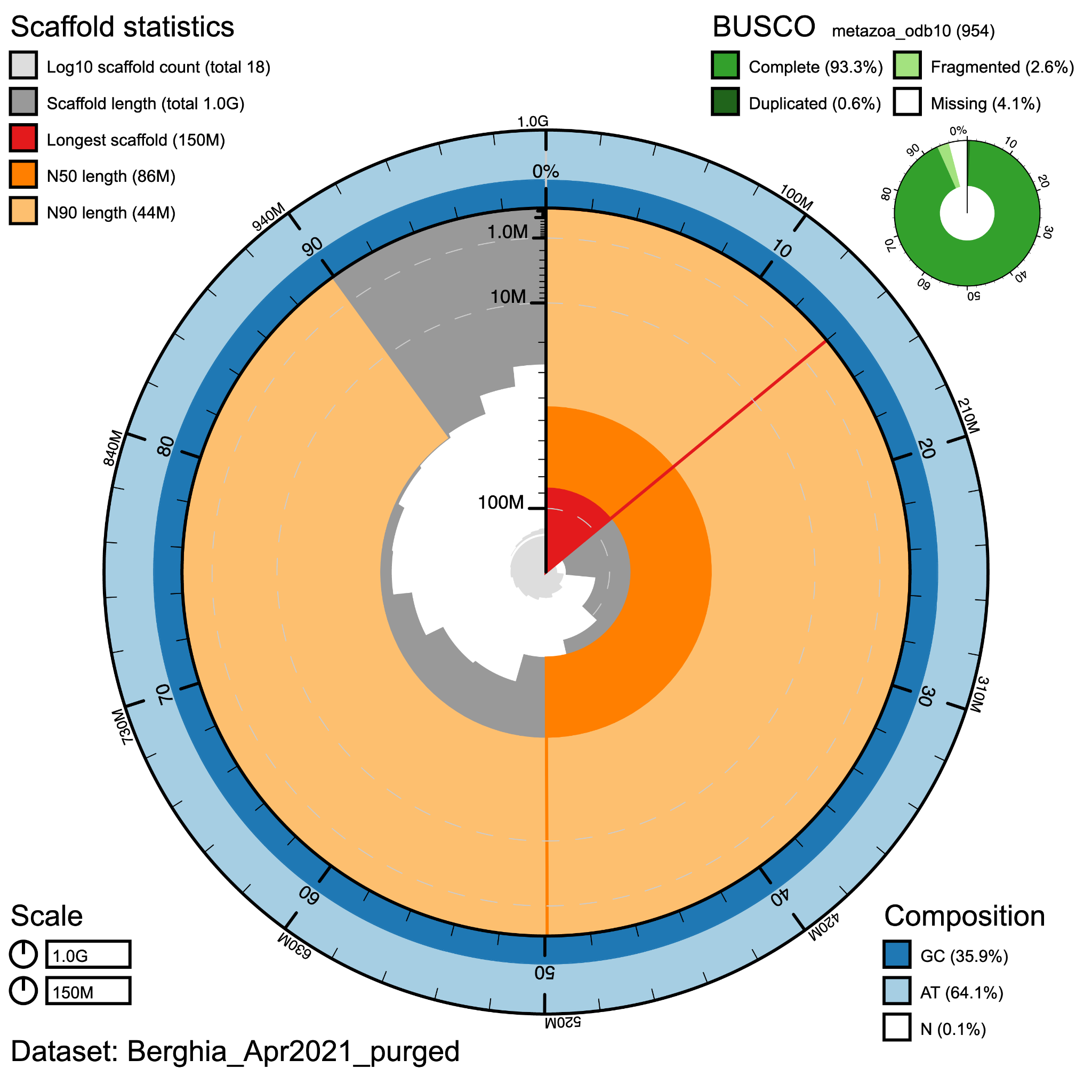

**Figure S2.** Snail plot showing high contiguity of the filtered genome (18 scaffolds), where 99% of the genome is found in the top 15 scaffolds. This plot also indicates the high levels of completeness (93.3%) when compared to metazoan single-copy ortholog datasets (BUSCO), with low levels of duplication (0.6%).

### OrthoFinder Summary Statistics - Figures S3-S4

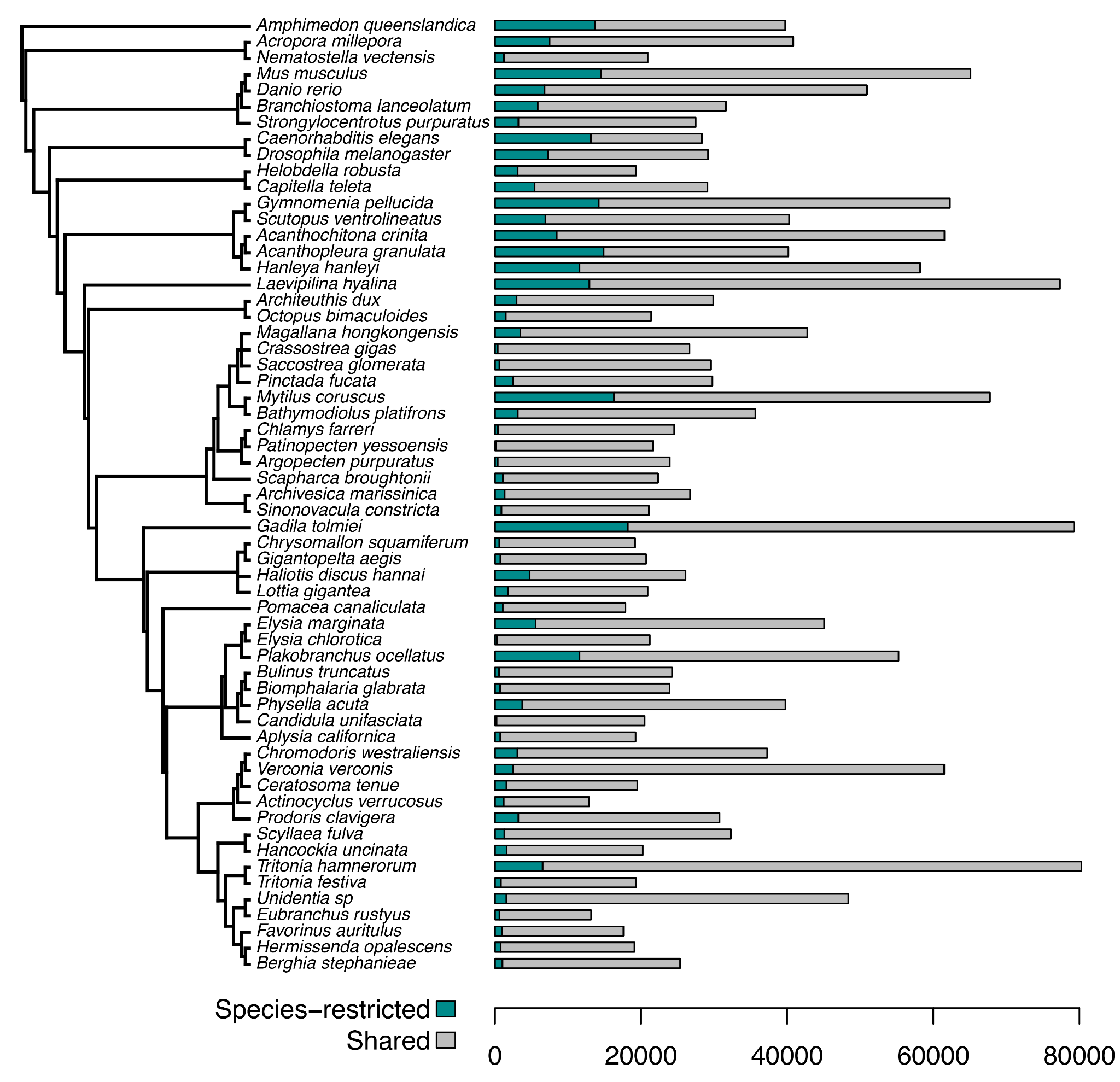

**Figure S3.** This chart shows the proteins in each proteome that were classified as species-specific based on the OrthoFinder and KinFin analyses.

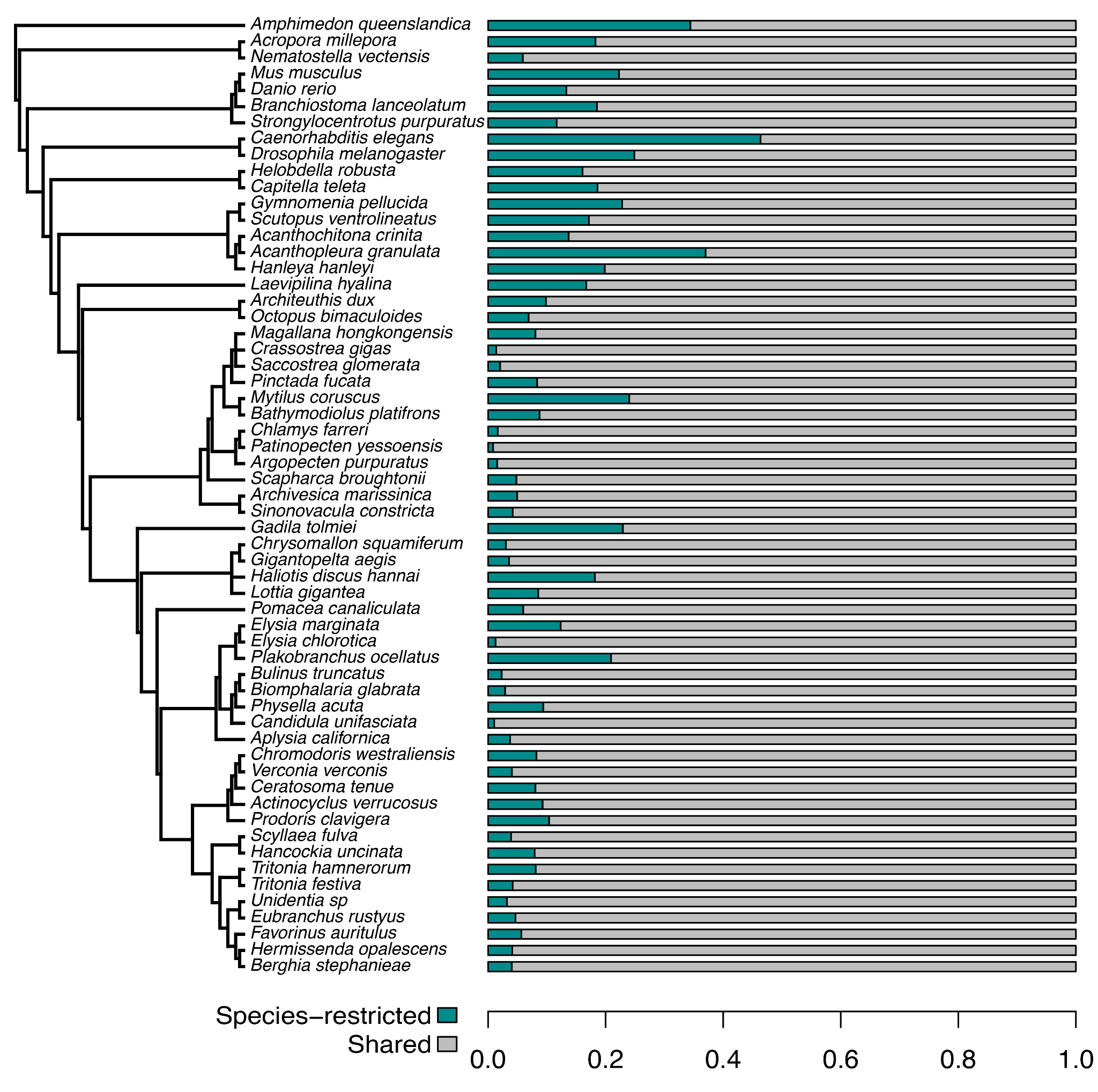

**Figure S4.** This chart shows the proportion of proteins in each proteome that were classified as species-specific based on the OrthoFinder and KinFin analyses.

### Rarefaction Curves - Figures S5-S9

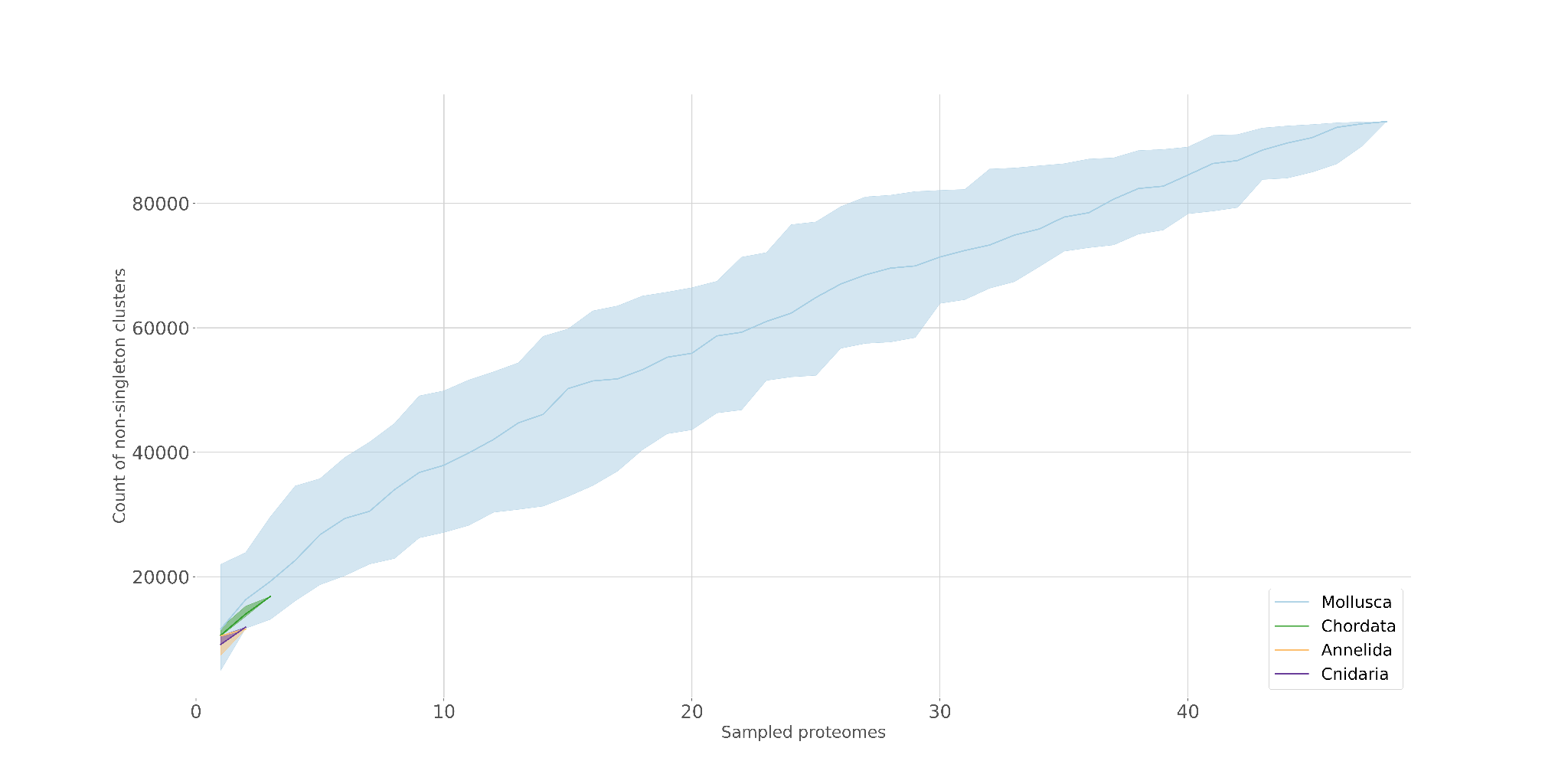

**Figure S5.** Rarefaction curve for gene sampling at the phylum level. This graph indicates that gene sampling for Mollusca (blue) is reaching an asymptote, which suggests that our sampling of these genes is sufficient for identifying *Berghia* genes that would match to other Mollusca proteomes.

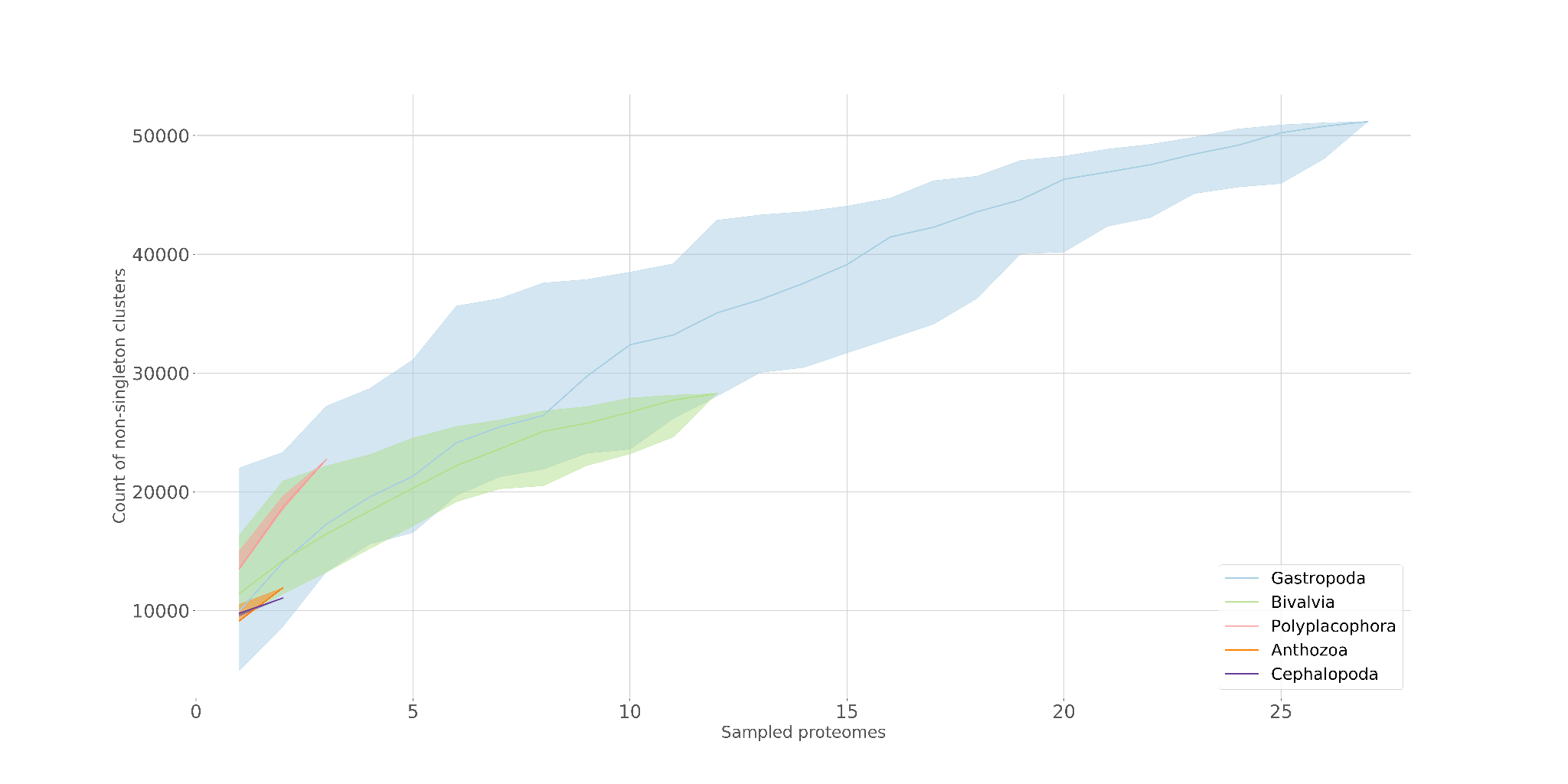

**Figure S6.** Rarefaction curve for gene sampling at the class level. This graph indicates that gene sampling for Gastropoda (blue) and Bivalvia (green) are reaching an asymptote, which suggests that our sampling of these genes is sufficient for identifying *Berghia* genes that would match to other gastropod proteomes.

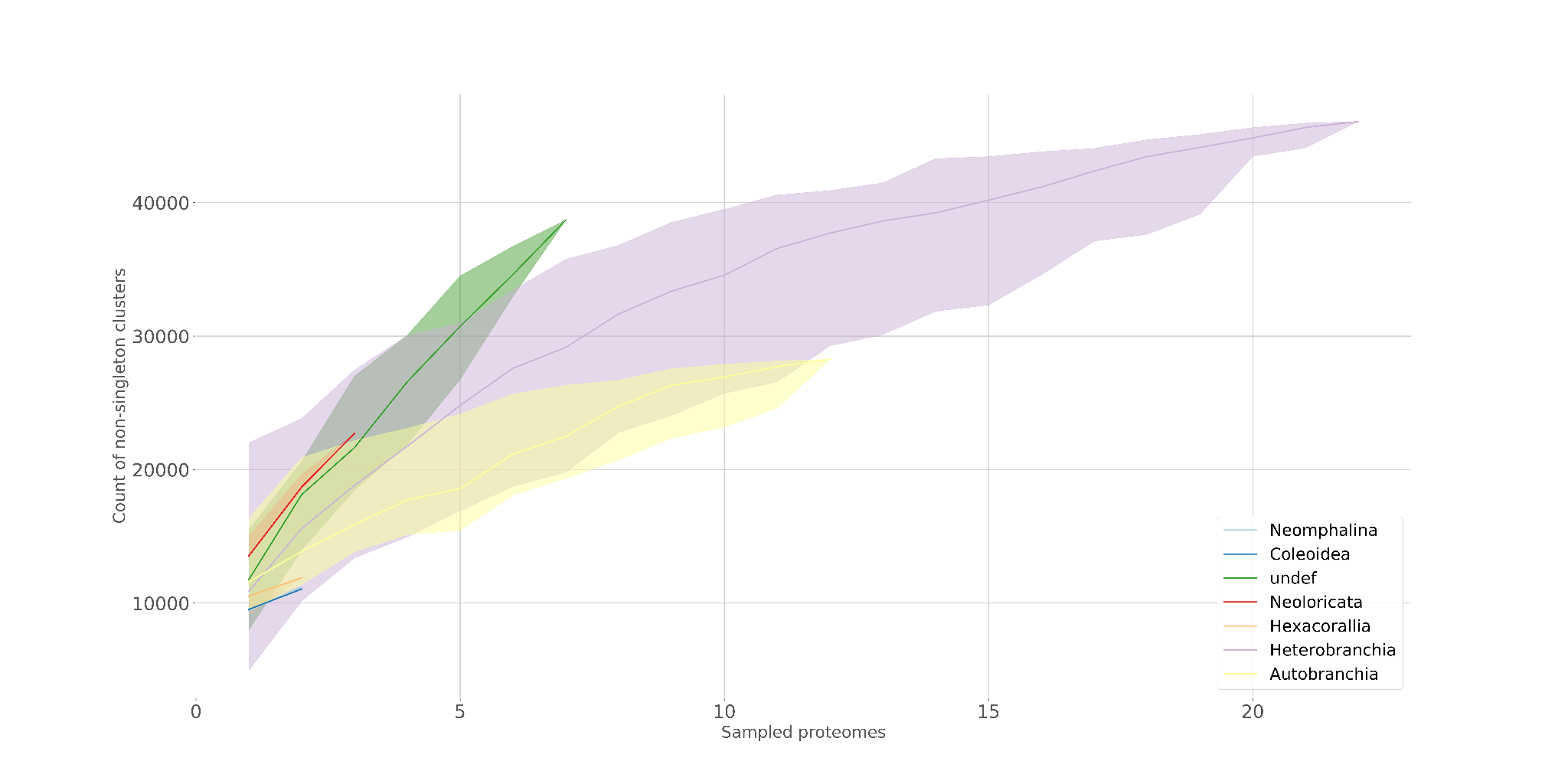

**Figure S7.** Rarefaction curve for gene sampling at the subclass level. This graph indicates that gene sampling for Heterobranchia (purple) are reaching an asymptote, which suggests that our sampling of these genes is sufficient for identifying *Berghia* genes that would match to other gastropod proteomes.

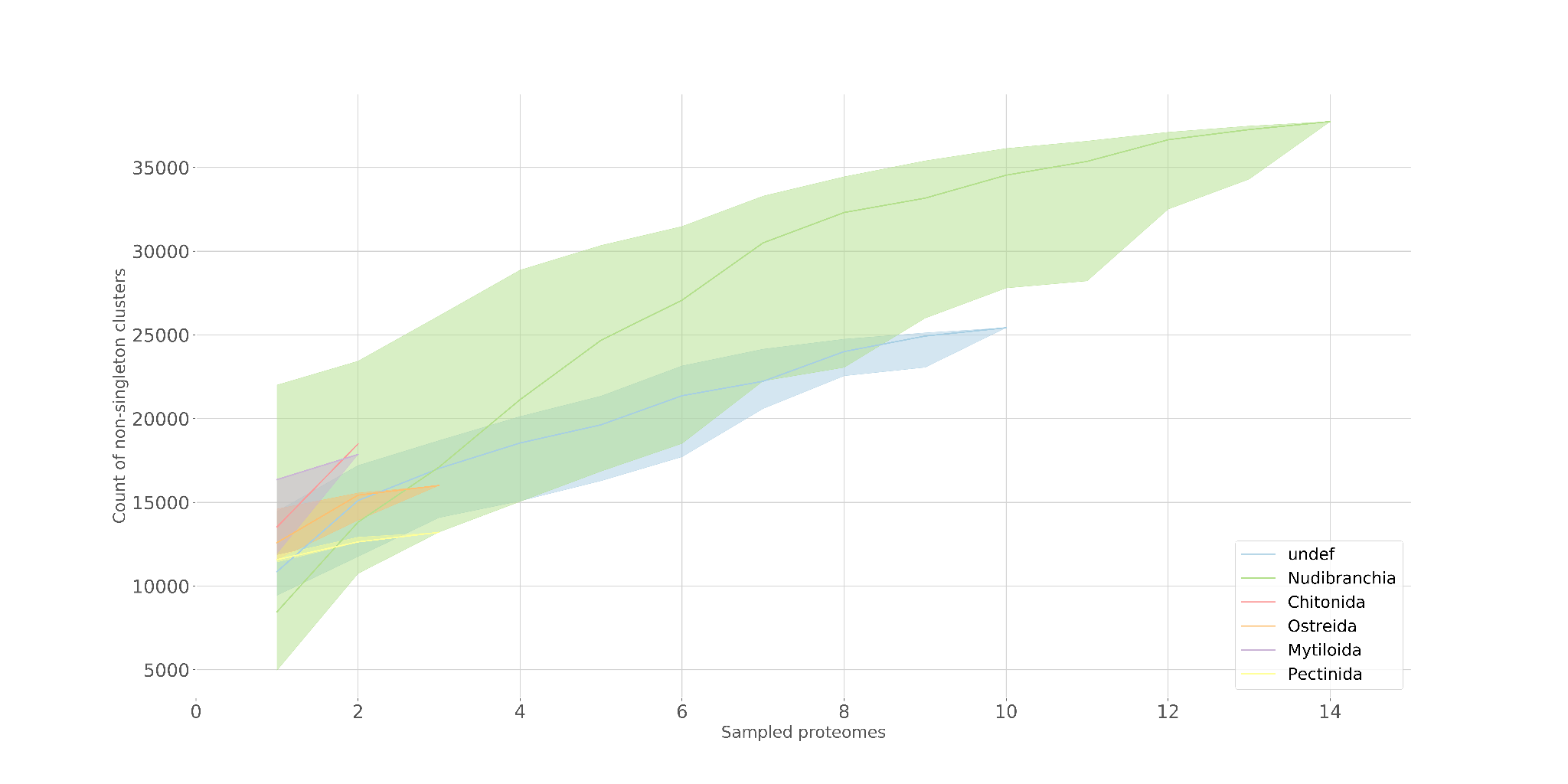

**Figure S8.** Rarefaction curve for gene sampling at the order level. This graph indicates that gene sampling for Nudibranchia (green) is reaching an asymptote, which suggests that our sampling of these genes is sufficient for identifying *Berghia* genes that would match to other nudibranch proteomes.

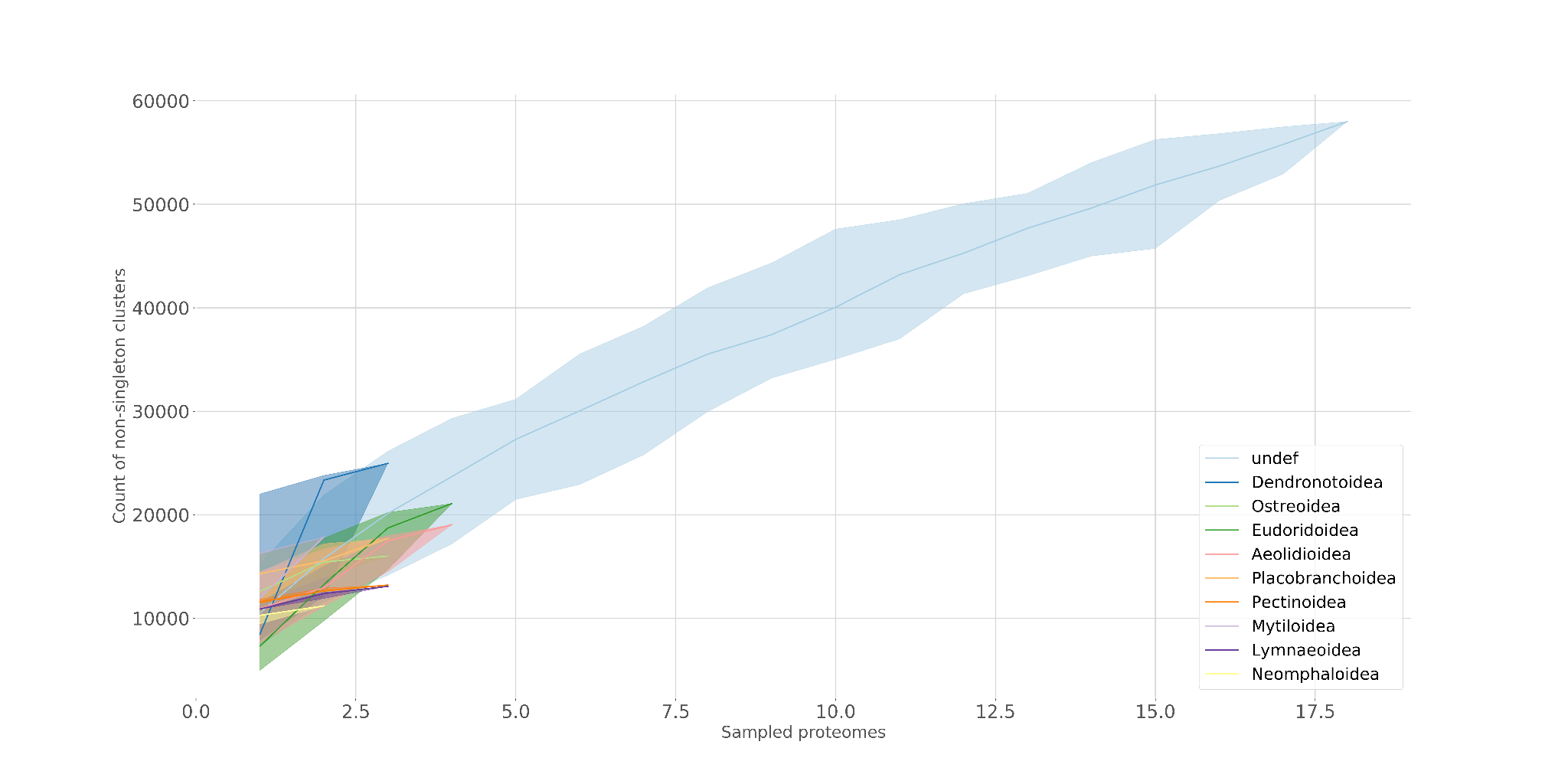

**Figure S9.** Rarefaction curve for gene sampling at the superfamily level. This graph indicates that gene sampling for Aeolidina (blue) is perhaps reaching an asymptote, though this group is not as well sampled as other levels. This still suggests that our sampling of these genes is sufficient for identifying *Berghia* genes that would match to other aeolid proteomes.

### KinFin Results - Figures S10-S11

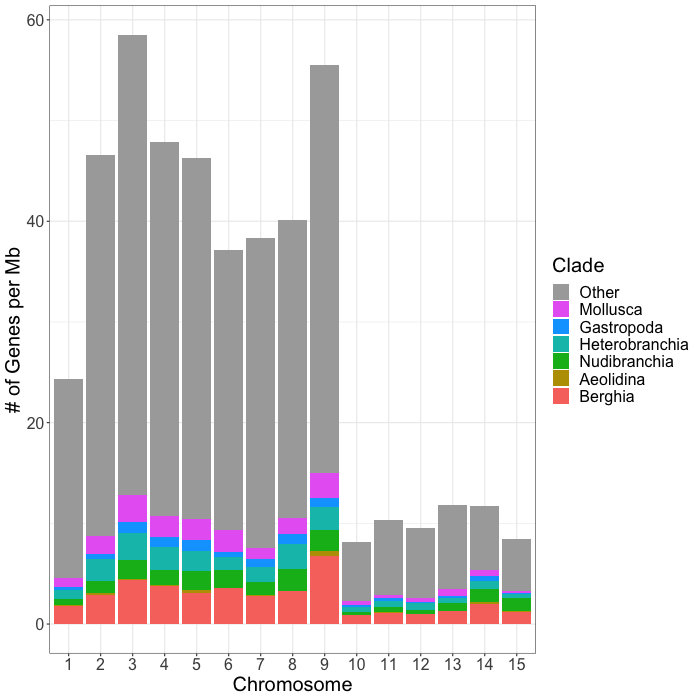

**Figure S10.** Distribution of both clade-specific and non-clade-specific (Other) genes across the putative chromosomes in our *Berghia stephanieae* genome. To the left is the raw gene counts per chromosome, and to the right is the number of genes per Mb of sequence to account for differences in chromosome lengths.
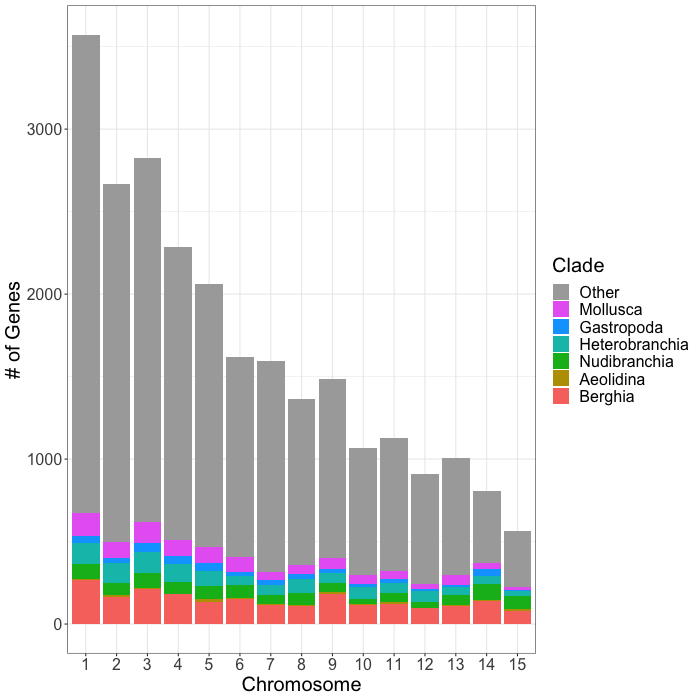

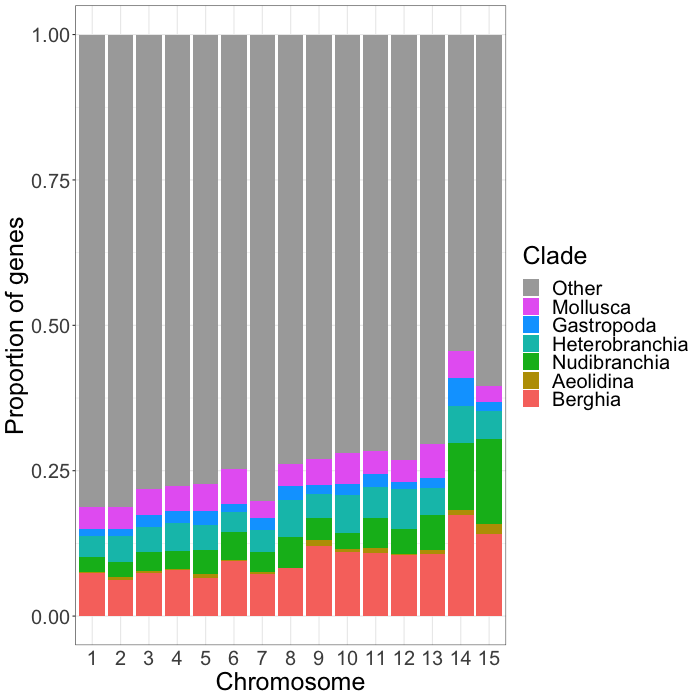

**Figure S11.** This chart shows the proportion of genes on each chromosome that are clade-specific and non-clade-specific (Other).

### Differential Expression Results - Figures S12-S18

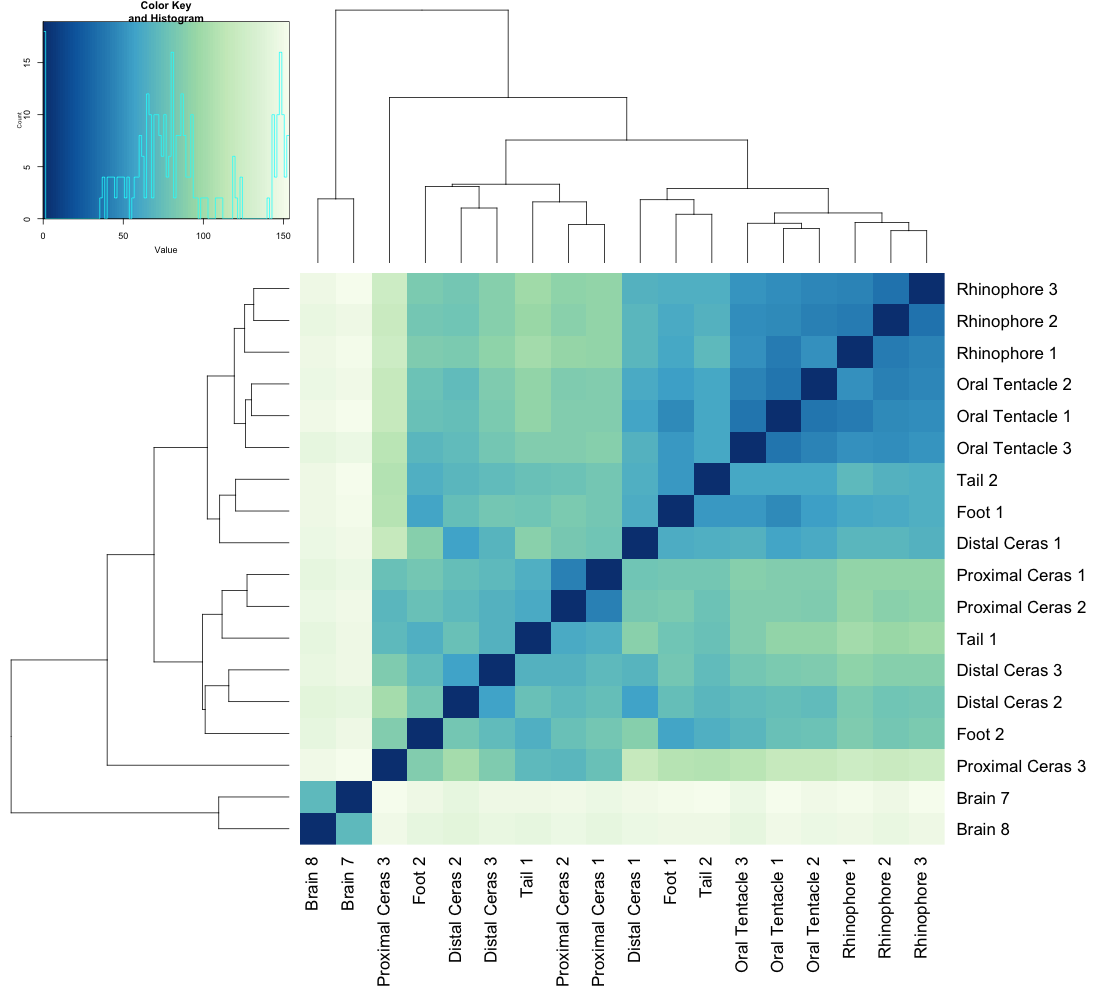

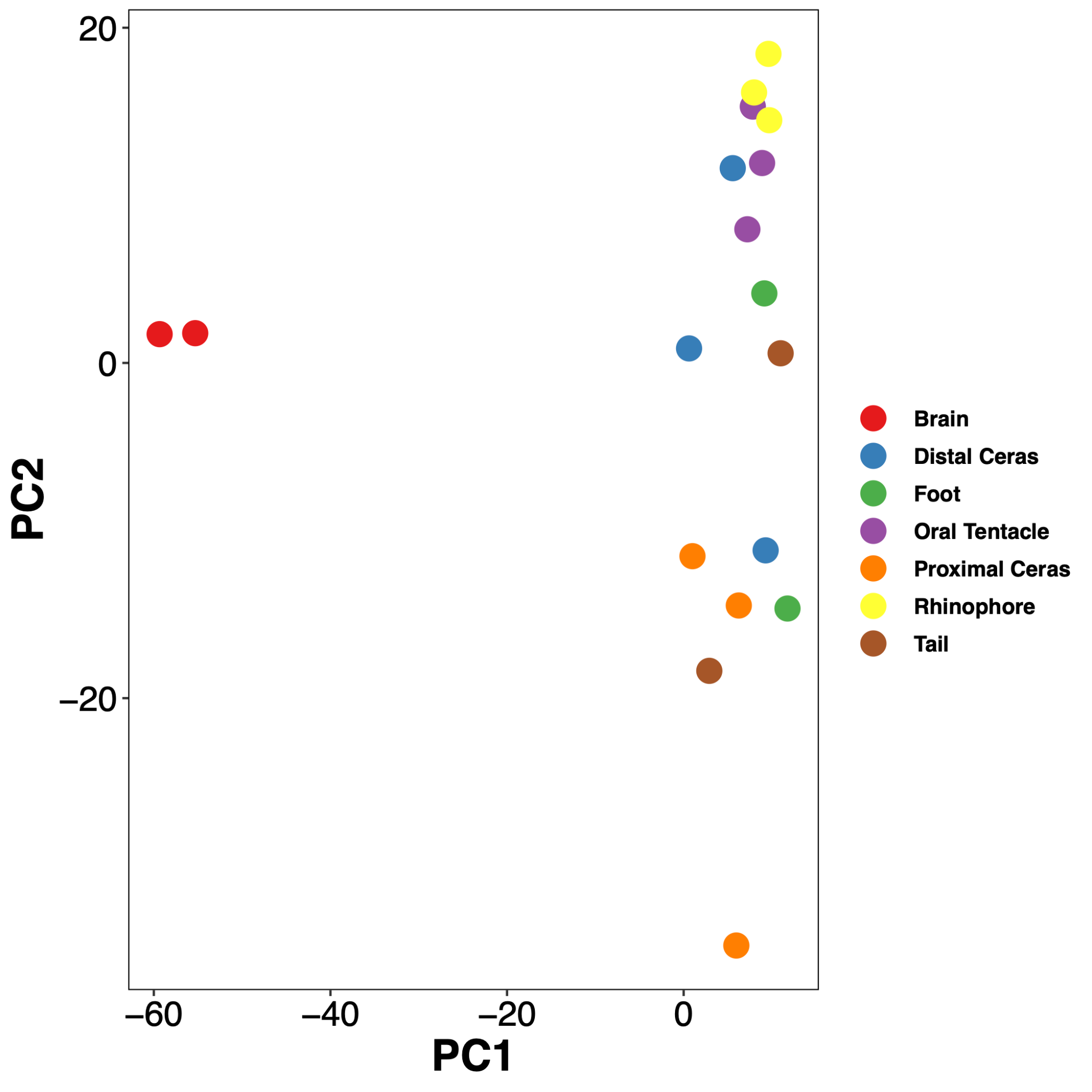

**Figure S12.** Gene expression across tissues in genes classified as *Berghia*-specific**.** Left: Heatmap of expression profiles; Right: PCA-plot showing similarity of expression within and among tissues.

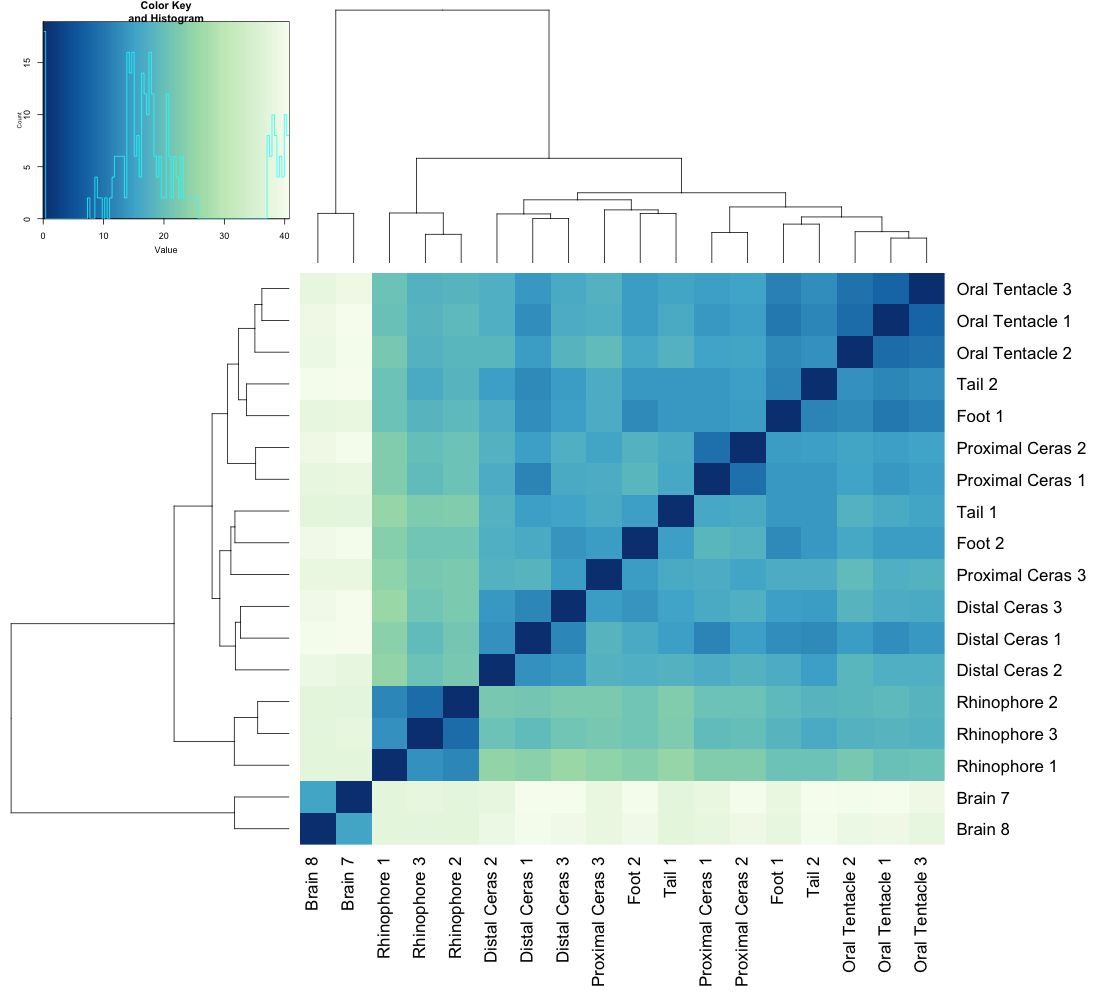

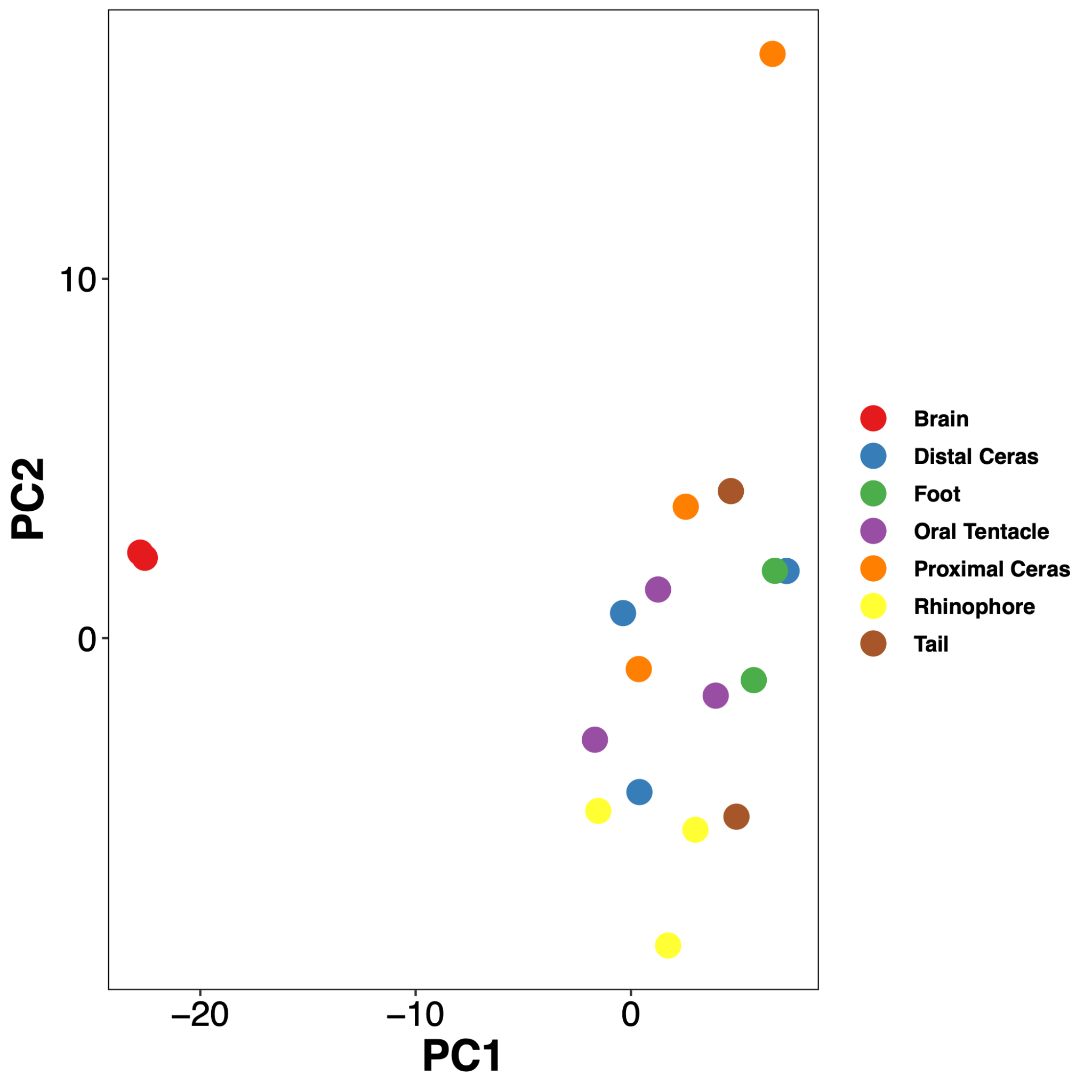

**Figure S13.** Gene expression across tissues in genes classified as Aeolidina-specific**.** Left: Heatmap of expression profiles; Right: PCA-plot showing similarity of expression within and among tissues.

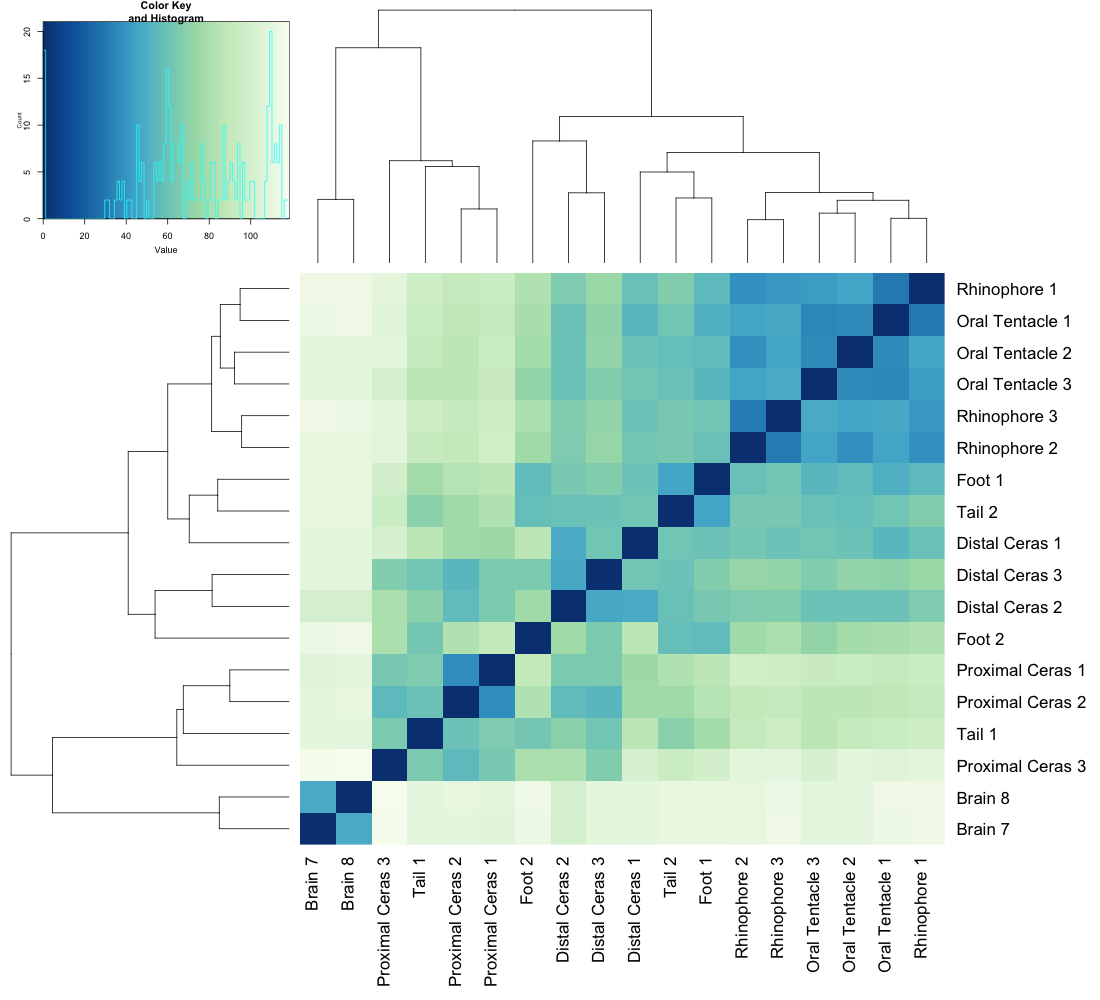

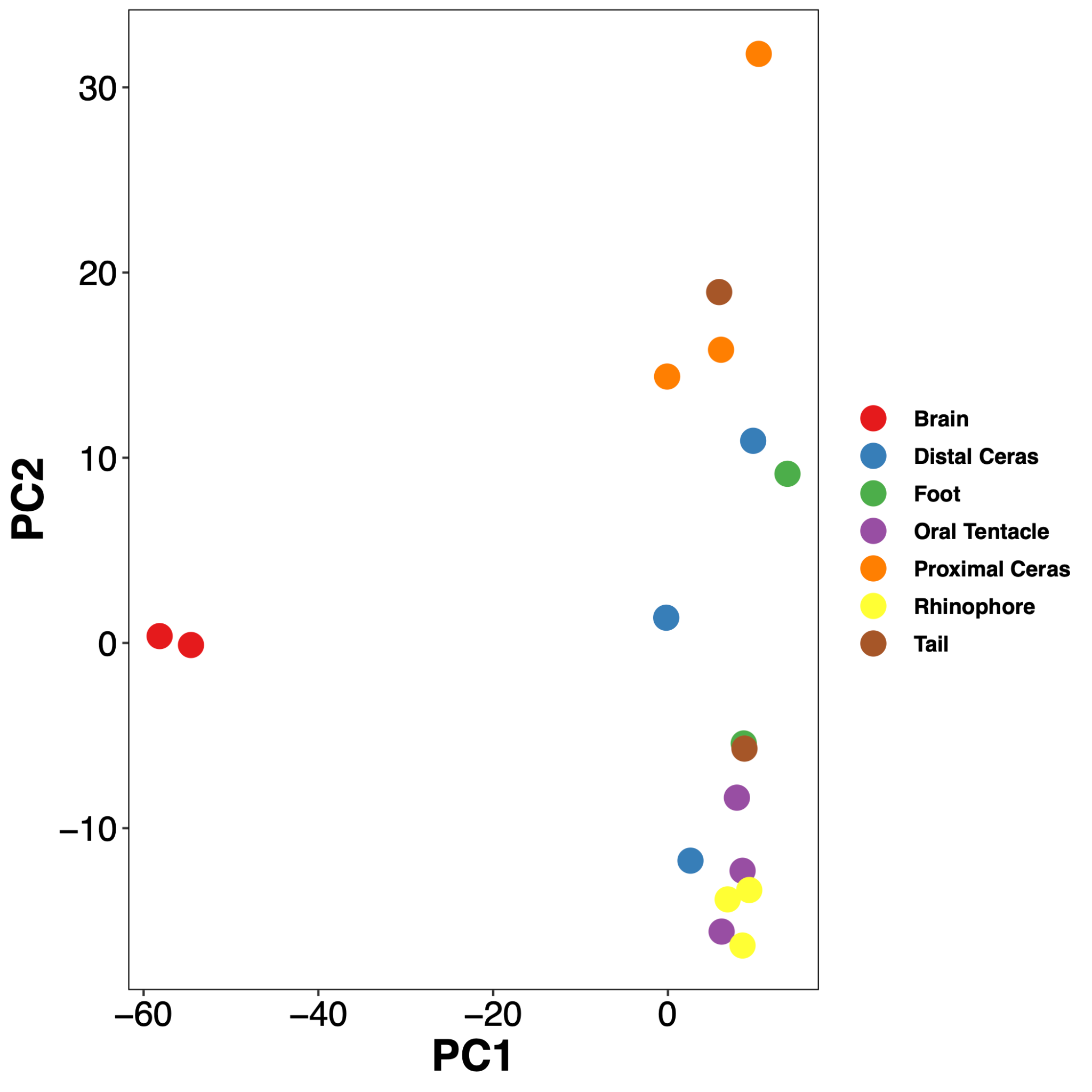

**Figure S14.** Heatmap of gene expression across tissues in genes classified as Nudibranchia-specific (left) and a PCA-plot showing similarity of expression within and among tissues (right).

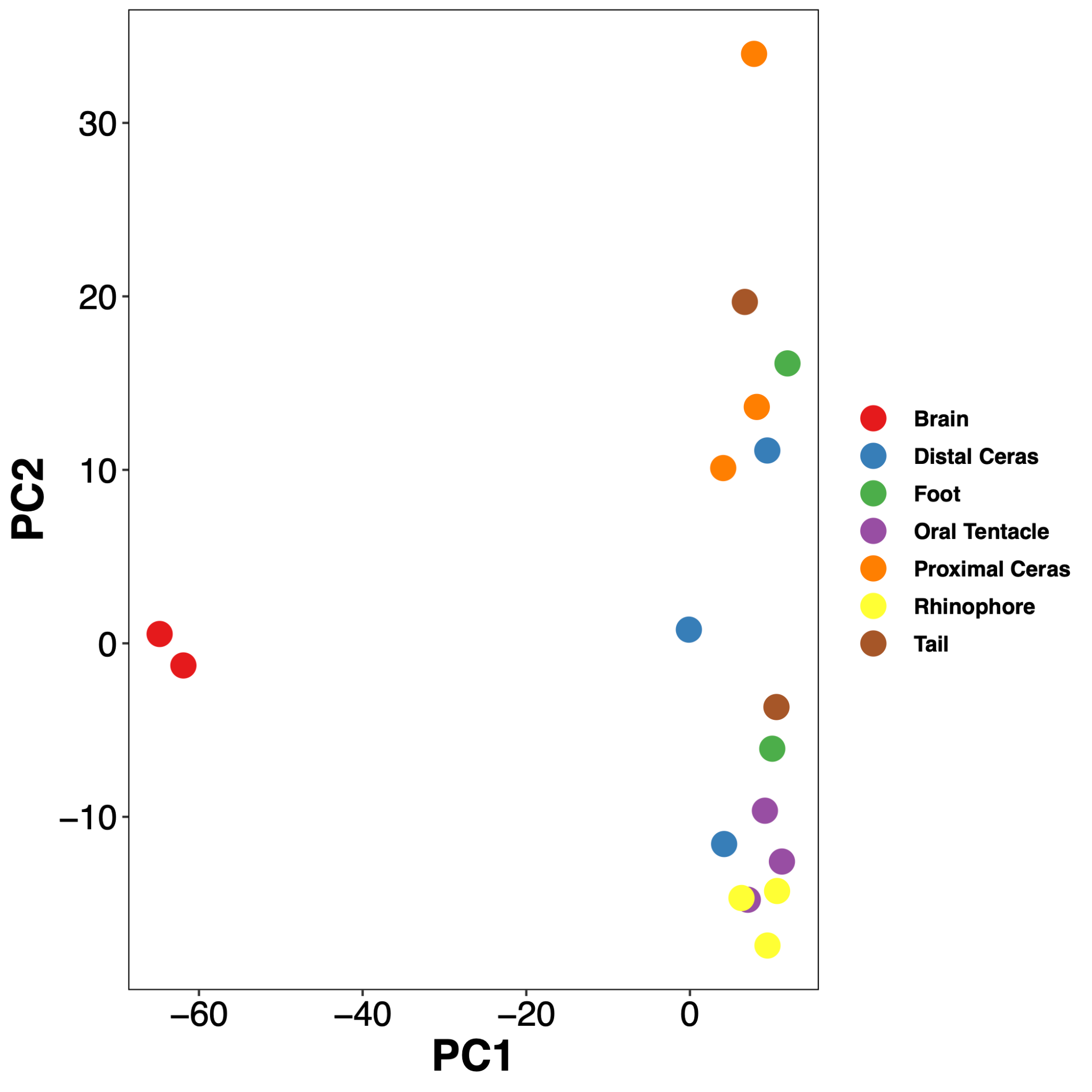

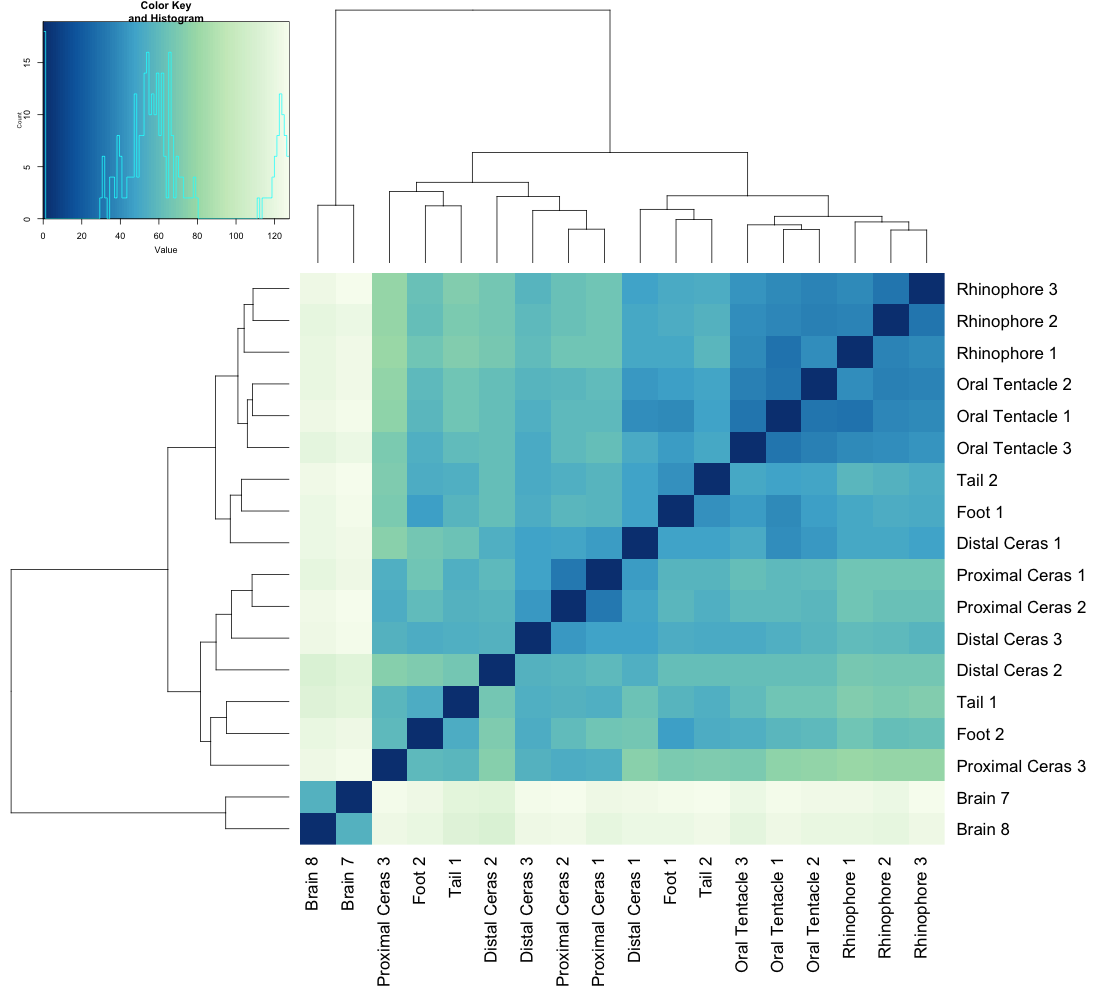

**Figure S15.** Gene expression across tissues in genes classified as Heterobranchia-specific**.** Left: Heatmap of expression profiles; Right: PCA-plot showing similarity of expression within and among tissues.

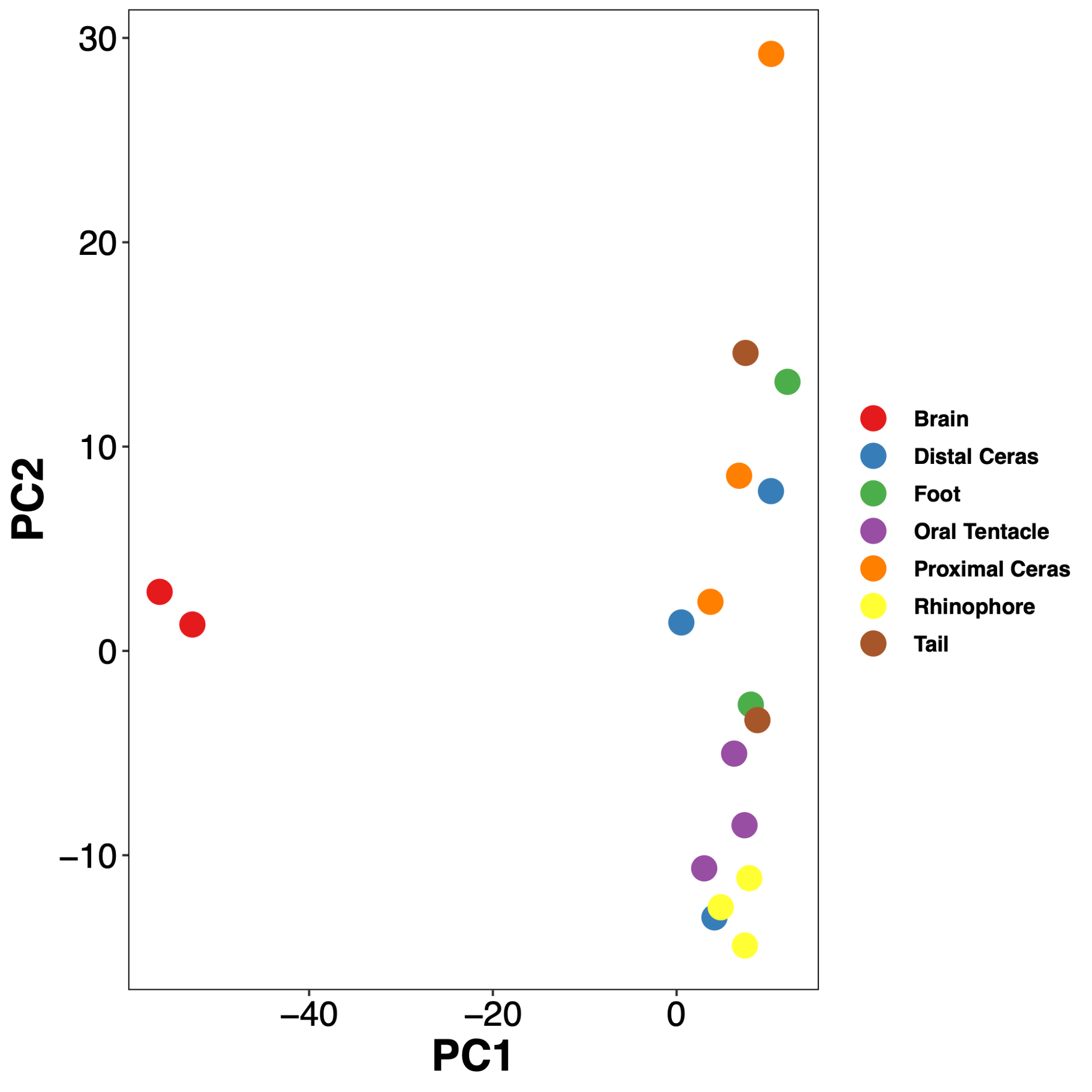

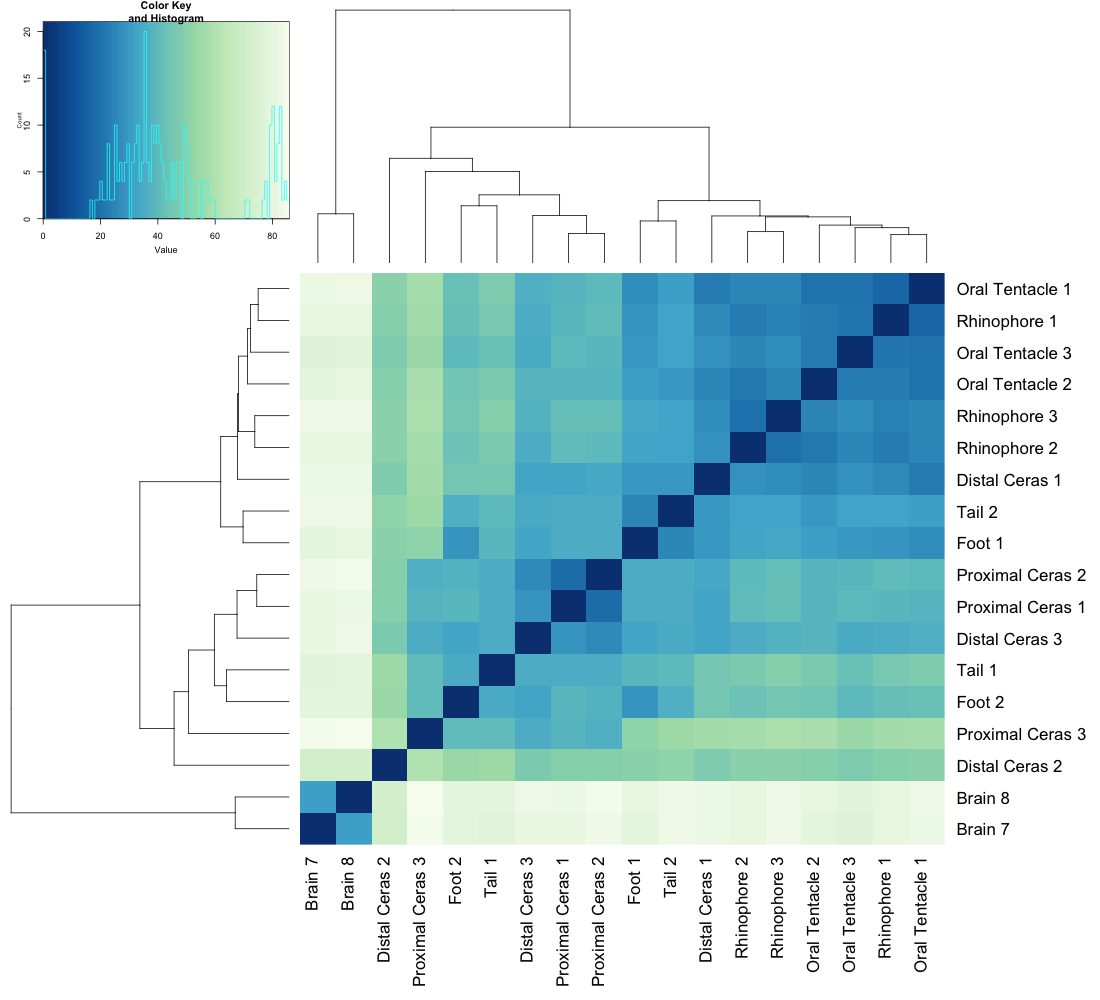

**Figure S16.** Gene expression across tissues in genes classified as Gastropoda-specific**.** Left: Heatmap of expression profiles; Right: PCA-plot showing similarity of expression within and among tissues.

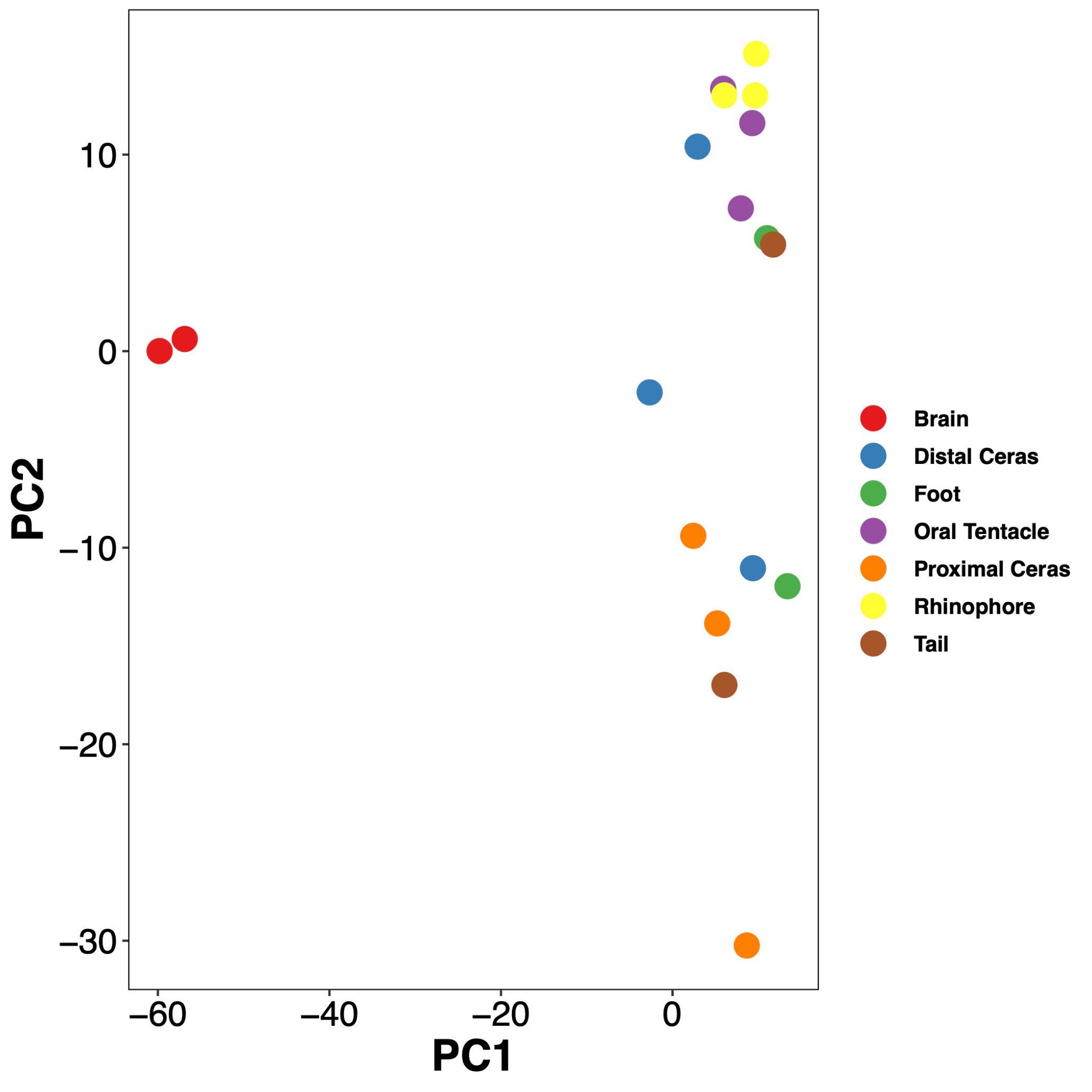

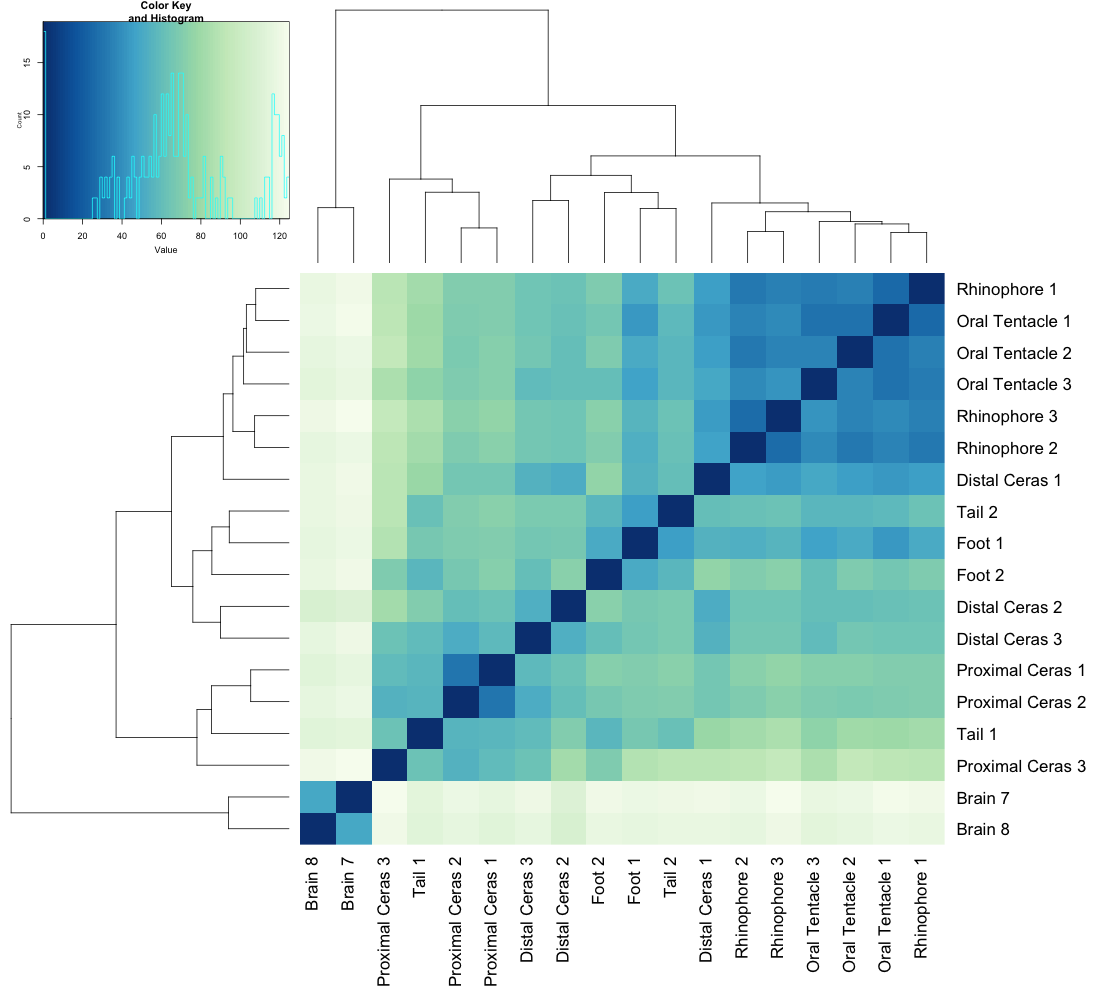

**Figure S17.** Gene expression across tissues in genes classified as Mollusca-specific**.** Left: Heatmap of expression profiles; Right: PCA-plot showing similarity of expression within and among tissues.

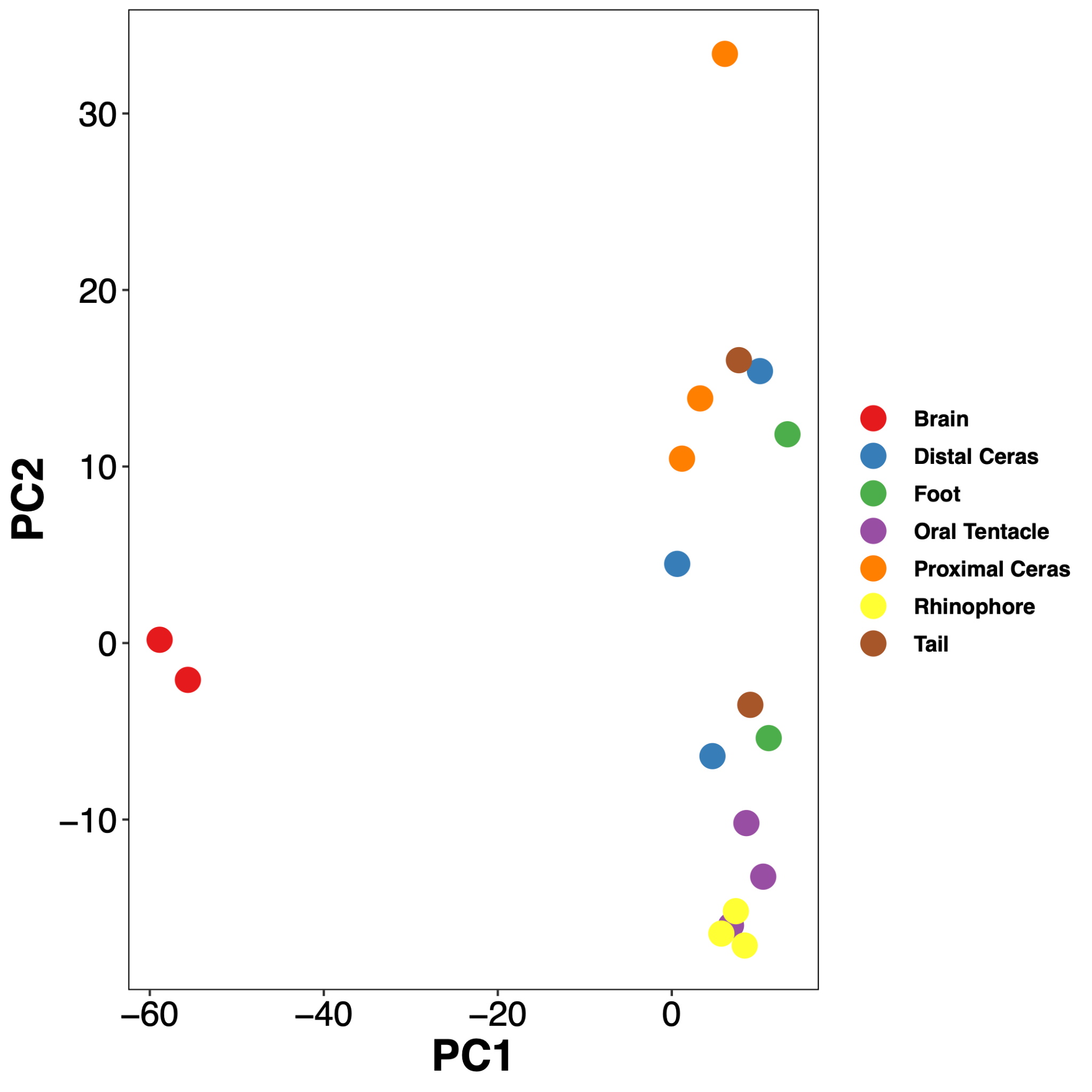

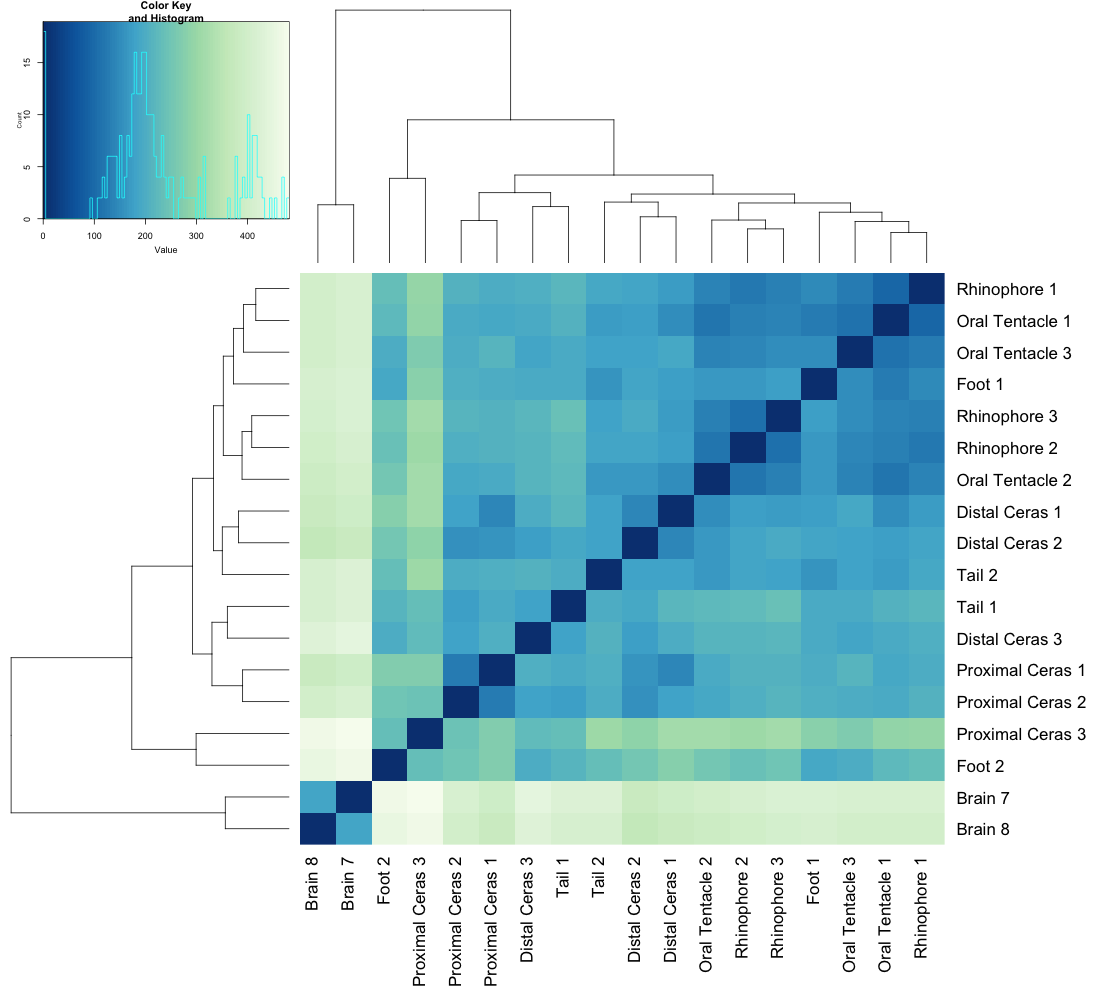

**Figure S18.** Gene expression across tissues in genes classified as “Other”**.** Left: Heatmap of expression profiles; Right: PCA-plot showing similarity of expression within and among tissues.
