## supplements version 2 for "A chromosome-level genome for the nudibranch gastropod *Berghia stephanieae* helps parse clade-specific gene expression in novel and conserved phenotypes": Additional_File_12_Commands_Used_Annotation.docx

Berghia genome annotation

### Genome Filtering

#### Run Purgedups

Input(s): Berghia_Apr2021_hirise.fasta (assembly file from Dovetail)

Output(s): Berghia_Apr2021_hirise_purged.fasta

*Minimap2 version 2.18-r1015*

*Purge_dups version 1.2.5*

Split fasta file into smaller sequences:

split_fa Berghia_Apr2021_hirise.fasta > Berghia_Apr2021_hirise.fasta.split

Map split sequences to themselves:

minimap2 -xasm5 -DP Berghia_Apr2021_hirise.fasta.split Berghia_Apr2021_hirise.fasta.split -t 8 | gzip -c - > Berghia_Apr2021_hirise.fasta.split.self.paf.gz

Remove duplicates:

purge_dups -2 -T cutoffs -c PB.base.cov Berghia_Apr2021_hirise.fasta.split.self.paf.gz > dups.bed 2> purge_dups.log

get_seqs -e dups.bed Berghia_Apr2021_hirise.fasta

mv purged.fa Berghia_Apr2021_hirise_purged.fasta

#### Run Blobtoolkit analyses

Input(s): Berghia_Apr2021_hirise_purged.fasta, XDOVE_2019_pacbioreads_merged.fastq.gz,

Output(s): Berghia_Apr2021_purged blobtools directory

*blobtools version 2.3.3*

*BUSCO version 5.1.2*

*Blastn version 2.11.0+*

*Minimap2 version 2.18-r1015*

Create blobtools directory for Berghia:

blobtools create --fasta Berghia_Apr2021_hirise_purged.fasta --meta Berghia_Apr2021_purged.yaml --taxid 1287507 --taxdump taxdump Berghia_Apr2021_purged

BUSCO - Metazoa odb10:

busco -m genome -i Berghia_Apr2021_hirise_purged -l metazoa_odb10 -o berghia_genome_Apr2021_purged_busco.v4.0.5 -c 20

Coverage

minimap2 -ax sr -t 16 Berghia_Apr2021_hirise_purged.fasta XDOVE_2019_pacbioreads_merged.fastq.gz | samtools sort -@16 -O BAM -o Berghia_Apr2021_reallypurged_pacbiomerged.bam

BLASTn runs

export BLASTDB=$BLASTDB:/ocean/projects/bio210009p/shared/data/databases/nt

blastn -db nt -query Berghia_Apr2021_hirise_purged.fasta -outfmt "6 qseqid staxids bitscore std" -max_target_seqs 10 -max_hsps 1 -evalue 1e-25 -num_threads 30 -out Berghia_Apr2021_purged.ncbi.blastn.out

Add coverage, busco, and blast hits:

```
blobtools add \
      	--hits Berghia_Apr2021_reallypurged.ncbi.blastn.out\
      	--hits Berghia_Apr2021_hirise_reallypurged.fasta.diamond.blastx.out \
      	--taxrule bestsumorder \
      	--taxdump /ocean/projects/bio210009p/shared/data/databases/taxdump \
      	--busco berghia_genome_Apr2021_purged_busco.v4.0.5/run_metazoa_odb10/full_table.tsv
      	--cov Berghia_Apr2021_reallypurged_pacbiomerged.bam \
      	Berghia_Apr2021_purged
--threads 8
```

#### Filter genome

Input(s): Berghia_Apr2021_purged blobtools directory

Output(s): Berghia_Apr2021_hirise_purged.filtered.fasta

Filtered genome by length and bacteria:

blobtools filter --param length--Min=200000 --param bestsumorder_phylum--Keys=no-hit,Bacteria-undef,Tenericutes --fasta Berghia_Apr2021_hirise_purged.fasta --summary STDOUT Berghia_Apr2021_purged

RECOMMENDED - Simplify scaffold names:

NOTE: The main reason for this is that BRAKER2 (Augustus specifically I think) has issues with scaffold names that have symbols in them or are too long. I fixed this much later in the process (which is why this file is not used in the next few steps), but this command would have removed the extra (and unnecessary) symbols.

sed 's/;HRSCAF=[0-9]*_[0-9]*//g' Berghia_Apr2021_hirise_purged.filtered.fasta | sed 's/_//g' > Berghia_Apr2021_hirise_purged.filtered.edited.fasta

### Repeat Masking

Input(s): Berghia_Apr2021_hirise_purged.filtered.fasta, Bsteph_ref.fa.no_tes.fa (reference proteome for Berghia from transcriptome assembly)

Output(s): Repeatmasked genome (*.softmasked, *.hardmasked)

#### Make Repeat Library (RepeatModeler)

*RepeatModeler Version 2.0.1*

*BLAST 2.10.0+*

[*https://blaxter-lab-documentation.readthedocs.io/en/latest/filter-repeatmodeler-library.html*](https://blaxter-lab-documentation.readthedocs.io/en/latest/filter-repeatmodeler-library.html)

Compile initial repeat library (with Classifications):

RepeatModeler -database berghia -pa 2 -LTRStruct

Blast proteome against RepeatMasker TE database:

blastp -query ../Berghia_alltissues_onerep_trinity291_transdecoder_cdhit95_noaliens_fulltranscripts_novectors_nocontaminants.fasta.transdecoder.pep -db /ocean/projects/bio210009p/shared/tools/miniconda3/envs/repeatmodeler/share/RepeatMasker/Libraries/RepeatPeps.lib -outfmt '6 qseqid staxids bitscore std sscinames sskingdoms stitle' -max_target_seqs 25 -culling_limit 2 -num_threads 8 -evalue 1e-5 -out Bsteph_ref.pep.vs.RepeatPeps.25cul2.1e5.blastp.out

Remove TEs from proteome:

fastaqual_select.pl -f ../Berghia_alltissues_onerep_trinity291_transdecoder_cdhit95_noaliens_fulltranscripts_novectors_nocontaminants.fasta.transdecoder.cds -e <(awk '{print $1}' Bsteph_re..pep.vs.RepeatPeps.25cul2.1e5.blastp.out | sort | uniq) > Bsteph_ref.fa.no_tes.fa

Blast proteome against RepeatModeler library:

makeblastdb -in Bsteph_ref.fa.no_tes.fa -dbtype nucl

blastn -task megablast -query consensi.fa.classified -db Bsteph_ref.fa.no_tes.fa -outfmt '6 qseqid staxids bitscore std sscinames sskingdoms stitle' -max_target_seqs 25 -culling_limit 2 -num_threads 8 -evalue 1e-10 -out repeatmodeller_lib.v.Bsteph_ref.fa.no_tes.25cul2.1e10.megablast.out

Remove hits from RepeatModeler library:

fastaqual_select.pl -f consensi.fa.classified -e <(awk '{print $1}' repeatmodeller_lib.v.Bsteph_ref.fa.no_tes.25cul2.1e25.megablast.out | sort | uniq) > consensi.fa.classified.filtered_for_CDS_repeats.fa

#### Repeat Mask Genome (RepeatModeler)

*RepeatMasker version 4.1.2-p1*

Hardmask for STAR aligner:

RepeatMasker Berghia_Apr2021_hirise_purged.filtered.fasta -e ncbi -lib RM_24730.MonJul121509252021/consensi.fa.classified.filtered_for_CDS_repeats.fa -pa 20

mv Berghia_Apr2021_hirise_purged.filtered.fasta.masked Berghia_Apr2021_hirise_purged.filtered.fasta.hardmasked

mv Berghia_Apr2021_hirise_purged.filtered.fasta.out Berghia_Apr2021_hirise_purged.filtered.fasta_hard.out

mv Berghia_Apr2021_hirise_purged.filtered.fasta.cat Berghia_Apr2021_hirise_purged.filtered.fasta_hard.cat

mv Berghia_Apr2021_hirise_purged.filtered.fasta.tbl Berghia_Apr2021_hirise_purged.filtered.fasta_hard.tbl

Softmask for Augustus/BRAKER:

RepeatMasker Berghia_Apr2021_hirise_purged.filtered.fasta -e ncbi -lib RM_24730.MonJul121509252021/consensi.fa.classified.filtered_for_CDS_repeats.fa -xsmall -pa 20

mv Berghia_Apr2021_hirise_purged.filtered.fasta.masked Berghia_Apr2021_hirise_purged.filtered.fasta.softmasked

mv Berghia_Apr2021_hirise_purged.filtered.fasta.out Berghia_Apr2021_hirise_purged.filtered.fasta_soft.out

mv Berghia_Apr2021_hirise_purged.filtered.fasta.cat Berghia_Apr2021_hirise_purged.filtered.fasta_soft.cat

mv Berghia_Apr2021_hirise_purged.filtered.fasta.tbl Berghia_Apr2021_hirise_purged.filtered.fasta_soft.tbl

### Alignment of RNA-seq reads (Illumina)

Input(s): Berghia_Apr2021_hirise_purged.filtered.fasta.hardmasked, combined Illumina short reads from all stages and tissues listed in paper

Output(s): Berghia_Apr2021_masked_starmappedAligned_sort.bam and Berghia_Apr2021_masked_starmappedAligned_sort.bam.bai

*STAR version 2.7.9a*

*HiSat2 version 2.2.1*

#### Align RNA-seq data

##### STAR

STAR runThreadN 8 --runMode genomeGenerate --genomeSAindexNbases 13 --genomeDir star_index_repeatmodeler_masked --genomeFastaFiles Berghia_Apr2021_hirise_purged.filtered.fasta.hardmasked --twopassMode --outSAMtype BAM SortedByCoordinate

STAR --runThreadN 8 --genomeDir star_index_repeatmodeler_masked --readFilesIn Berghia_alltissues_allstages_withbrain_R1.fq.gz Berghia_alltissues_allstages_withbrain_R2.fq.gz --outFileNamePrefix Berghia_Apr2021_repeatmodeler_masked_starmapped --readFilesCommand zcat

samtools view -bS Berghia_Apr2021_repeatmodeler_masked_starmappedAligned.out.sam > Berghia_Apr2021_repeatmodeler_masked_starmappedAligned.out.bam

samtools sort Berghia_Apr2021_repeatmodeler_masked_starmappedAligned.out.bam -o Berghia_Apr2021_repeatmodeler_masked_starmappedAligned_sort.bam

samtools index Berghia_Apr2021_repeatmodeler_masked_starmappedAligned_sort.bam

##### HiSat2

hisat2-build -p 8 Berghia_Apr2021_hirise_purged.filtered.fasta.hardmasked hisat2_index_repeatmodeler_masked

hisat2 -p 8 -x hisat2_index_repeatmodeler_masked -1 Berghia_alltissues_allstages_withbrain_R1.fq.gz -2 Berghia_alltissues_allstages_withbrain_R2.fq.gz -S Berghia_Apr2021_repeatmodeler_masked_hisat2mapped.sam

samtools view -bS Berghia_Apr2021_repeatmodeler_masked_starmappedAligned.out.sam > Berghia_Apr2021_repeatmodeler_masked_starmappedAligned.out.bam

samtools sort Berghia_Apr2021_repeatmodeler_masked_starmappedAligned.out.bam -o Berghia_Apr2021_repeatmodeler_masked_starmappedAligned_sort.bam

samtools index Berghia_Apr2021_repeatmodeler_masked_starmappedAligned_sort.bam

### Alignment of IsoSeq reads (PacBio)

Input(s): Berghia_Apr2021_hirise_purged.filtered.fasta.hardmasked, combined IsoSeq long reads from all stages and tissues listed in paper

Output(s): Berghia_Apr2021_masked_starmappedAligned_sort.bam and Berghia_Apr2021_masked_starmappedAligned_sort.bam.bai

*minimap2 version 2.18-r1015*

#### Align RNA-seq data

```
minimap2 -t 30 -ax splice -uf --secondary=no -C5 -O6,24 -B4 \
Berghia_Apr2021_hirise_purged.filtered.fasta \
lyons_alltissues_flnc.fasta \
     	> berghia_flnc_isoforms.sam \
     	2> berghia_flnc_isoforms.sam.log

samtools view -S -b berghia_flnc_isoforms.sam > berghia_flnc_isoforms.bam
samtools sort berghia_flnc_isoforms.bam -o berghia_flnc_isoforms.sorted.bam
samtools index berghia_flnc_isoforms.sorted.bam
```

### BRAKER2 Annotation

Input(s): Berghia_Apr2021_hirise_purged.filtered.fasta_edited.softmasked,

Output(s): Annotation gtf and fasta files for masked genome

*BRAKER2 version 2.1.6*

#### Initial Annotation

braker.pl --species=Bsteph --genome=Berghia_Apr2021_hirise_purged.filtered.fasta_edited.softmasked \

--prot_seq mollusca_odb10_with_berghiabuscos.fasta \

--bam=Berghia_Apr2021_repeatmodeler_masked_starmappedtwopassAligned.sortedByCoord.out.bam,Berghia_Apr2021_repeatmodeler_masked_hisat2mapped_sort_shortheaders.bam,berghia_flnc_isoforms_bestlocus.sorted.bam \

--etpmode --softmasking -cores 12 \

--useexisting --gff3

#### Filter Annotations

Input(s): BRAKER2 files augustus.hints.gtf and hintsfile.gff

Output(s): Filtered annotation files

*BRAKER2 version 2.1.6*

##### Filter sequences

From BRAKER/scripts/prediction_analysis:

python selectSupportedSubsets.py --anySupport augustus.anysupport.gtf --fullSupport augustus.fullsupport.gtf --noSupport augustus.nosupport.gtf augustus.hints.gtf hintsfile.gff

sh filter_seqs.sh

filter_seqs.sh:

```
#pull out the transcripts from the filtered gtf file
awk '{print $10}' augustus.anysupport.gtf > anysupport.genes

#filter so that you get a file containing unique names
cat anysupport.genes | uniq > anysupport.genes.uniq

#remove the " and ; from each name
sed -i 's/"//g' anysupport.genes.uniq
sed -i 's/;//g' anysupport.genes.uniq

#perl one liner that uses the id file you just created to pull out corresponding sequences in the fasta file produced by braker
perl -ne 'if(/^>(\S+)/){$c=$i{$1}}$c?print:chomp;$i{$_}=1 if @ARGV' anysupport.genes.uniq augustus.hints.aa > augustus.hints.anysupport.aa
```

##### BUSCO Assessment

```
busco -m proteins -i augustus.hints.aa -l metazoa_odb10 -o  berghia_genome_annotation_Sept2021_busco.proteins.v4.0.5 -c 10

busco -m proteins -i augustus.hints.aa -l mollusca_odb10 -o berghia_genome_annotation_Sept_busco.mollusca.proteins.v4.0.5 -c 10

busco -m proteins -i augustus.hints.anysupport.aa -l metazoa_odb10 -o berghia_genome_annotation_Sept2021_anysupport_busco.proteins.v4.0.5
 -c 10

busco -m proteins -i augustus.hints.anysupport.aa -l mollusca_odb10 -o berghia_genome_annotation_Sept2021_anysupport_busco.mollusca.proteins.v4.0.5 -c 10
```

### Functional Annotations

#### BLASTP

Input(s): augustus.hints.anysupport.aa and blast databases

Output(s): Blast output files

*Protein-Protein BLAST 2.11.0+*

*genomeGTFtools (https://github.com/wrf/genomeGTFtools)*

##### BLASTP runs

blastp -query augustus.hints.anysupport.aa -db uniprot_sprot.fasta -num_threads 8 -max_target_seqs 1 -outfmt 6 -evalue 1e-3 > blastp.uniprot-sprot.outfmt6

blastp -query augustus.hints.anysupport.aa -db refseq_protein -num_threads 8 -max_target_seqs 1 -outfmt 6 -evalue 1e-3 > blastp.refseq.outfmt6

##### BLASTP output to genome annotation

blast2genomegff.py -b blastp.uniprot-sprot.outfmt6 -d uniprot_sprot.fasta -g augustus.anysupport.gff3 -x -p blastp > berghia_sprot.genome.gff

##### Create BLASTP table with gene names for use in R

1. Downloaded blastp results to my computer
2. Pulled out UniProt and RefSeq IDs to search for metadata
   a. awk '{ print $2 }' blastp/blastp_augustus_RMiso.uniprot-sprot.outfmt6 | awk 'BEGIN { FS = "|" } ; { print $2 }' | sort | uniq > uniprot_ids_filtered.txt
   b. awk '{ print $2 }' blastp/blastp_augustus_RMiso.refseq.outfmt6 | sort | uniq > refseq_ids_filtered.txt
3. Used download_uniprot.pl and download_uniprot_refseq.pl to download uniprot hits for IDs where uniprot metadata exists
   a. perl download_uniprot.pl uniprot_ids_filtered.txt > uniprot_data.txt
   b. perl download_uniprot_refseq.pl refseq_ids_filtered.txt > uniprot_data_refseq.txt
4. Run first part of R-script Bs_genome_blasthits.R to get the list of RefSeq annotations that are missing
5. Split the nouniprothits output file for batch entrez downloads
   a. split -l 2000 Bs_protein-nouniprothits-info_unique.txt Bs_protein-nouniprothits-info_unique.
6. Upload files to Batch Entrez (https://www.ncbi.nlm.nih.gov/sites/batchentrez) and download then combine summary text files
   a. cat protein_result_[A-Z].txt > protein_result_allRefSeq.txt
7. Process batch entrez summary files to get a useable table for refseq results
   a. bash process_batch_entrez.sh
8. Run second part of Bs_genome_blasthits.R to get the full blast list
   a. **Output file:** Bs_protein-blasthit-ALL-info.txt

#### InterProScan

Input(s): augustus.hints.anysupport.aa

Output(s): Interproscan annotation files - *.gff3, *.json, *.xml, *.tsv

*InterProScan v.5.52-86.0*

*CDD-3.18,Coils-2.2.1,Gene3D-4.3.0,Hamap-2020_05,MobiDBLite-2.0,PANTHER-15.0,Pfam-33.1,Phobius-1.01,PIRSF-3.10,PIRSR-2021_02,PRINTS-42.0,SFLD-4,SignalP_EUK-4.1,SMART-7.1,SUPERFAMILY-1.75,TIGRFAM-15.0,TMHMM-2.0c*

*MAKER script - iprscan2gff3*

##### InterProScan Run

interproscan.sh -i augustus.hints.anysupport.aa -b berghia_RM_iso_anysupport_ipscan_2021_11 -goterms -dp -t p -appl CDD-3.18,Coils-2.2.1,Gene3D-4.3.0,Hamap-2020_05,MobiDBLite-2.0,PANTHER-15.0,Pfam-33.1,Phobius-1.01,PIRSF-3.10,PIRSR-2021_02,PRINTS-42.0,SFLD-4,SignalP_EUK-4.1,SMART-7.1,SUPERFAMILY-1.75,TIGRFAM-15.0,TMHMM-2.0c

##### InterProScan output to genome annotation (from MAKER scripts)

iprscan2gff3 berghia_RM_iso_anysupport_ipscan_2021_11.tsv braker_annotations.gff3 > braker_iprscan_annotations.gff
